## Supplementary material for "ARTseq-FISH reveals position-dependent differences in gene expression of micropatterned mESCs": Supplementary Information_2024-02-13.docx

**The Supplementary Information includes:**

Supplementary Notes 1: Experimental development of ARTseq-FISH

Supplementary Notes 2: Computational pipeline development of ARTseq-FISH

Supplementary Fig. 1-31

Supplementary Tables Caption

References

### Supplementary Notes 1 (SN1)

**Experimental development of ARTseq-FISH**

### 1. Barcode and probe design for ARTseq-FISH

ARTseq-FISH utilizes the colour barcode scheme inspired by previous work ^1^. We designed four different readout sequences for each target to scale up the multiplexing capacity. In order to combine the protocol of sequential RNA FISH with protein detection based on oligonucleotide-labelled antibodies (Supplementary Fig. 1a), the limitation of the length of oligonucleotides conjugated to antibodies ^2^ requires an amplification step. This step is also required in order to combine RNA and protein detection. Therefore, we adopted the rolling circle amplification (RCA) of padlock probes (PLPs) as previous reported ^3^. Combining the barcode scheme and the RNA RCA *in situ* hybridization, we introduced PLPs, bridge probes (BrPs) and readout probes (RoPs) in the detection (Supplementary Fig. 1b-d and Table 1). Protein targets are detected by antibodies conjugated with a unique 46 nucleotides (nt) single stranded (ss) DNA. This unique ssDNA contains a 10 nt poly A (polyadenylate) linker and a specific 36 nt sequence that is complementary to a target-specific PLP. Each PLP contains two arms (18 nt of 5' arm and 18 nt of 3' arm of PLP) complementary to the targeting region, one unique 28 nt sequence identical to the BrP and one common RCA primer binding region. Besides the 28 nt sequences identical to paired PLPs, BrPs are composed of four overhang sequences with 18 nt each corresponding to different RoPs (Supplementary Fig. 1b-d and Table 1).

**Table 1: The component of oligonucleotides, padlock probes, bridge probe and readout probes.**

| **Oligonucleotides (46 nt)** | 5'- azide-AAAAAAAAAA- [36 nt unique sequence]-3' |
| --- | --- |
| **Padlock probe (84 nt)** | 5' -phosphorylated- [18 nt 5'-arm] - [28 nt unique sequence] - [20 nt RCA primer] - [18 nt 3'- arm] - 3' |
| **Bridge probe (108 nt)** | 5' - [readout 1] -AA- [readout 2] -AA- [28 nt unique sequence] -AA- [readout 3] -AA- [readout 4] - 3' |
| **Readout probe (18 nt)** | 5'-fluorophores- [18 nt sequence] |

We aimed to detect 67 targets in total, including 30 mRNAs and 37 proteins. We designed the barcode by assigning each target with known fluorescent readout combinations per hybridization round. We separated 14 different readout probes into three fluorescent channels (ATTO 488, TAMRA and CY5) (Table 2). Each readout probe was assigned to 16 targets. With five hybridization rounds, we detected 67 targets. In the fifth hybridization round, we repeated two channels of the first hybridization round to obtain error-correctable barcodes and validate our method.

**Table 2: Sequences of readout probes.**

|  | **Ordering code** | **Sequence** | **Ex/Em** | **Fluorophore** |
| --- | --- | --- | --- | --- |
| **1** | R1-Tm1 | GGAGCGAGAGACCATAGC | 546/579 | TAMRA |
| **2** | G1-AT1 | GGT TGA ACG CAA TGC CGC | 502/522 | ATTO488 |
| **3** | FR1-CY1 | GGA ATT GAC CGA CCG TGG | 647/668 | CY5 |
| **4** | R2-Tm2 | TTA AGC GCC CCG TAC CTA | 546/579 | TAMRA |
| **5** | G2-AT2 | GCT GTT ACG CGC TCG GTT | 502/522 | ATTO488 |
| **6** | FR2-CY2 | CCG CTA CAC GAT CGT ACT | 647/668 | CY5 |
| **7** | R3-Tm3 | TTA CGA GCG CTT GGA TCC | 546/579 | TAMRA |
| **8** | G3-AT3 | CTT AAC CGA ACT GAC GGC | 502/522 | ATTO488 |
| **9** | FR3-CY3 | CTA TTC TAA GCC GGC GGT | 647/668 | CY5 |
| **10** | R4-Tm4 | ATG AGG ACG AAT CTC CCG | 546/579 | TAMRA |
| **11** | G4-AT4 | TGT ACC GTT TAT CGG GCG | 502/522 | ATTO488 |
| **12** | FR4-CY5 | TGC TCG CAT ACC CGA TGT | 645/665 | CY5 |
| **13** | R_d-TmD1 | GCC GCT AGA AAG ACC TCG | 559/583 | TAMRA |
| **14** | R_d-TmD2 | GTC CCG ACA GTG GTT AAC | 559/583 | TAMRA |

### 2. Development of ARTseq-FISH protocol

#### 2.1 Negative controls in ARTseq-FISH

To make sure the signals were not false positives, we performed seven different negative controls in both the protein and mRNA detections. For protein detection, samples of negative controls were incubated without antibodies, or incubated with the antibodies without oligonucleotides-conjugated, or only added oligonucleotides but no antibodies. For mRNA detection, the negative controls contained scrambled (i.e., non-complementary) RCA primers or no RCA primer. Negative controls in both protein and mRNA detection include no PLP in the reactions (Supplementary Fig. 3). All the negative controls detected an average of less than 0.1 spots per cell (Supplementary Fig. 3h).

#### 2.2 Different methods of PLPs-based RNA FISH

The hybridization and ligation of the PLPs to the targeted mRNA was performed in three ways to determine the optimal strategy (Supplementary Fig. 4a-c). Cellular mRNAs were targeted either directly by the two arms of PLPs or indirectly through a bridge, which can be a linker probe bridging mRNA with PLPs, or cDNA generated by *in situ* reverse transcription. Due to the different activities of ligases on DNA/RNA and DNA/DNA hybrids, we compared three mRNA detection methods by using T4 DNA ligase or the splintR T4 DNA ligase which is known to show better activity on DNA/DNA hybrids ^4, 5^. Therefore, when mRNAs hybridize to a linker probe (L probe), which contains a hangover sequence corresponding to the PLPs, T4 DNA ligase is used to circularize the PLPs. Similarly, when mRNAs are *in situ* reverse transcribed into cDNA, PLPs recognize targeted mRNAs through cDNA sequences and are circularized by T4 DNA ligase ^6^. Alternatively, when cellular mRNAs directly hybridized to PLPs, the gaps of PLPs are sealed by splintR. We used five L probes to target *Nanog* transcript and then added PLPs corresponded to the hangover sequences of the L probes, which did not detect any *Nanog* mRNAs in mESCs (Supplementary Fig. 4d, g). In addition, similar to previous work ^3^, the relative low efficiency of *in situ* reverse transcription leads to very low detection comparing to the methods used splintR (Supplementary Fig. 4e-g).

#### 2.3 Optimizing the hybridization and ligation of RNA PLPs

As the hybridization and ligation of RNA PLPs are essential steps in RNA RCA-FISH, we sought to find ideal conditions for these reactions. First, we examined the detection efficiency which we found to be highly correlated with PLP hybridization time at both 37°C and 45°C. The longer hybridization time, the more spots per cell were detected (Supplementary Fig. 5a-d).

Next, we performed the hybridization and ligation of RNA PLPs with either Ampligase buffer or SplintR buffer. It turned out that the SplintR ligase worked better in the Ampligase buffer (Supplementary Fig. 5e-h). Comparing the ingredients of these two buffers, the only difference is that there is 0.5 mM NAD in Ampligase buffer, while 1 mM ATP in SplintR buffer. As previously reported, the product yields of SplintR are improved at lower concentration of ATP, which would explain these results^7^.

#### 2.4 Optimizing the concentration of PLPs

It has been reported that the concentration of probes contributes to the results of smFISH^8^. Here, we assessed the influence of PLP concentration in RNA RCA-FISH by performing the PLPs hybridization over a range of PLP concentration from 100 nM to 800 nM at 37°C or room temperature of ligation. We found that the mRNA detection efficiency is highly PLP concentration dependent. From 100 nM to 800 nM the spots per cell gradually increased with the concentration increasing (Supplementary Fig. 6). Although previous studies have shown that increasing reaction temperature could improve activity of splintR ^9^, our results showed that the detection efficiency of *Nanog* mRNA is higher when the PLP ligation occurs at room temperature than at 37°C (Supplementary Fig. 6). Likewise, we found that the protein detection efficiency is also dependent on the PLP concentration in Ab-FISH (Supplementary Fig. 7).

#### 2.5 Optimizing the RCA

Rolling circle amplification (RCA) is a highly efficient, isothermal enzymatic process, which makes it a powerful tool in molecular detection ^10, 11^. The length and amount of RCA products are determined by the template length, RCA time and the enzyme concentration. Here, we examined both *Nanog* mRNA and SOX2 protein with different RCA times. Dramatically increased detection efficiency was found when increasing the RCA time (Supplementary Fig. 8a-d and i). In addition, we also detected mRNA with different concentrations of Phi29 (DNA polymerase) in RCA. 0.5 U/μL and 0.25 U/μL of Phi29 in RCA resulted in considerably more RCA products compared to 0.375 U/μL and 0.125 U/μL of Phi29 in RCA (Supplementary Fig. 6e-h).

#### 2.6 Optimizing the visualization of targeted RNA

To reduce probe concentration, we also tested different concentrations of BrPs and RoPs. Similar trends have been observed for both the BrPs and the RoPs. A decrease in the probe concentration resulted in a lower number of spots (Supplementary Fig. 9). The drop in spots per cells was not directly proportional to the decrease in probe concentration (Supplementary Fig. 9). These findings indicated that increased RoPs concentrations are necessary to improve the sensitivity of mRNA detection.

#### 2.7 RNA PLP selection

To explore the effect of PLP concentration as well as the number of PLPs on detection of mRNAs, we selected three different PLPs corresponding to different regions of the *Nanog* transcript (Supplementary Fig. 10a). In each PLP hybridization, the final concentration of PLPs is 300 nM. When three different PLPs are mixed together, each PLP is 100nM in the reaction. It appears that the three different PLPs targeting *Nanog* mRNA demonstrated different detection efficiencies (Supplementary Fig. 10), and the best PLP showed better results than the combination of all three PLPs (Supplementary Fig. 10c). Although there appears to be varying efficiency of mRNA detection for different PLPs targeting different regions on mRNA, we decided to continue with one PLP per mRNA target, since we are interested in relative mRNA differences. Additionally, this kept the protocol for RNA RCA-FISH similar to the protocol for Ab-FISH. Should more rigorous quantification of mRNA be necessary at least 3 PLPs per target are likely required.

#### 2.8 Optimizing the fixation after RCA

In ARTseq-FISH, we strip off the RoPs after imaging and rehybridize another set of RoPs. We therefore tested if an extra post-fixation step is necessary after RCA. From the results, after rehybridization of a new set of RoPs in round 2, there are much less spots in channel 1 (ATTO488) in the non-post-fixed samples than the post-fixed samples (Supplementary Fig. 11). In short, it is indeed necessary to fix the sample one more time after RCA.

#### 2.9 Optimizing the antibody dilution

In order to determine an optimized concentration of the antibodies, we performed antibody dilution experiments in two steps. First for one specific protein, we performed a dilution range of the antibody to identify the concentration range in which there is a decrease in detected signal (i.e., from 1:2000 to 1:4000, Supplementary Fig. 12b). Next, we performed these same two dilutions when detecting 32 proteins at the same time (i.e., in the same hybridization round). In this case, the 1:4000 dilution shows a larger number of detected spots per cell. We diluted the antibodies with different ratios for both a single antibody and the antibody panel consisting of 32 antibodies (Supplementary Fig. 12). As a result, when the single antibody was used in Ab-FISH, the detection efficiency depended on the concentration: the higher concentration results in more detected targets (Supplementary Fig. 12).

#### 2.10 Generation of ARTseq-FISH protocol

To finalize the ARTseq-FISH experimental procedure and detect proteins and mRNAs simultaneously, we combined Ab-FISH and RNA RCA-FISH (Supplementary Fig. 2). In Ab-FISH, fixed samples were permeabilized with pre-chilled 100% methanol for 10 minutes at -20°C and blocked with 5% BSA for 1 hour at room temperature. The oligonucleotides-conjugated antibodies were then incubated for 4 hours at room temperature, followed by PLP ligation and hybridization. Once the PLPs were circularized, the RCA was performed overnight at 30°C to generate repeats of PLPs, called rolling circle products (RCPs). To visualize the proteins, the BrPs were hybridized to specific regions of RCPs first, and then the fluorophore-labelled RoPs were bound to designed regions of BrPs. In contrast to Ab-FISH, the hybridization and ligation of PLPs was accomplished in two separate steps in RNA RCA-FISH. The longer hybridization time of RNA PLPs increases the detection efficiency (Supplementary Fig. 2a-d) ^12^, while the ligation reaction of splintR takes place at room temperature instead of at 45°C for T4 DNA ligase. From the RCA step onwards, Ab-FISH and RNA RCA-FISH are performed simultaneously (Supplementary Fig. 2a). Although the spot counts per cell decreased slightly in the combined protocol compared to the Ab-FISH or RNA RCA-FISH protocol, the intensity of the spots did not change, which makes it possible to simultaneously image and analyze the spots for both protein and RNA detection (Supplementary Fig. 2b-e).

### 3. Validation of ARTseq-FISH

#### 3.1 Validation of conjugated antibodies and probes

We labelled all the antibodies panels with a unique 46 nt oligonucleotide per antibody as described in more detail in the methods section (Table S1 and S2). Before we used the whole panel of antibodies and probes in ARTseq-FISH, we tested individual antibodies by Ab-FISH (Supplementary Fig. 16) and the probes of all the mRNA targets by RNA RCA-FISH (Supplementary Fig. 17). Additionally, we quantified the spots per cell for individual targets (Supplementary Fig. 16b, 17b).

#### 3.2 Validation of ARTseq-FISH by sequentially detecting one protein

To validate the sequential hybridization by ARTseq-FISH, we detected one protein (SMAD1) with its four different RoPs. As the schematics show (Supplementary Fig. 13a), we hybridized the four different RoPs in four successive hybridization rounds, generating four different sets of images with different fluorophores (Supplementary Fig. 13b). We compared the colocalization of the detected spots in same area from four hybridization rounds (Supplementary Fig. 13b, c). Although the intensity of spots varies in different rounds, the positive signals overlapped well (Supplementary Fig. 13c).

#### 3.3 Sequential hybridization of ARTseq-FISH

In order to multiplex mRNA and protein detection simultaneously, we sequentially hybridize different sets of RoPs, image the regions of interest (ROI) of samples, and strip off the RoPs (Supplementary Fig. 13-16). We imaged the ROI of samples before and after stripping off the RoPs and compared the intensity of the same ROI. As the raw data show, after stripping the intensity of the previous positive spots dropped back to background levels compared to positive signals (Supplementary Fig. 16). To check the colocalization efficiency of the ARTseq-FISH, we rehybridized the same set of readout probes used in round 1 in the fifth round. We therefore selected the same ROI for all three channels in round 1 and round 5. Intensity plots of the selected area showed peaks (i.e., spot signal) at same locations in round 1 and 5 (Supplementary Fig. 15).

### 4. Comparison of ARTseq-FISH with immunocytochemistry

We selected several protein targets (Phos-YAP, OCT4, SOX2 and RB) and detected these targets by both immunocytochemistry and ARTseq-FISH (Supplementary Fig. 19a-d). Comparing the intensity of each protein measured by immunocytochemistry and the spots per cells of each protein measured by ARTseq-FISH, they showed similar expression levels of these proteins indicating the reliability of ARTseq-FISH (Supplementary Fig. 19a-d).

### 5. Comparison of ARTseq-FISH with smFISH

In addition, we compared the expression level (spots/cell) of *Nanog* mRNA by conventional smFISH and ARTseq-FISH. Although the expression level of *Nanog* mRNA is lower when detected by ARTseq-FISH compared to smFISH, the signals of spots in ARTseq-FISH showed 3.75-folds higher signal to noise ratio compared to those in smFISH due to the signal amplification (Supplementary Fig. 20), likely contributing to a low false positive rate in ARTseq-FISH (Supplementary Fig. 3).

### Supplementary Notes 2 (SN2)

**Computational pipeline development of ARTseq-FISH**

### 1 Introduction

#### 1.1 Overview of Raw Imaging Data (SN2 Fig.1)

In this section we outline the individual steps undertaken to analyse the ARTseq-FISH images and build a single cell dataset including explanations of the controls and assumptions embedded in the data analysis, we begin by giving 3 overviews of the raw data structure and the computational pipeline developed to build a single cell dataset.

The data for ARTseq-FISH can be summarized as follows; we micropatterned mouse embryonic stem cells, the cells are performed by ARTseq-FISH and the micropattern is imaged at 60x magnification at a resolution of 512x512 pixels using a spinning disc confocal microscope (see previous section for more details). The number of images depends on the size of the micropattern, however, to obtain a full view of a micropattern of 750 µm diameter, 6-8 connecting positions across the micropattern are captured (SN2 Fig. 1A). At each of these positions we took an image at 4 different wavelengths (colour channels), 3 to detect the hybridization signals of the probes that bind to targets and 1 to detect the nuclei stained with DAPI dye (SN2 Fig. 1B). For each of these we have approximately 100 z-stacks (depending on the number of cells that stack on top of each other) to eventually reconstitute the image in 3D (SN2 Fig. 1C). Note that this is done for every hybridization round, thus reconstituting a micropattern in 3D leaves us with number of images defined by

$$N_{I}=N_{M}* N_{p}*N_{c}*N_{z}*N_{r}$$

The number of micropatterns $N_{M}$we image, multiplied by; the number of positions $N_{P}$ we capture on each micropattern, the number of color channels $N_{c}$ sampled at each position, the number of z-stacks $N_{z}$ taken for each color channel and the number of hybridization rounds $N_{r}$. Thus, for each micropattern with 5 hybridization rounds we obtain 6 positions * 4 channels * 100 z-stacks * 5 hybridization rounds = **12.000 images**. Considering the number of micropatterns (2 different sizes, fixed at different times, replicates) we are left with >250.000 images to process, a necessary step to build a single cell dataset. This can only be done using an automated image analysis pipeline that resolves the hybridization signals and assign targets to observed signal clusters between hybridization rounds (decoding), which in turn are assigned to individual cells at each.

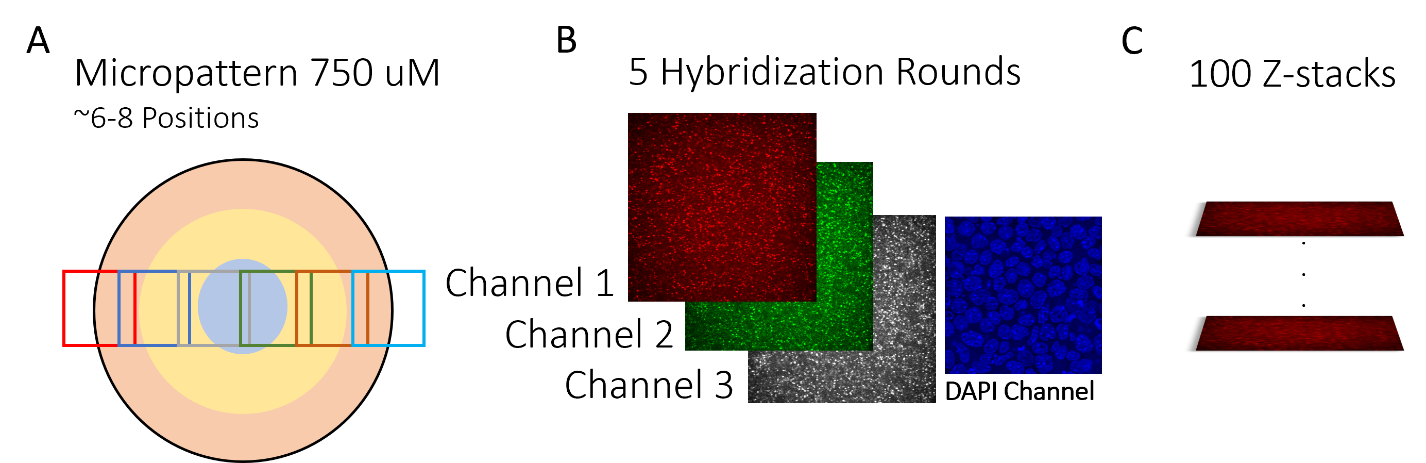

***SN2 Fig.* *1.*** ***An overview of the data collected by ARTseq-FISH.*** ***(A-C)*** *The experiments were performed on micropatterns using mouse embryonic stem cells. The cells grow and fill the micropattern, and we capture a singular line of images across the pattern using a spinning disc confocal microscope at 60x magnification.* ***(A)*** *To capture the entire micropattern we take images at 6-8 positions.* ***(B)*** *For each position we have 4 colour channels, three to match the fluorophores of the hybridization probes and 1 for the DAPI staining of the nuclei.* ***(C)*** *To capture the entire stack of cells in 3D we have around 100 z-stacks at each position.*

#### 1.2 Overview of Image Analysis Operations (SN2 Fig. 2)

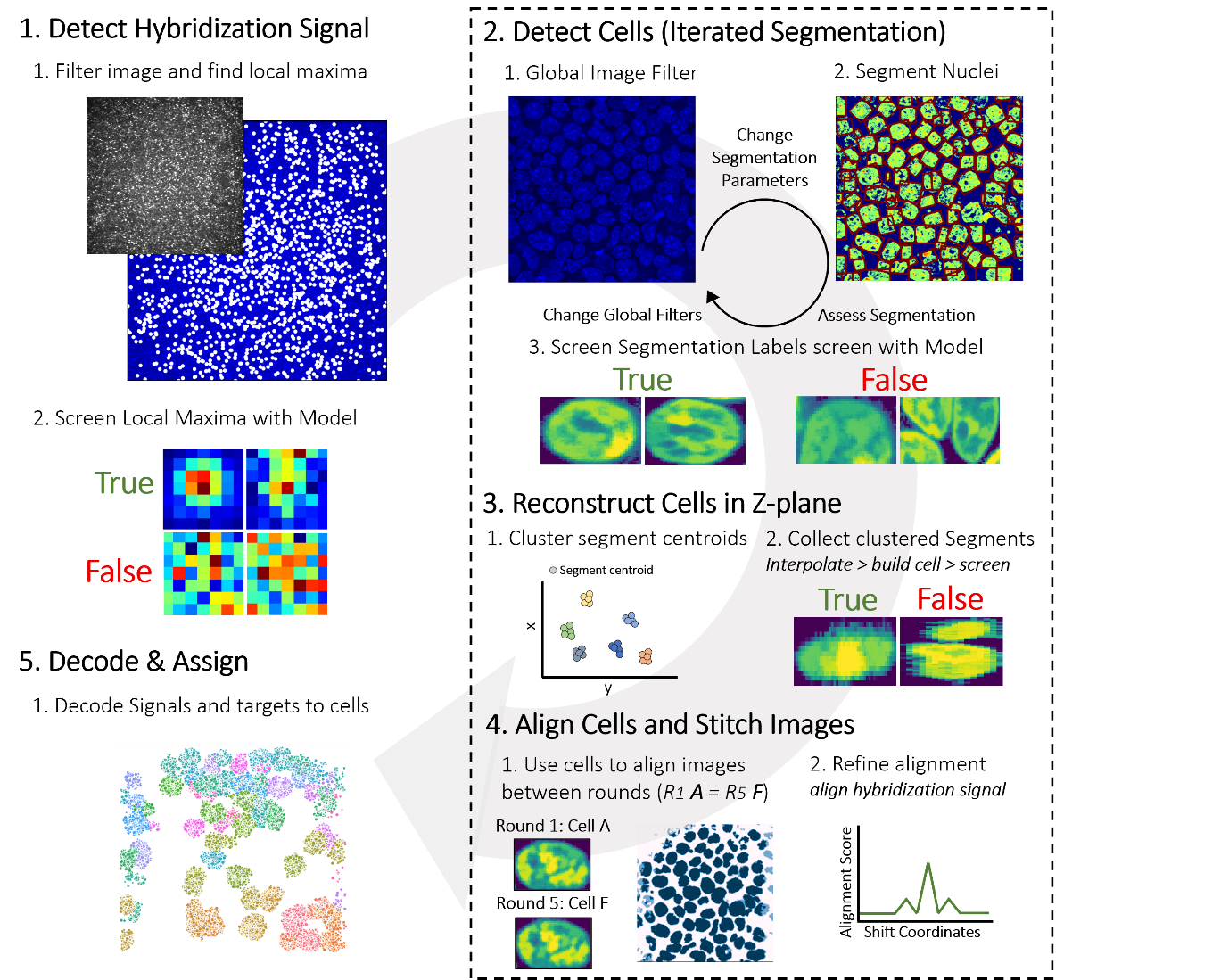

***SN2 Fig.* *2****. An overview of the Image analysis pipeline to build a single cell dataset from the ARTseq- FISH images. The pipeline is broken down into 5 steps: detecting the hybridization signal (step 1), detecting, reconstructing and aligning cells in 3D (steps 2-4) and finally decoding the hybridization signals and assigning them to specific cells (step 5). In step 1 we detect the hybridization signal, in step 2 the cells, in step 3 the cells a reconstructed in 3D, in step 4 the images are corrected for pixel drift and in step 5 the hybridization signals are decoded and assigned to cells.*

We outline the image analysis pipeline to build this dataset in overview SN2 Fig.2. We start by breaking the problem down into 5 steps. In step one, we detect the hybridization signals present in each colour channel and at each z-stack. A raw image of a single z-stack slice of the CY-5 fluorophore is shown in step 1. There is a two-step process, initially we detect the ‘speckles’ in the image that present as local maxima in the image array since these maxima correspond to a potential hybridization signal, next they are excised from the raw image. These small excisions are subsequently used to train a support vector machine (SVM) model to distinguish noise from a gaussian point spread function commonly associated with a fluorescent signal (not always, not all hybridizations appear as point spread functions, yet clear gaussian PSF’s are hybridization signals) because there is no guarantee that a detected local maximum in an image is indeed representative of a successful hybridization (even after filtering the images). For each position that is imaged, we store the location of each local maxima using 3 spatial coordinates (x,y,z) and the hybridization round.

In steps 2-4 we focus on the cells, specifically reconstructing the cells in 3D and using the cells themselves as markers to align the images (needed for step 5, resolving, and decoding the hybridization signal clusters). In step 2, the observed cells are segmented in each slice of the image z-stack. Segmentation is a challenging, therefore we make use of an iterative AI based approach, combining traditional segmentation algorithms (watershed) with an SVM model that maps onto a single cell (trained to detect successful segmentation). In step 3, we collect all the segments and reconstruct the cells in 3D by clustering their respective centroids using a node-based graph. After this we stack the segments of clustered centroids, and a 3D reconstruction is built. To screen if a single cell is actually reconstructed, an SVM model is used as a filter. In step 4, we use the cells as global markers representing an imaging round and align these rounds to the same coordinate system (initially shifted due to an incorrect repositioning of the microscope). With this, the hybridization signals between rounds can be aligned and decoded, and each decoded target can be assigned to a cell.

#### 1.3 Overview of Software (input/output) (Schema 1)

The structure of the pipeline is centred around a specific folder structure and is written in Python 3.8 (python software foundation, Delaware US), a full overview of packages used can be found in^13-20^. The pipeline processes a single position, this means each imaging position has its own folder, within these we find subfolders that contain the hybridization round which in turn contain 4 raw images with a different colour channel. Shema 1 gives an overview of the software as used to analyse the images in the subfolder and create the single cell dataset. The 3 colour channels are passed onto a module to detect the hybridization signal, the DAPI image is segmented, and its cells are reconstructed in 3D. The former passes on the coordinates of all the local maxima that were detected in the image. The latter passes on the coordinates of every pixel present in each cell, its centroid and the individual 2D segment labels. Both are passed onto the alignment module; this module takes the coordinate system of the first hybridization round as the default reference round. A global alignment operation is performed using the cells as markers, calculating the average shift of the same cells between rounds and passes this global shift coordinate (x,y,z) onto the local alignment where the hybridization signals are aligned.

The local alignment module makes minor adjustments to the global shift and stores the drift correction coordinates. This is the final alignment step, the drift coordinate is then passed onto the decoder, which resets all coordinates to the reference coordinate system. This leaves the final step, finding clusters of hybridization signals between rounds that have the same coordinates and can be decoded. In the decoding module, these signal clusters are assigned to targets, which in turn are assigned to a cell. The results are stored at each stage.

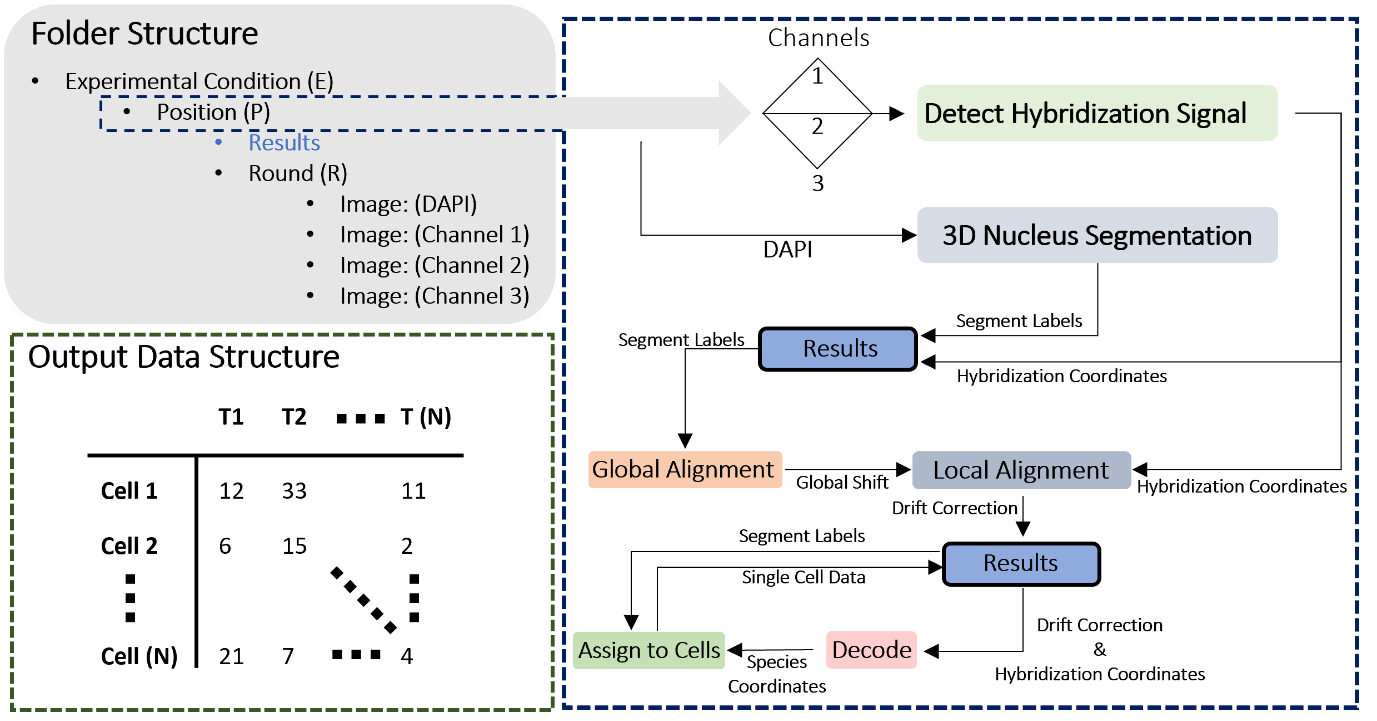

***Schema 1****. An overview of the data structures as and software modules (with input and output). The 3 colour channels (wavelengths) at which hybridization signals are detected and the DAPI constitute the raw images. The output consists of count tables where rows are individual cells and columns the species counts (T).*

2. Detecting Hybridization Signals (SN2 Fig. 3-4)

In step one of the analysis pipelines, we identify the locations of the hybridization signals in the image stacks. This is done by finding and excising the local maxima in each image, this is in itself is a multistep process. First, we assess if a clear separation between signal and background noise can be observed. SN2 Fig. 3 shows the distribution of the pixel values, specifically, 3 histograms corresponding to 3 raw images (a single z-slice) of an ARTseq-FISH experiment.

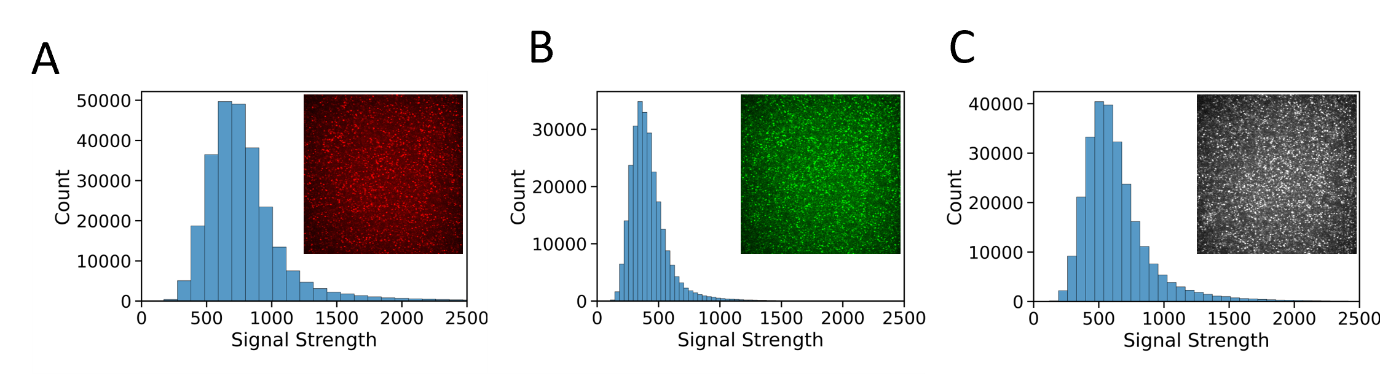

***SN2 Fig.* *3.*** *Histograms of a single z-slice for each colour channel from left to right, red, green and grey. We observed a normal distribution with lengthened right-end tails, suggesting that the hybridization signals are present in these tails.* ***(A)*** *The red channel on the left corresponds to the TAMRA fluorophore,* ***(B)*** *the green channel in the middle ATTO-488 fluorophore and* ***(C)*** *the grey channel CY-5 fluorophore.*

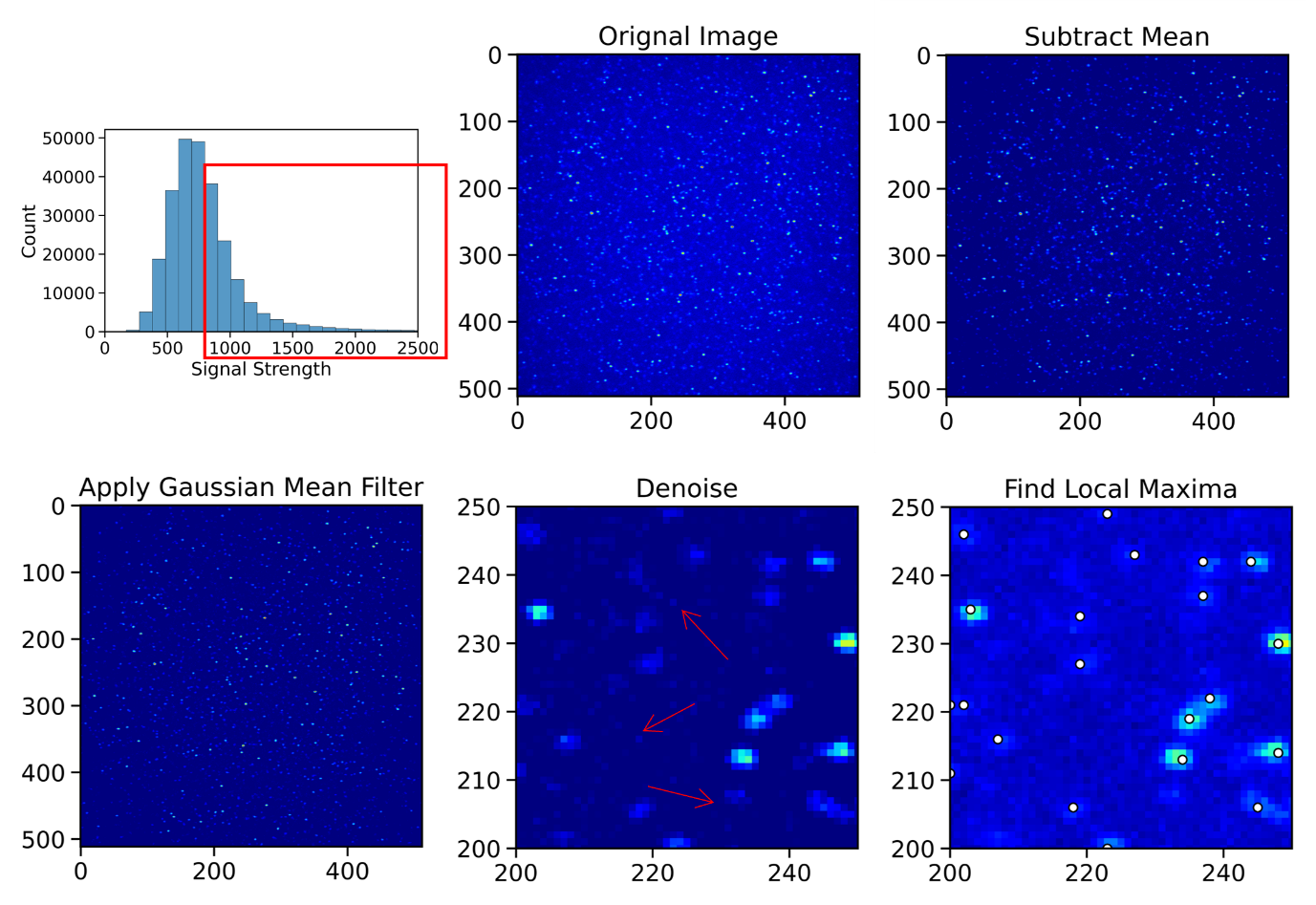

***SN2 Fig.* *4.*** *The individual image analysis steps from the top left to bottom right and their effect on the image when applied. We observe that this signal from the original image is in the right-end tail of the distributions (as indicated by the box). To isolate these signals, we subtract the mean from the original image, apply a gaussian mean filter to isolate gaussian point spread functions and separate these from noisy pixel islands. The image is denoised (examples of noise indicated by red arrows). Finally, the local maxima are left and detected (white circles). For the last two panels we zoom in on the centre of the original image.*

We do not observe a clear bimodal distribution, separating noise from signal for any of the fluorophores^21^. Instead, we observe a right-tailed normal distribution, where the length of the tails varies between colour channels, with the red channel showing the best separation of noise from the signal. Noting that the hybridization signal is present in these tails, a standardized approach is developed to find the relevant local maxima in a series of image filtering steps shown in SN2 Fig. 4^22^. This is achieved by subtracting the mean of this distribution form the original distribution shifting ~55% of the signal below 0. We subsequently state that all pixels with negative pixel values become 0, thereby removing over half the signal that was initially observed. Next, a gaussian mean filter is applied to this image. This smooths the image, intuitively this means that pixels with high values ‘overflow’ into neighbouring pixels Thus, a signal with a gaussian point spread function will retain its structure when this smoothed image is subtracted from the original image, whereas noisy pixels are isolated. In the final filtering step, we denoise the image by setting a threshold for the size of "pixel islands". If a pixel has more than 5 nonzero neighbours, it is retained. Finally, we detect the local maxima in the image using a 2D local maxima finder, in the final panel of SN2 Fig. 4 we indicate these local maxima with a white dot overtop the original image.

#### 2.1 Training an SVM Model on Point Spread Functions (SN2 Fig. 5)

This approach assumes that for the image taken, signal probes are bound to their target and fluoresce. When these steps are applied to a control experiment where no hybridization should have occurred, it will still find local maxima as a small tail and sizeable pixel islands are likely to be present regardless. To address the local maxima that are excised from the image are screened with an SVM. SN2 Fig. 5 shows the confusion matrix (SN2 Fig. 5B) and a sample the dataset it was trained on (SN2 Fig. 5A). The SVM has a binary outcome where ‘True’ is set of data containing local maxima with a 7x7 pixel excision i.e., the point spread function, and ‘False’ contains maxima excised from control experiments (images from experiments without hybridization probes) and random noise. Both of these are manually curated. The dataset for ‘True’ contains variants, mainly of hybridizations localized next to each other.

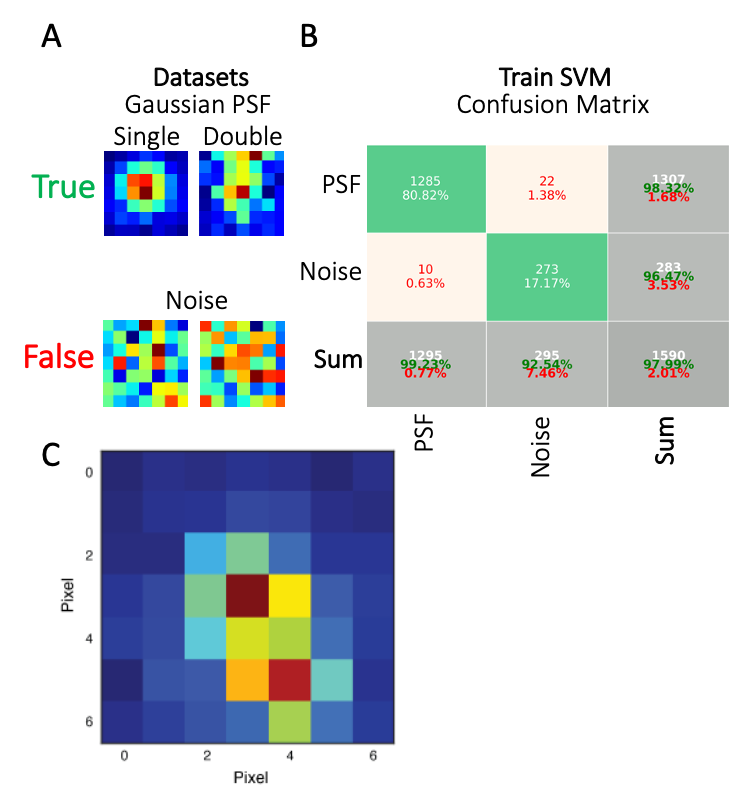

***SN2 Fig. 5. (A)*** *An example of the dataset and* ***(B)*** *the results of an SVM model trained to detect the points spread function of a hybridized signal probe. The test/train data was split 50/50 and as this is a relatively simple problem the both the number of false positives and false negatives are < 1.5%. For the point spread function dataset, variants include excisions of local maxima where a second target is within a distance of 7 pixels. The size of the datasets N(True) = 615, N(False) = 2650.* **(*C)*** *The points spread functions added to the training data (as true positive) to ensure we can detect signals in crowded conditions*

#### 2.2 Removing Duplicate Signals (SN2 Fig. 6)

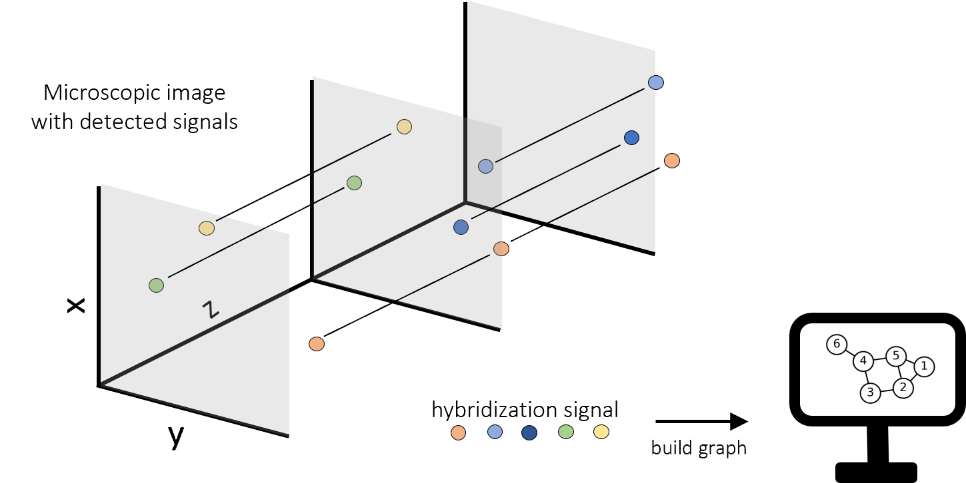

***SN2 Fig. 6.*** *The steps after excising and screening the local maxima. The z-stacks overlap, to prevent counting the same signal twice, the maxima are stacked in a node-based graph the local maxima will overlap a thus be part of the same subgraph where a single signal penetrates multiple z-stacks. Now, it is possible to keep track of duplicate signals and ensure we do not overcount a target.*

The confocal microscope scans the z-stack ‘continuously’ using Nyquist sampling. This means that there is a small overlap between connecting slices in the z-stack. Because of this the same signal can appear up to 3 times within an image. To ensure that we do not overcount the same signal these duplicates have to be accounted for^23^. This is achieved by turning the individual slices into a 3D binary matrix, where a local maximum is 1. This matrix is subsequently converted into a node-based graph where the subgraphs with more than 1 node represent repeat signals in the z-stack (SN2 Fig. 6). This is the final step in workflow to detect hybridization signals. Note, for control experiments where only 1 probe is added (targeting a single species), this is done automatically. For an experiment where clusters of maxima need to be deconvoluted and decoded, removing duplicates is the last step to ensure a target is not overcounted.

3. Reconstructing Cells in 3D

Here we outline the cell segmentation procedure and show the raw data of the DAPI stained nuclei. Specifically, we outline the type of problems that need to be solved and address them one by one.

#### 3.1 mESCs in Micropatterns and Segmentation (SN2 Fig. 7-8)

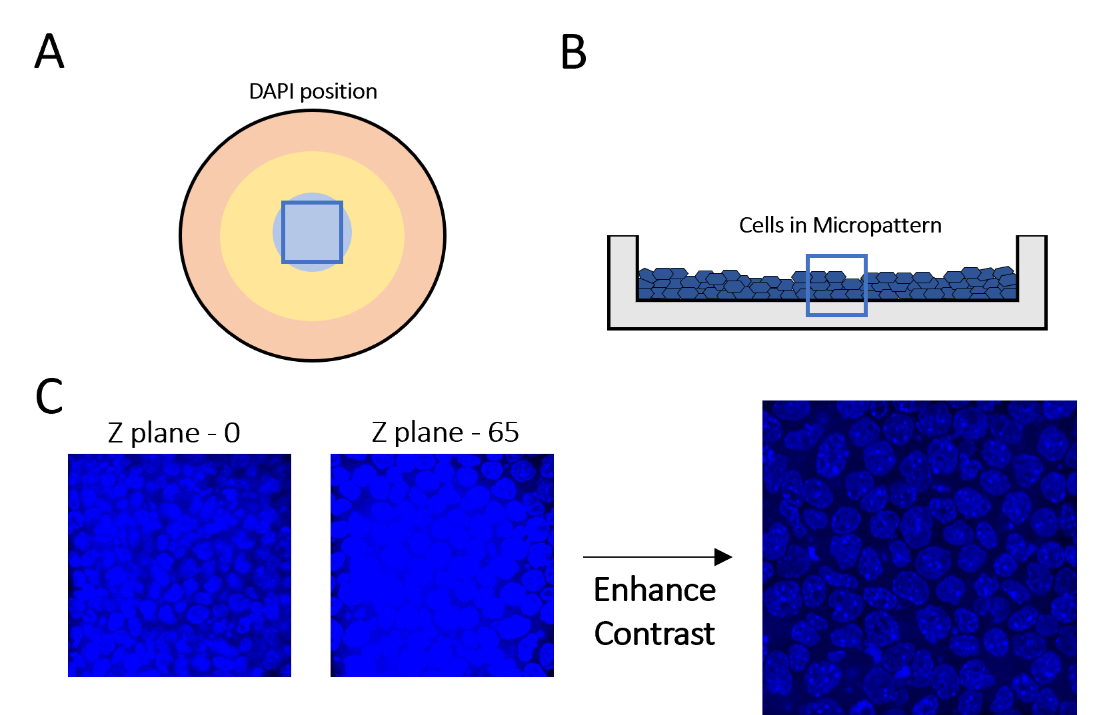

***SN2 Fig. 7.*** ***(A)*** *The figure outlines the DAPI data taken from a micropattern. A) shows the location where the DAPI image was taken (at 48 hours).* ***(B)*** *The abstraction of cells in the micropattern, the cells stack on top of each other forming several layers of cells.* ***(C)*** *Two raw DAPI images taken from different positions on the z-stack. The cells shown at stack 0 lie on top of the cells at stack 65. For both the contrast needs to be enhanced to facilitate segmentation.*

Consistent segmentation is challenging for multiple reasons^24^. First, when the mESCs divide they fill the micropattern and stack on top of each other forming layers of disjointed cells. This is illustrated in SN2 Fig. 7B and 7C in the form of an abstract representation and two raw DAPI images respectively, the raw images shown are taken at different location in the z-stack. These show that the cells detected at this height are not the same as those detected at the bottom of the micropattern, moreover the cells do not stack in clearly separated layers. Second, the images themselves can be noisy, which, amplified by the fact that cells stack on top of each other, blurs the contours between nuclei, making it harder to segment the image. This highlights the need to enhance the contrast of the nuclei contours through a set of image filter steps prior to any segmentation. Finally, we have a large distribution of cell sizes and shapes progressing through the micropattern from top to bottom caused by both the spatial orientation of a cell and the biological state it is in. Spatially, some cells are oriented vertically, penetrating deeper into the z-stack whereas others are oriented horizontally. Biologically the cell cycle phase G1, S and G2 differs between cells, which results in differences in size and brightness^25-27^.

Taking these factors into account, consistent segmentation is challenging because a segmentation algorithm that maps onto one nucleus might not map onto another resulting in only a fraction of a single nucleus (and total nuclei) in the image being detected. Yet, to faithfully reconstitute a cell in 3D we need to capture enough segments of the cell at each position in the z-stack. This ‘parameter problem’ is demonstrated in SN2 Fig. 8, where the segmentation parameters are modified which causes cell segments that were correctly segmented previously to fail, and cells that were segmented incorrectly to succeed. This indicates that a segmentation ‘model’ that maps onto all the nuclei will only work with a range of segmentation parameters sets, not a single parameter set.

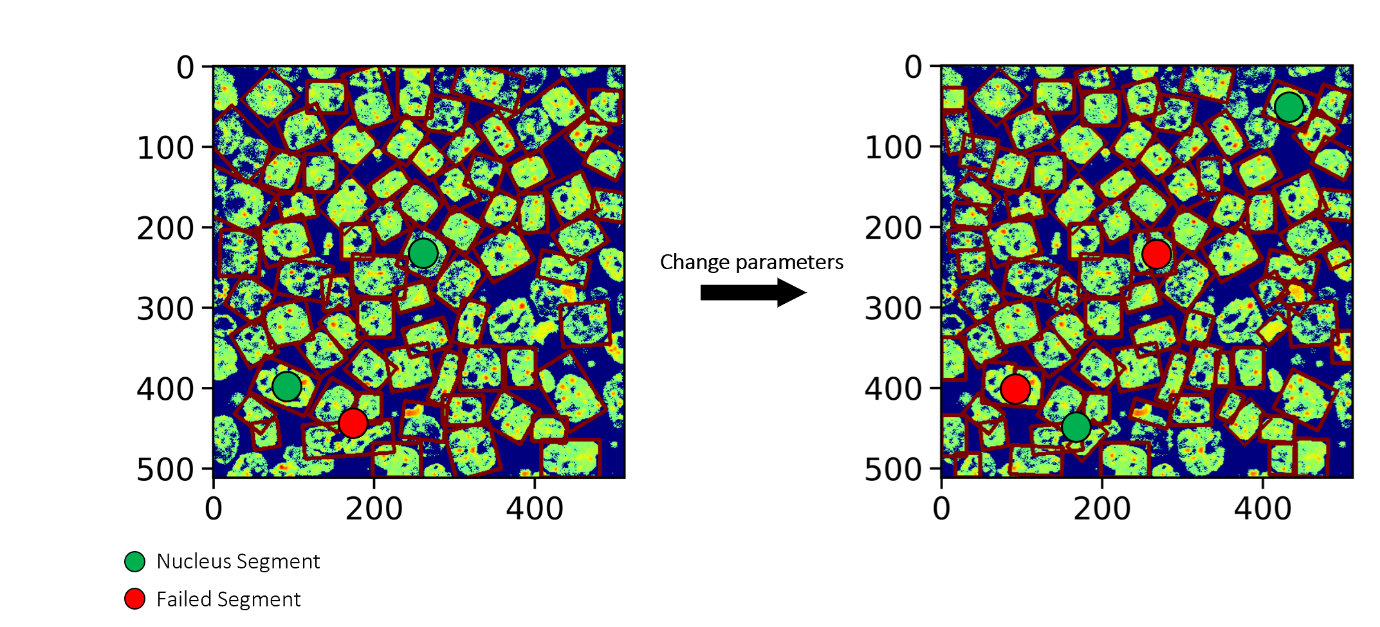

***SN2 Fig. 8.*** *The effect of changing the segmentation parameters, the red bounding boxes represent the edges of the segmentation, a green dot represents a manually chosen segment that maps onto the nucleus appropriately, the red dot highlights a failed segmentation. The only difference between the image on the left and right is a small change in the parameters chosen for the watershed segmentation algorithm. On the right we decrease the minimally allowed cell size by 10%. Note that on the left, big cells are split into two segments by the watershed algorithm, whereas double cells - initially segmented incorrectly, are now split in two. This indicates that a segmentation ‘model’ that maps onto cells requires a range of parameters instead of a single parameter set.*

#### 3.2 Iterative Segmentation (SN2 Fig. 9)

To solve each of these problems, we opted for an iterative approach, segmenting the cells repeatedly using different global image filters and segmentation parameters each iteration (SN2 Fig. 9). With this, we address differences in observed image contrast between experiments and it allows us to try a range of segmentation parameters instead of a single parameter set, ultimately maximizing the number of cell segments found. To assess if the segmentation labels found map onto single cells during this process (not half a cell, or 2 cells etc.) we train an SVM model of a cell segment in 2D. We break down each step individually, starting with the image filter steps.

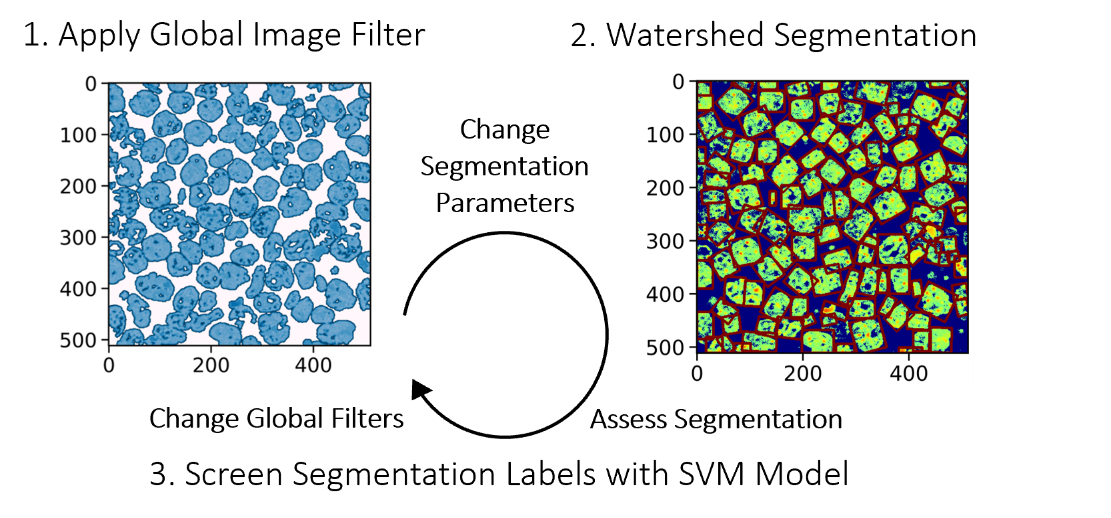

***SN2 Fig. 9.*** *An overview of the segmentation process, we apply a set of image filters that enhance the contrast between nuclei, when ready; the image is segmented using a watershed algorithm, the segments are screened by an SVM model and designated as either true or false. True indicating that the segment label indeed maps onto a single cell, false indicating that the segmentation did not succeed, usually because an individual cell is either split in two or multiple cells count as a single segment.*

#### 3.3 Image Analysis of DAPI Stained Nuclei (SN2 Fig. 10-11)

In the first step we define the set of image filters that can be applied to the DAPI stained nuclei to enhance the contrast and contours of the nuclei (which in turns facilitates the segmentation). The workflow is as follows; separate the contours of the nuclei, remove any noise left in the image, binarize the image, fill up any holes within the binarized nuclei present after removing noise. To achieve this, we apply one of three different contrast enhancement techniques, differing every iteration. Specifically, a **Contrast Limited Adaptive Histogram Equalization (CLAHE) of the image,** contour subtraction, or contour addition (SN2 Fig. 10). For each of these the parameters for individual operations can also be altered, for the contours we modify the image in 3 steps. First, the contours are extracted from the image by smoothing the image with a gaussian filter, the degree to which the image is smoothed depends on the window size taken for the smoothing mask, a small window means a very local mean is used to smooth the image resulting in a sharper image, a large window normalises the image to a more global mean resulting in a more blurred effect. The smoothed image is subtracted from the original, this leaves the contours of the nuclei, this contour is subsequently either added of subtracted from the original image (SN2 Fig. 11).

After one of the three methods is applied and the noise is removed and the binarized image is left with gaps in the nuclei that need to be filled, not doing so increases the propensity for the watershed algorithm to split a single cell into two. However, we do not want to fill the cytoplasm between connecting nuclei, therefore the size and location of all the gap-like features in the image are inventoried and a gap size threshold parameter is introduced, any gap that is larger than size N is deleted from this inventory. Again, this parameter is modified each iteration.

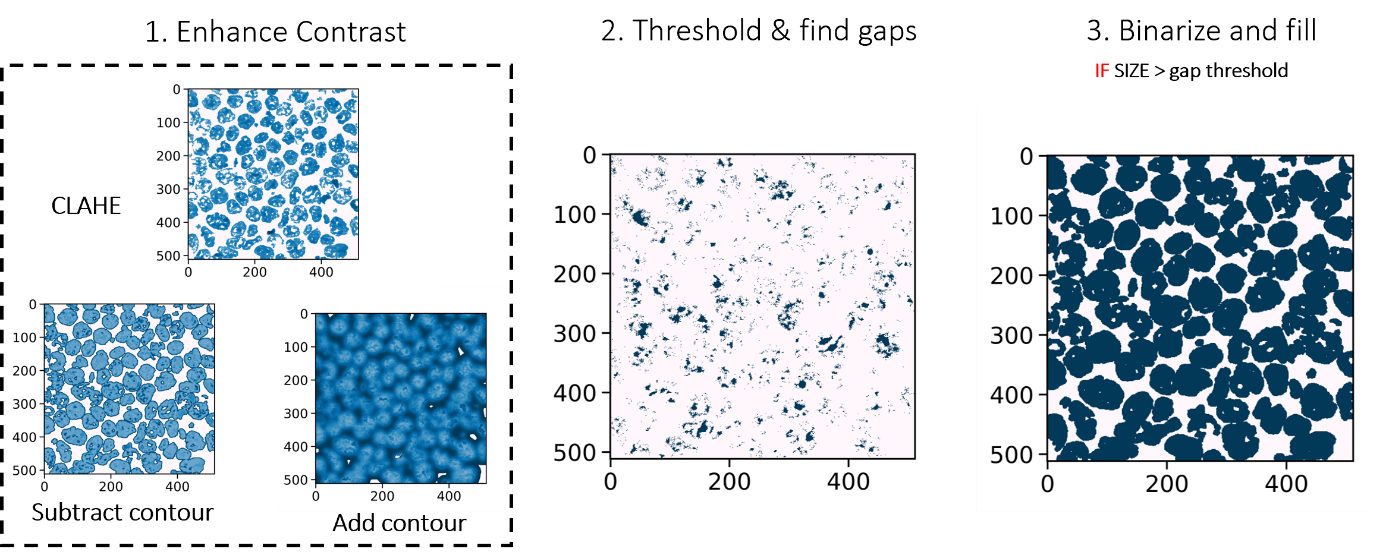

***SN2 Fig. 10.*** *Three different methods to enhance the contrast of an image, after enhancing the contrast, the image is binarized and the holes left over from the previous steps need to be identified and filled. The size and sort of gaps in the binary image depends on the method used to enhance the contrast, too large and cytoplasmic gaps in the image are filled, too small and the nuclei are left with holes which can cause issues during segmentation.*

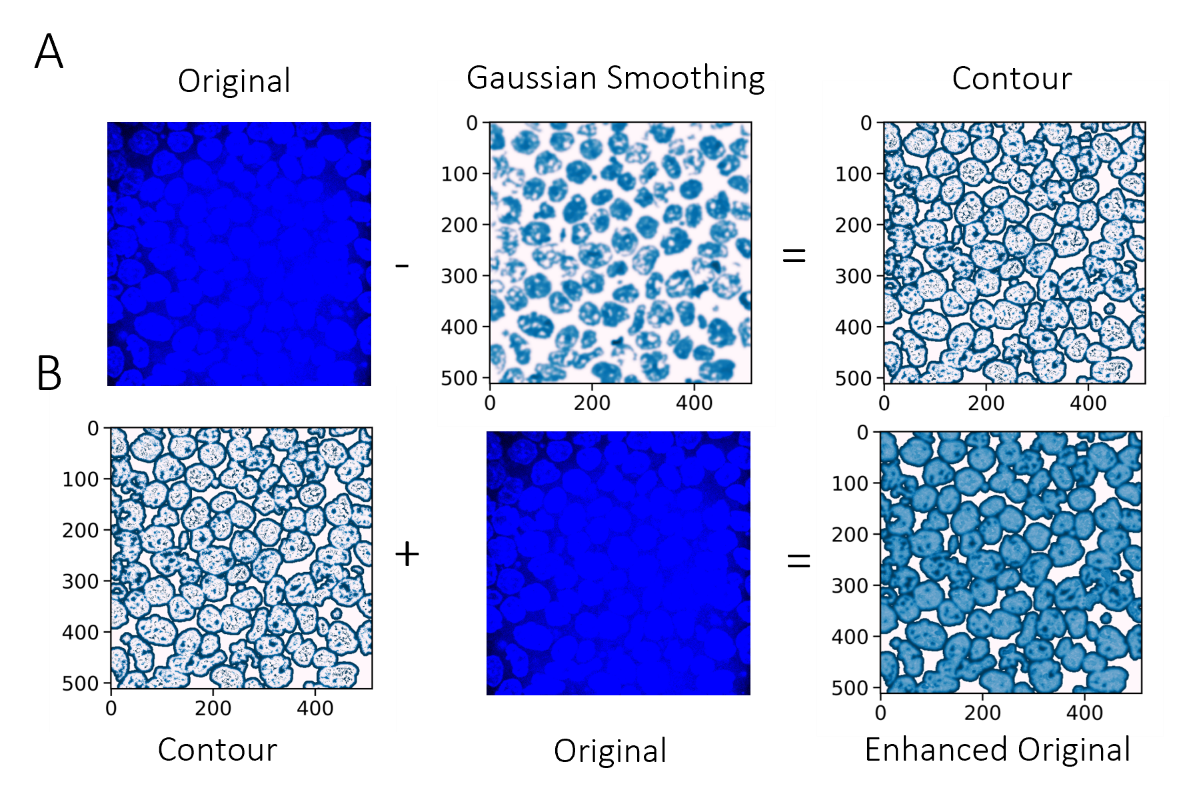

***SN2 Fig. 11.*** *The individual steps to enhance the contrast of the nuclei using the contour addition, the contours are obtained after subtracting a gaussian smoothed image from the original, the contrast of the original is subsequently enhanced by adding or subtracting the contours from it.*

#### 3.4 Segmentation (SN2 Fig. 12)

The ‘blobs’ in the binary image are segmented by the watershed algorithm, identifying the location of individual cells. As previously alluded to, the watershed algorithm has two key parameters that determine its outcome, the allowed distance between centroids and the dimensional constraints of the segments (maximum cell size). In SN2 Fig. 8 we show that different values for these parameters map onto a different subset of the cells stressing the need for an iterative approach. However, this still leaves us with a database with segments that either map onto a nucleus or have failed and thus need to be identified. To address this, we apply a similar strategy used to determine whether a local maximum corresponds to a hybridization signal (see section 1.4). We train an SVM to filter the true segments versus failed segments. SN2 Fig. 12 shows the confusion matrix and an example of the kind of data used. The dataset was constructed by segmenting 20 DAPI images from experiments both with and without a micropattern and excised an image cut-out from the raw image using the coordinates of the bounding box of the segment labels. These cut-outs have different shapes, sizes and intensity ranges; therefore, we reshape, warp and normalise the image to 1 to ensure the size and rotation of the cut-out is consistent and the intensities are comparable.

Next, we provide the model with a training and test set (0.5-0.5 test/train split) of manually curated cut-outs including either the full panel of possible segmentation failures (False) or the segments that capture a slice of a single cell (True). For the failed segmentations the most common occurrences were single nuclei split into two segments or a cluster of touching nuclei segmented as a single entity.

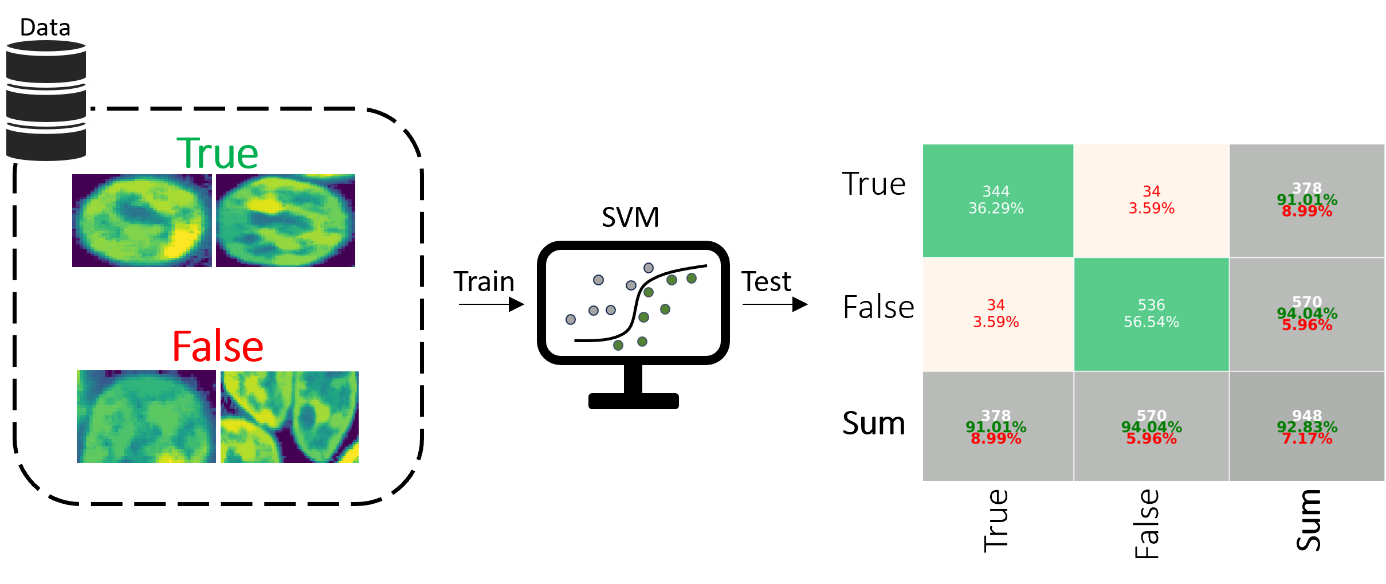

***SN2 Fig. 12.*** *The dataset and confusion matrix of an SVM model trained to recognize segmentation labels that map onto a nucleus. Both the false positive and false negative rate < 4% after the test data. The dataset itself was manually curated with >1000 segment labels in each category The segment labels that do not map onto a single cell often present as a split cell or a cluster of cells where the cytoplasm, which should be a gap between the nuclei is likely filled up during the image analysis steps. N(False) = 780, N(True) = 1115.*

#### 3.5 Sampling Segmentation Parameters (SN2 Fig. 13)

The SVM model of the segmented nuclei is used to assess if a segmentation succeeded, thus, we can score different segmentation parameters. Specifically, we can count the total number of segments detected for each segmentation parameters set and establish a range for each individual parameter, maximizing the number of detected segments.

We sampled the parameter space (Sobol Sampling) of both the image analysis filters and the watershed algorithm to find an optimal set of parameters sets^28^. SN2 Fig. 13 shows effect some of these parameters have on the number of detected segments for different taken of DAPI stained nuclei. We sampled a total of 8 different images and highlight the effect of 3 parameters on 2 images. Notably the effect of the method that is used to enhance the contrast results varies from image to image. However, for the segmentation parameters itself (i.e., the allowed distance between segment centroids and the maximum size of the cell’s bounding box) a low to mid-range value is preferred each mapping onto differently sized cells. If both become too big the total number of detected segments decreases (SN2 Fig. 13B-C). For the method chosen to enhance the contrast, the total number of detected segments varies between images (SN2 Fig. 13A). Ultimately 10 parameter sets were chosen.

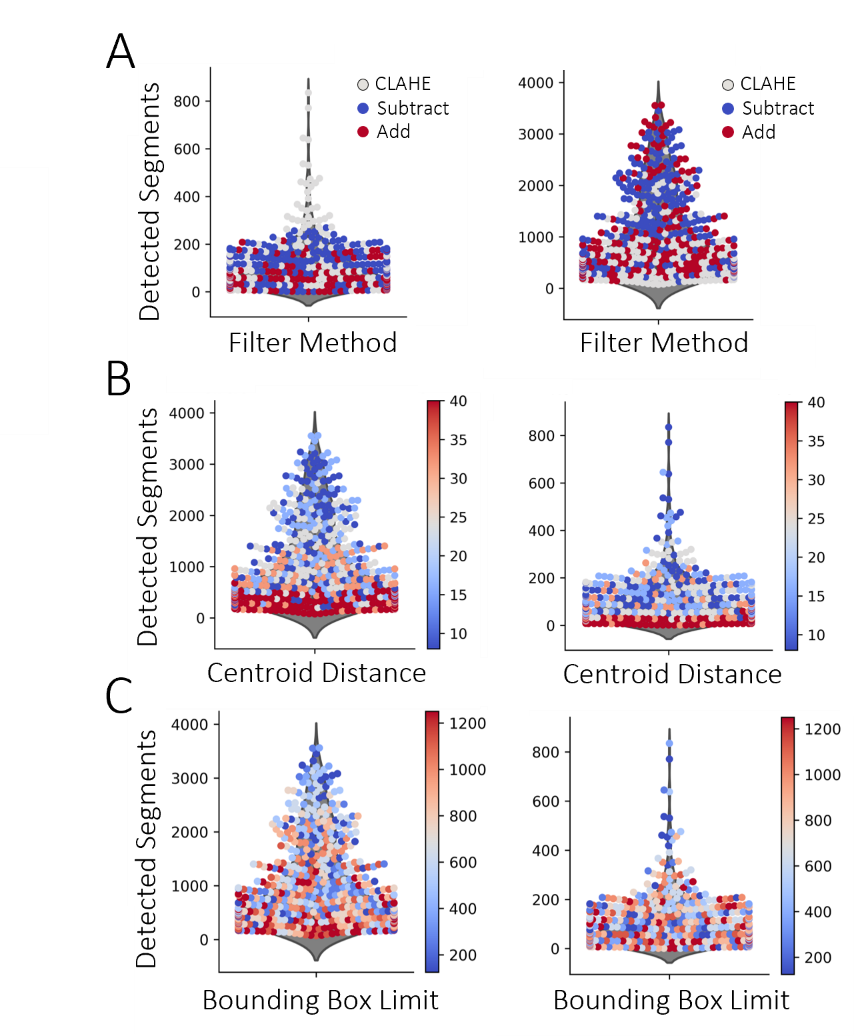

***SN2 Fig. 13.*** *Violin swarm plots with the number of detected segments (per the SVM model) for a given parameter value. The colour of the swarm plot dots indicates the parameter value.* ***(A)*** *The effect contour enhancement method has on total number of segments detected in an image; the efficacy of the method depends on the image.* ***(B and C)*** *The effect of the segmentation parameters, notably small the mid-range values need to be chosen.*

#### 3.6 Reconstructing the Cells in 3D (SN2 Fig. 14-15)

The segment labels deemed ‘TRUE’ by the SVM model, including duplicates after multiple segmentation rounds with different parameters, are used to reconstruct the cells in 3D. The labels are clustered by their centroids connecting segments between z-stacks by building a node-based graph where the minimal distance between centroids that cluster together corresponds to the segmentation parameter ‘minimal distance allowed between segments’. This step collects all the labels that likely belong to the same cell (SN2 Fig. 14). We assess the collected set of segments with a small algorithm.

1. Screen for overlap between segments
2. Check for continuity in z-stack
3. Interpolate missing segments
4. Test outcome with SVM model

In the first step, the algorithm checks if the segments that cluster also overlap. To confirm that the segments are part of the same cell, a pairwise comparison for all segment labels––across z-stacks––is performed and outliers are removed (assuming a mislabelled segment). Next, the algorithm tests if each individual z-stack has a segment that corresponds to the potential cell. If not, we interpolate an approximated by averaging the closest neighbouring segments. For example, if no segment is detected for a potential cell at z-stack *n* we interpolate segment *n+1* and *n-1* to approximate the shape of z-stack *n*. Finally, the reconstructed in 3D cell is obtained by stacking the segments on top of each other in a binary 3D matrix. However, that is no guarantee that each cluster is representative of a single cell, it is possible that cells stack directly on top of each other, and a faulty segmentation label remains part of the cluster. To solve this, we introduce another screening step, by––again––using an SVM however, this time the segment labels are based on a side view of a nucleus (SN2 Fig. 15), the reference image is still 2D and still uses the raw DAPI image. The validity of the reconstruction is tested by taking a slice along centroid value of the x-axis and excising the coordinates within its bounding box from the raw DAPI image. This raw DAPI image is again warped, resized and the intensities are normalised to 1.

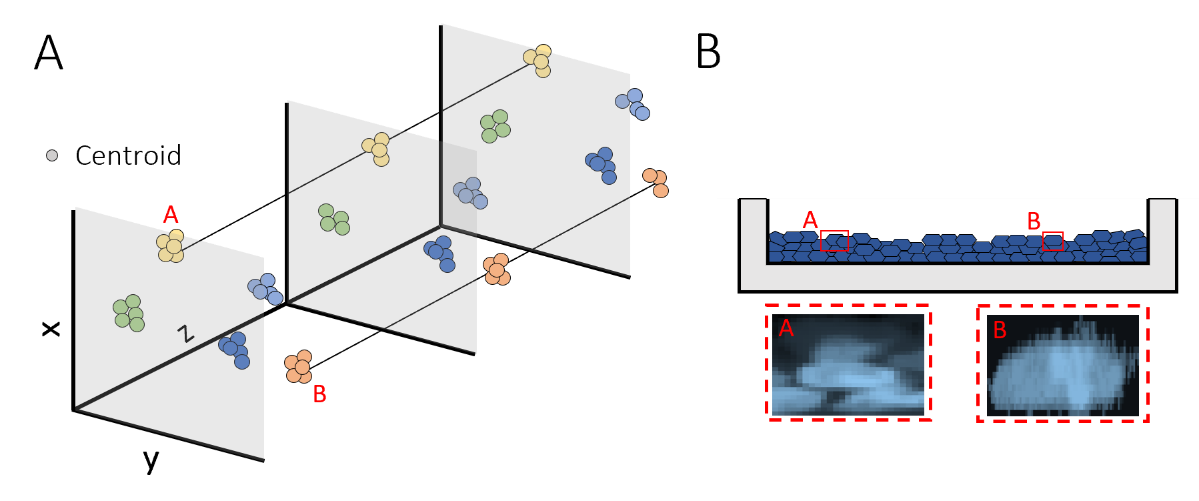

***SN2 Fig. 14.*** ***(A)*** *The centroids at each z-stack that cluster together.* ***(B)*** *A sideview of the cells as they are reconstructed where an example of a failed and successful reconstruction is shown (these are subsequently used to train an SVM to recognize failed and successful reconstructions).*

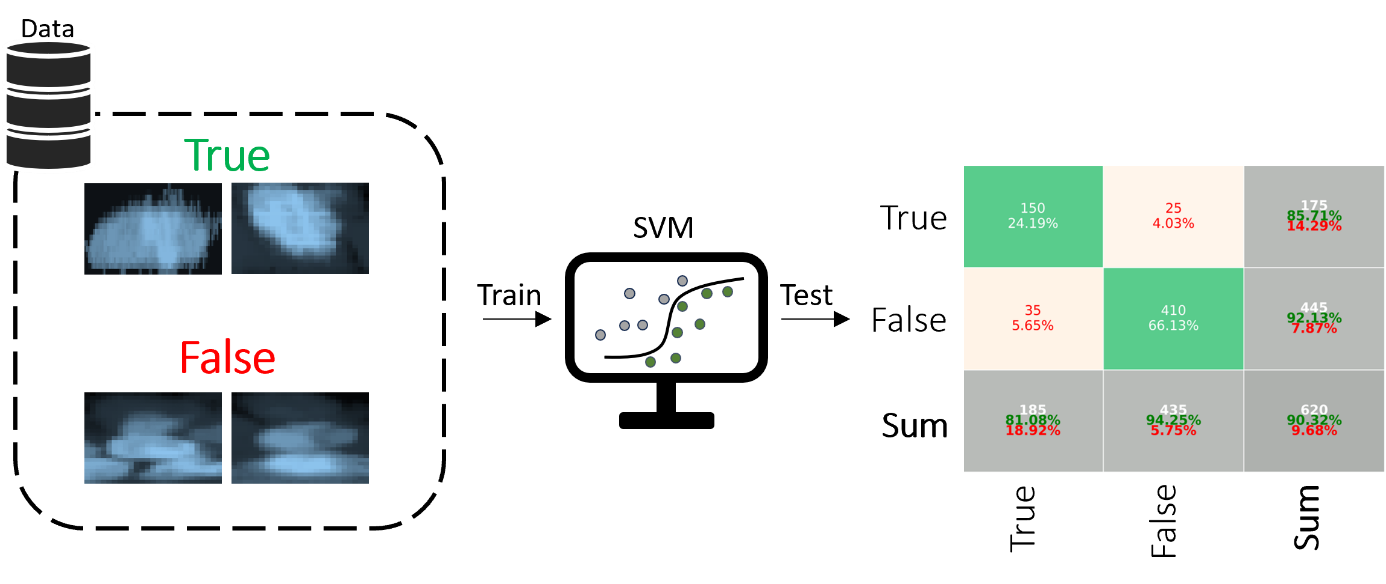

***SN2 Fig. 15.*** *The dataset used to train the SVM, the failed reconstruction of occur because cells stack right on top of each other. The confusion matrix, the SVM was trained using a 0.5-0.5 split test/training data. The test and training data is pre-processed, the cut-outs are warped and resized, and the signal values are rescaled between 0-1. N(False) = 380, N(True) = 860.*

#### 3.7 Aligning the Coordinate Systems Between Rounds (SN2 Fig. 16-17)

The hybridization signals that are detected in each round can only be decoded if they share the same coordinate system. Therefore, the images between rounds need to be aligned. This follows from the global drift in imaging positions. Each round we hybridize the readout probes to the bridge probe and move the camera to 6-8 pre-programmed positions (that slightly overlap) to image our cells across the micropattern. We subsequently strip the hybridization probe, re-hybridize a new read-out probe and repeat the imaging. This means we move the camera up to 40 times which in turn causes shifts between the same position for different rounds. Moreover, the drift of individual pixels also needs to be addressed, where a local signal maximum appears in any pixel within the point spread function, though this is significantly less pronounced than the global shift.

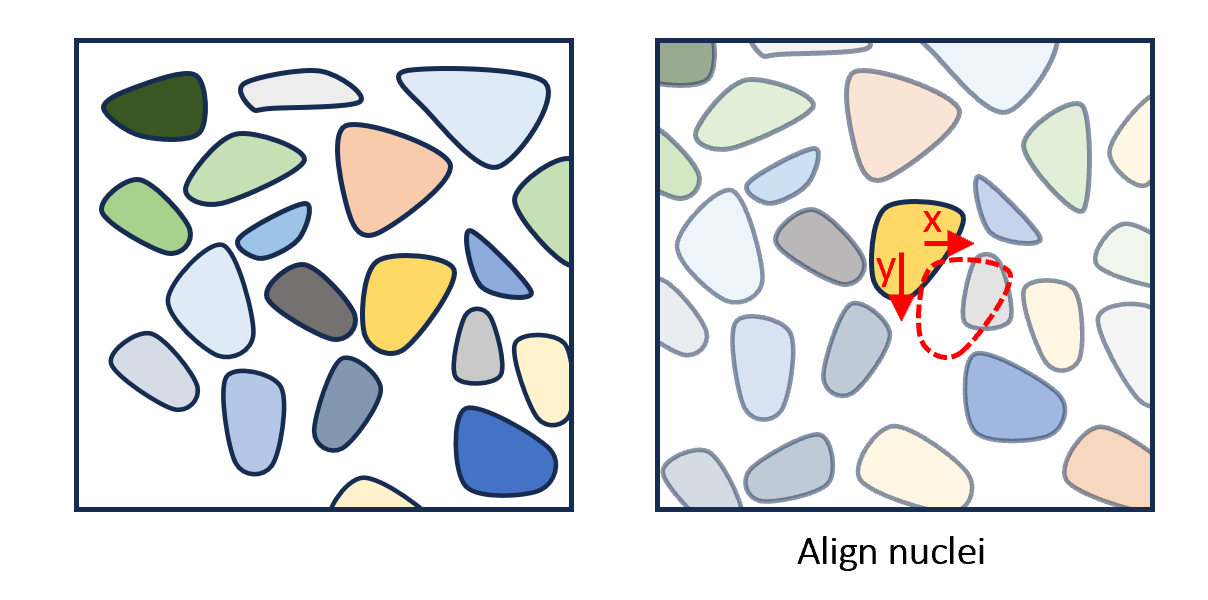

***SN2 Fig. 16.*** *The global alignment compares cells in the reference round (whose coordinate system will be used) to the cells imaged during the other hybridization, scoring the likeness using a sum of squares fit function, summing the difference of all individual pixels. The shift coordinates of cell pairings with the lowest scores (and are close to one another) are averaged. The likeness is only scored if some heuristic conditions are met with respect to cell dimensions, specifically cell size. If the length and width of 2 segments are too far apart (25%), the comparison is removed. b) The shift as applied to the image where the dark blue areas indicate overlapping structures.*

The drift problem is solved by applying 2 sequential alignment operations. For the global alignment we make use of the previously found cell label segments as markers. Each cell within our sample is akin to a fingerprint, unique in shape and its distribution of DNA which means it can be as a global position marker (SN2 Fig. 16). We sample a subset of the cell segments at multiple locations in the z-stack from hybridization round 1, considered the reference round. The cells segments found in all the other rounds are subsequently compared to the reference round to find a matching pair using a sum of squares scoring function and some heuristic metrics like original cell size as a pre-screening step. Because the shift between all cells should be consistent the best scores of different cells shift in the same direction. we subsequently take the average shift of centroids that cluster together with the best scores, this removes potential outliers (misaligned cells) present.

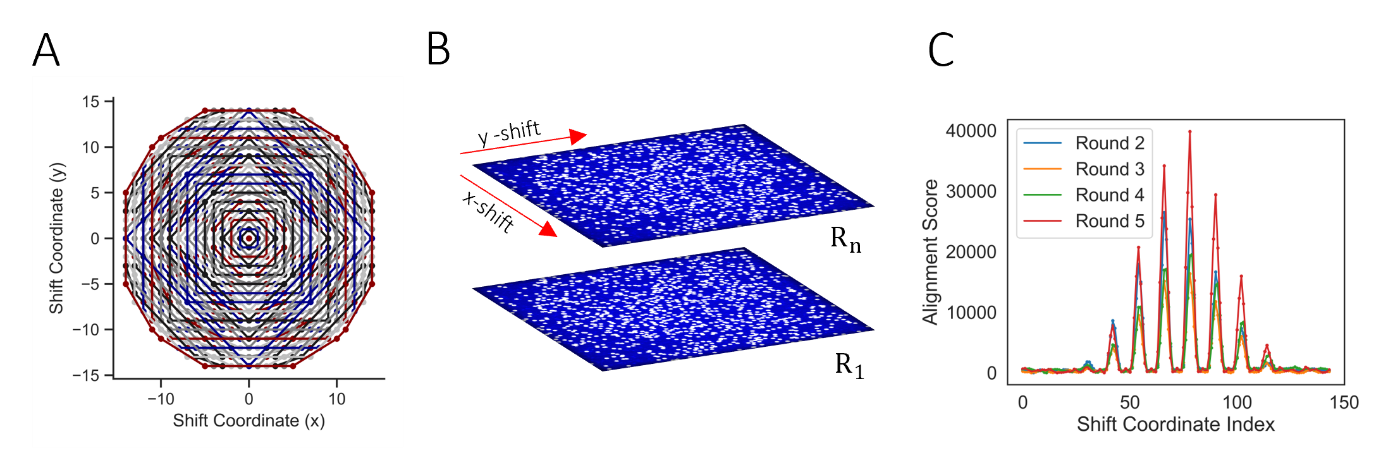

***SN2 Fig. 17.*** ***(A)*** *The search pattern, starting from the centre,* ***(B)*** *the image to be aligned to the reference round (R1) is shifted overtop the reference round and the overlap between hybridization signals is scored.* ***(C)*** *The overlap between rounds 2-5 and round 1, the scores shown are corrected for spots that overlap by random chance.*

After the global alignment the hybridization signals are within a few pixels of perfect alignment. To achieve a ‘perfect’ alignment and prevent misassignment, the individual hybridization signals are aligned in a second alignment step. A 4D binary matrix is built using the hybridization of every round (their coordinates updated after the first alignment), each hybridization signal is represented by 9 pixels, the coordinates of the local maxima including 8 neighbouring pixels representing a minimal point spread function. The images are subsequently shifted along a circular grid (SN2 Fig. 17A-B) over the reference round until we find a set of coordinates where we count the largest amount of overlap. The overlap is scored for each shift along the grid. An example is shown in SN2 Fig. 17C where the alignment score increases when the optimal shift coordinates are found (local maxima when part of the point spread functions overlap). Note that this operation is crucial to deconvolute the signal. If the pixel shift is too large after the global alignment the targets would need a larger target area. This area (number of pixels) is defined as belonging to a single target and once stacked, results in the ‘detected’ hybridization signals.

4. Decoding Hybridization Signals (SN2 Fig. 18-19)

To decode the hybridization signals in the images, the detected codes (i.e., the hybridization signals that have the same coordinates set between rounds) are referenced to a codebook. This codebook comes in the form of an excel sheet with the expected colour channels present in each round of hybridization of each individual target. Experimentally, it represents the order in which hybridization probes are added to the sample. We have 3 fluorophores representing the different colour channels, split over 5 rounds. On some occasions multiple fluorophores hybridize to the target sequence, and we expect 3 overlapping signals in a single round. This means we have 7 potential codes per target per round (Red, Blue, Green, Green & Blue, Green & Red, Red & Blue, Green & Red & Blue) or 16807 potential targets.

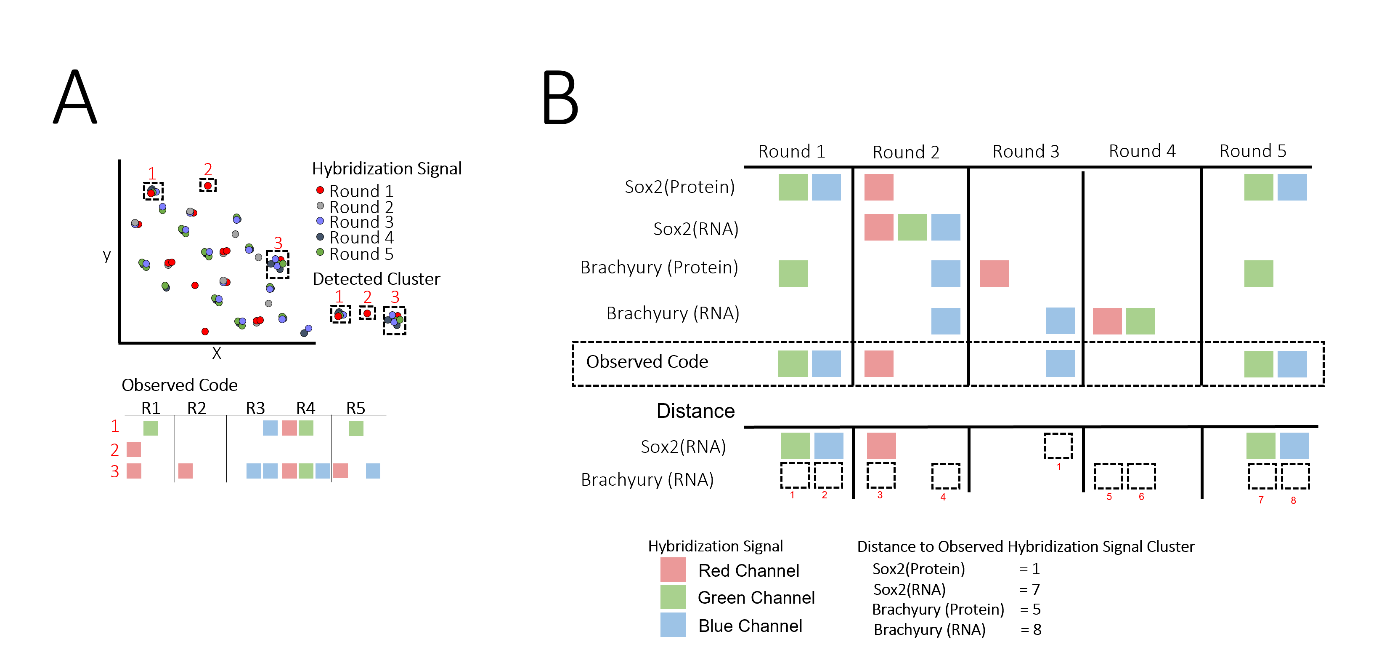

***SN2 Fig. 18.*** ***(A)*** *An abstraction of detected hybridization signals from different rounds clustering together after alignment. The observed code can be matched to the codebook shown in* ***(B)****. the distance between an observed code and a detected code is calculated by summing all the signals that do match. In the example above, SOX2 Protein has a near perfect match with the codebook (one extra signal Is detected in round 3). Whereas Brachyury mRNA is furthest removed from what is observed.*

The codebook design balances the distance between codes versus the number of hybridizations rounds under the assumption that a crowded signal would make it difficult to distinguish targets between rounds if there is a base level of noise present. By doing so we increase the certainty that an observed code is true. To deconvolute the observed codes a graph of the codebook is created where the distance between nodes reflects the distance between codes. The set of all possible observations and their distance to an existing code is included in this codebook. Thus, if we observe a signal that does not match 1 to 1 with the codebook, we can find its nearest neighbour as defined in SN2 Fig. 18B. Where a hybridization signal, either missing or present but not in the codebook counts as +1, and the shortest path to a code is likely to be the target. However, this comes with a caveat, namely that assigning codes in this manner will results in mis-assignments. Particularly, in cases where multiple targets are equidistant to a detected code, or two targets within each other’s point spread function and therefore have hybridization signals that overlap.

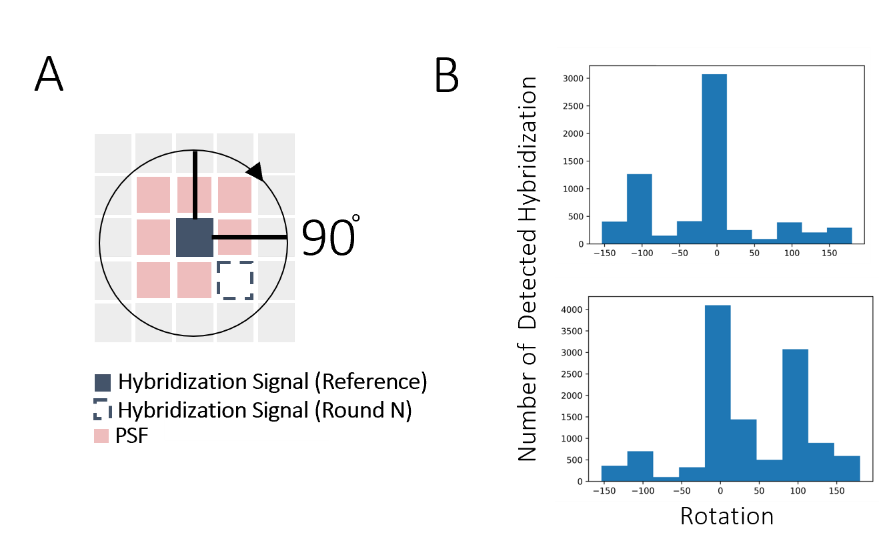

***SN2 Fig. 19.*** ***(A)*** *One of the metrics, ‘rotation’, used as a predictor, namely the rotation of targets vis a vis a hybridization signal from the reference round. There might still be a slight bias after aligning the images e.g., where hybridization signals from rounds 2-5 favour a specific position if not exactly on top of the reference caused by the true shift is rounded of the then nearest whole pixel, this can cause pixels between one of these rounds, relative to the reference round, to consistently favour a second pixel for all signals.* ***(B)*** *The distribution of the signal rotations of two rounds relative to the reference round.*

We address this by decoding the detected signals in steps, using the information gained at each step about the structure of the observed codes. Thus, if a detected code does not match a code in the codebook and it is not clear where it can be assigned, this additional information serves to build predictors. The following factors are considered:

- The frequency a specific fluorophore is detected outside any observed code (false positives)
- The frequency they are detected in an observed code
- The position number of the local maxima with respect the reference channel signals
- The relative position of different fluorophores presents in the same pixel
- Initial distribution of detected species

The initial distribution of detected species represents the observed codes that are a 1 for 1 match with the codebook, these serve as a sample reflecting the relative percentage of species present compared to one another. The steps follow schema 2, if there is no direct match to the codebook the next step tests if the observed cluster of hybridization signals reflects two separate hybridization signals. Next, we assess if the observed code is larger than any code found in the codebook but cannot be split in two, likely caused by the introduction of unneeded local maxima. These signals are screened by a set of metrics to assess what the odds are that a specific signal can be considered noise. For example, if the dataset contains a significant number of local maxima in a channel wherefore a lot of these maxima have been detected in isolation it is likely that this is noise and vice versa. Moreover, the hybridization of multiple fluorophores to the same target in the same round can serve as a benchmark as these signals have the exact same pixel coordinate between channels. Thus, if multiple fluorophores are detected within the same round at a certain location but they are distanced further apart, it indicates that only one is likely to be bound to the target. Finally, the spatial rotation as seen from the mean position of spots between the reference round and a round to be compared serves as another marker.

We observed that a very minor global pixel drift between each round and the reference round persists after the image alignment. Concretely, this means hybridization signals between for example round 1 and round 2 either overlap or the maximum intensity falls in neighbouring pixel with a 90-degree rotation, between round 1 and round 3 this rotation can be 270 degrees (SN2 Fig. 19B). This means, that if ever there is a conflict between two hybridization signals (i.e., one must be deleted) then these become the deciding factors. Finally, if a detected code is of at least size 4 and the difference between this observed code and code in the codebook is 1, it is assigned.

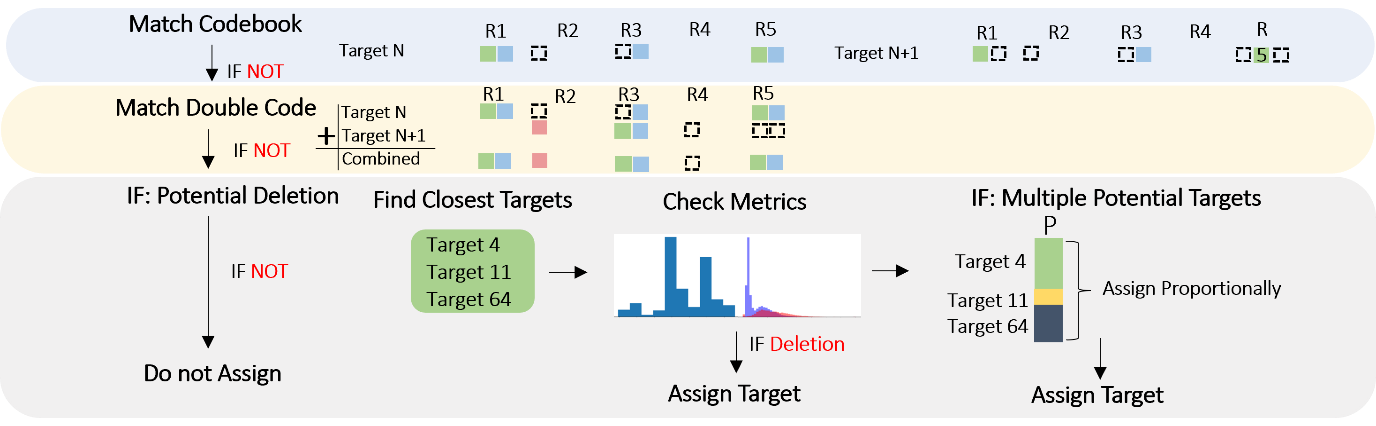

***Schema 2.*** *Overview of the steps taken to decode an observed signal cluster. First, the observed code is matched against the codebook, if a match if found a target is assigned, if not, the code is matched against all possible combinations of codes assuming the signals are close together, if not, we assess if hybridization signals can be deleted from the observed code under the assumption there is a noisy signal. If not, not nothing is assigned to this cluster, if true, the closest targets and its corresponding required deleted are screened against a set of metrics, if multiple targets are still feasible, all possible targets are assigned to a signal cluster, these are assigned proportionally in the final step depending on the relative abundance of each species.*

5. Data Analysis
   5.1 Experimental Reproducibility (SN2 Fig. 20)

With the single cell dataset new analyses can be done. Firstly, reproducibility, the bulk reproducibility of repeat experiments is assessed for 3 timepoints. The data are shown in SN2 Fig. 20, including 4 replicates comparing both the edge and the true centre. The correlations are high, especially for the final timepoint. We have shown that the abundance of targets is––in part–– predicated by cell density, as seeding is not unique and semi random clusters of cells are present in the image, thus it is likely that more variance between replicates cam be observed especially at time points where the system is in a more transient state. Overall, the correlations are very strong (0.5-0.95) between replicates. Demonstrating that experiments are reproducible, the same needs to be done for the analyses.

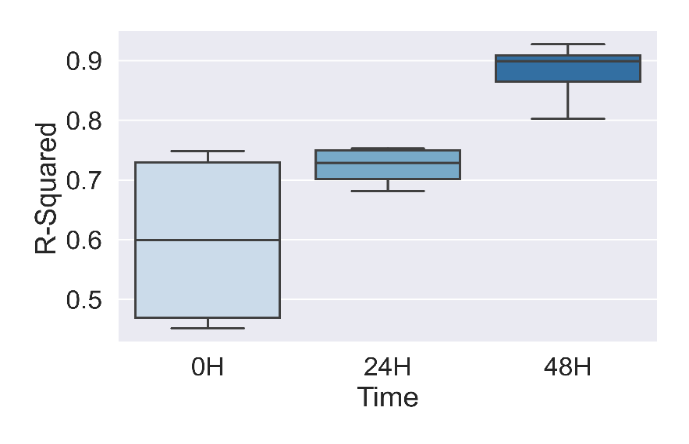

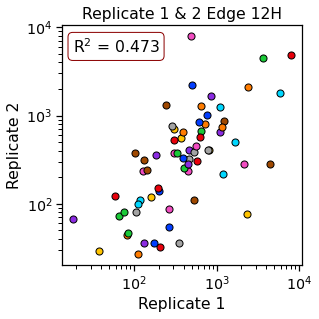

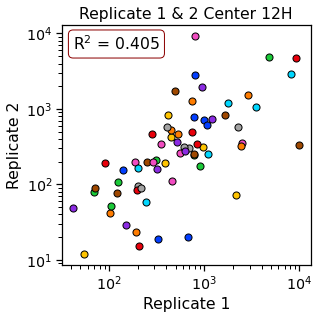

***SN2 Fig. 20.*** *Boxplot plotting the mean r-squared value between total target counts of different replicates. The expression differences between experiments converge over time, where at time 0 has the most variance between batches is observed and after 48 hours the least, these data contain 6 position replicates, between 2 repeat time series experiments taking both the edge and the true centre of the micropattern to compare. Line in the boxplot denoting the median (left). On the right we show the correlation between edge and centre for the cells measured at 12H, the lowest values but similar to 0H indicating that the cells are still in flux (right).*

5.2 Data Normalisation for Data Analyses

Single cell analysis can be sensitive to the normalisation method and an abundance cut-off applied. To validate any result from a technical standpoint it is therefore prudent to apply both different filters to the data and double confirm a result with a different analysis method. If these analyses come to consensus, it can be assumed that the patterns observed consistently e.g., after clustering with a UMAP (uniform manifold approximation projection), or mapping the changes of target groups, are not an artefact of or overly sensitive to the mapping technique with respect to the dataset or the data processing itself. For most analysis, only an abundance threshold is applied (this threshold automatically filters out cells not reconstructed properly and reduces the impact of repeated 0 counts). For the UMAP analyses the data is normalised to reflect a fold change instead of an absolute count. For the violin and swarm plots in Figure 1 and 2 of the main text, we specifically choose not to apply a normalisation or introduce a threshold to provide the reader with an unfiltered sense of the dataset. This means a small number of cells with a very low total count––affecting the mean––were included in those figures (potentially due to failed reconstructions of the cell). For reference, 48 cells were reconstructed at the very edge of the micropattern, 12 cells did not have more than a total of 300 detected targets, of these 12, 50% have fewer than 100 spots.

**Abundance Threshold:** Cells with fewer than 300 targets (total) are removed from the dataset.

**Data Normalisation:** A log (2) normalisation of the target counts is applied prior to **clustering.**

Next, we 5 briefly describes the data analysis for the validation data of figures 4-7 in the main text:

- Analysis of figure 4e. We divide the mean number of detected spots per cell detected at the edge by the mean number of detected spots per cell in the centre. If the difference is smaller than foldchange < 1 we take the inverse and normalise to 0, thus the reported foldchanges are the same and have to be sorted category to know if edge or centre is favoured.
- Analysis of figure 5c. We binned the number of detected spots on the micropattern from edge to centre, notably, the bins include the average of relative number of spots per species in the group and normalised these for the number of cells by taking a middle z-stack slice. We subsequently plotted these bins in a hexplot. The bins are normalised for the maximum abundance (i.e., bin with highest value) across all 4 time points.
- Analysis of figure 5e. We take that relative increase or decrease of each species counted per cell (normalised to timepoint 0) and average these relative changes for each group of targets that regulate a similar process e.g. cell cycle 1 and cell cycle 2 etc. The shaded around the mean (thick line) represents the 95% CI. Note that the relative change is with respect to time zero.
- Analysis of Figure 5e. Quantifying NANOG expression does not require segmentation we sample 5 positions on each image and calculate the average intensity of these positions. We note that the small deviations from the observed mean indicate that the positions were sufficiently representative of the image at large with being skewed by areas without cells (as would be the case for the whole image especially at the edge). The mean and variance (95% CI) of these 5 positions are plotted in the main figure highlighting a difference in expression.
- Analysis of Figure 6c, e, f Proteins synthesis intensity across the micropattern and DAPI intensity across the micropattern and normalized DAPI intensity across the micropattern at 0, 12, 24 and 48 hours after LIF withdrawal resulting in donut shaped curve. The average intensity is calculating by calculating a moving average across the micropattern from left to right, the centre of the moving window size moves through the centre of the micropattern and has a radius of 50 pixels.
- Analysis of Figure 7g We filtered and segmented the cells in the image, cells with sufficient activity will register as an independent segment in the image. These segments are counted at each timepoint. The lineplot shows the interpolated average and the individual counts registered by the segmentation algorithm. The cell can either have a red, green, or mixed signal (i.e., it registers in both colour channels).
  5.3 UMAP of Single Cell Data (sensitivity to analysis method) (SN2 Fig. 21-22)

The outcome of a UMAP applied to single cell dataset depends on 2 parameters: number of graph connections and distance of connection, and more importantly and filters applied to the data itself, which, in turn, determined by the scaling/normalization method chosen for the dataset and the cut-off (i.e. which cells are counted). Because the dataset contains both mRNA and proteins there is a lot of variation in expression levels.

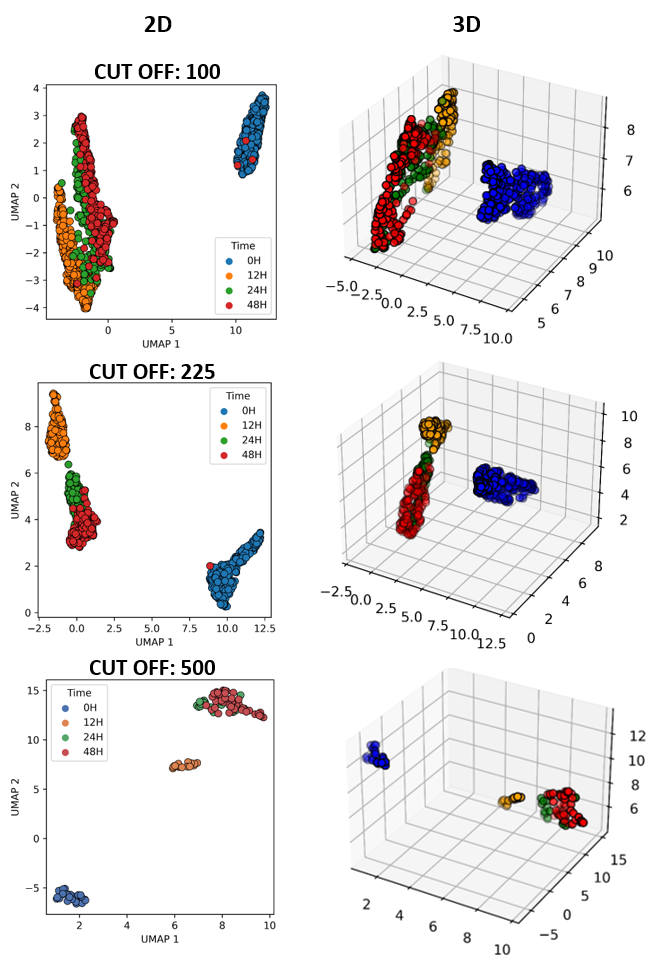

***SN2 Fig. 21.*** ***(A)*** *The result of using a different abundance threshold in two dimensions. We observe an increased variance in the grouping of the cells when we account for cells with a total spot count of less than 100.*

SN2 Fig. 21. Shows a repeated the UMAP analysis for different data sets (applying different filters and normalizations). We plot both the 2D and 3D projections for a UMAP of the dataset to determine the robustness of the data separation (SN2 Fig. 21A). The abundance threshold for the minimal number spots per cell is tested, ranging between 100-500 (SN2 Fig. 21C). We finally chose 225 as an abundance threshold (for any value above this threshold the analysis does not change drastically, below this value more 0 events are added increasing the overall variance in the data). In SN2 Fig. 22 we tested if a different normalization of the data results in different groupings in the UMAP, however this is not the case.

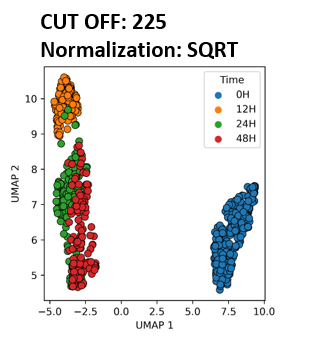

***SN2 Fig. 22.*** *The result of using a different normalization taking the square root of the data instead of log2 for a 225 cut-off. We observe no significant difference between the data grouping using either log2 or a square root normalization.*

5.4 Correlation of nuclear versus whole cell spots count data (SN2 Fig. 23).

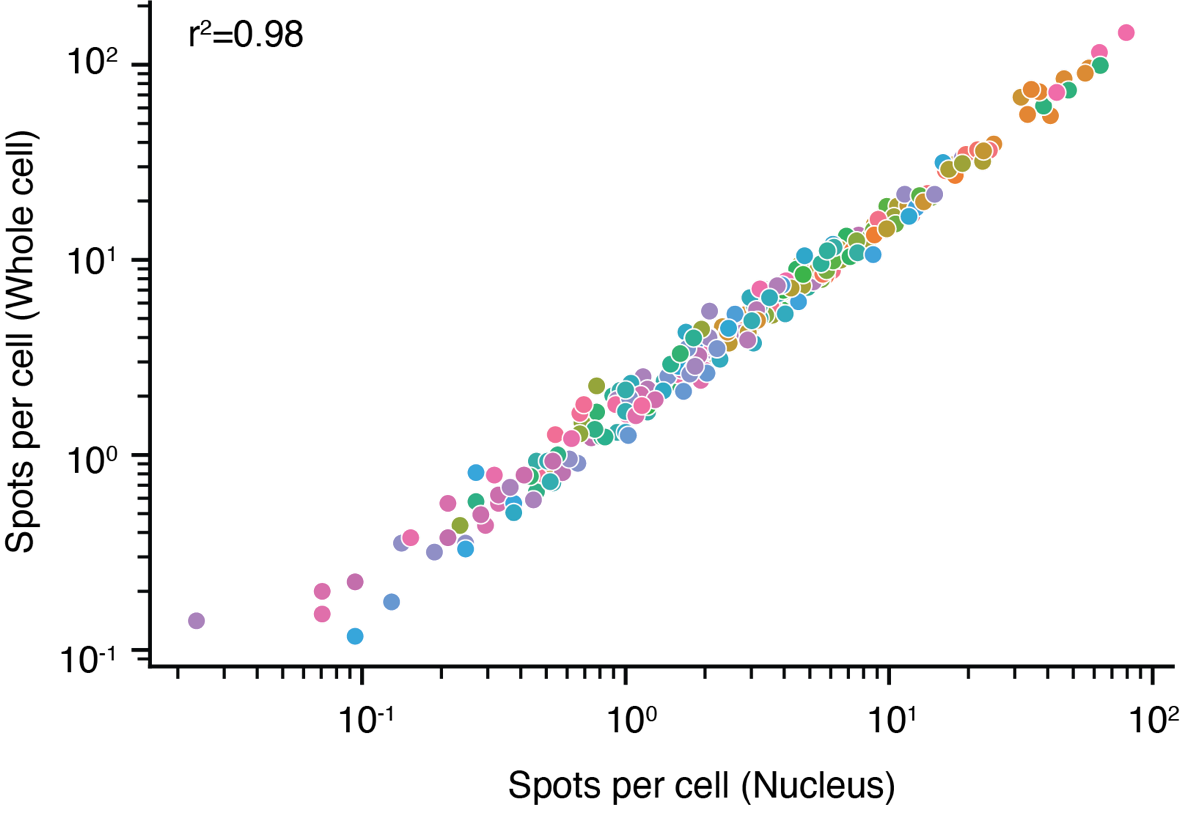

***SN2 Fig. 23****. Correlation calculated between the average spots per cell in the nucleus and the whole cell. Data includes 3 positions taken from 3 separate images.*

SN2 Fig. 23. Shows the correlation between average counts in cell segments including both the nuclei and cytoplasm versus just the nuclei. In our analysis we assign cytoplasmic spots to cells on the basis of a shortest path approach to the detected nuclei. We did not observe any major deviation of the data subset with is counterpart, indicating that the inferred cell wall position does not drastically alter the relative relationships between species by either including or excluding the cytoplasm in the segmentation on the basis of a shorted path preference.

### Supplementary Figures

Supplementary Fig. 1: Schematics of oligonucleotides labelled antibody and probes in ARTseq-FISH.

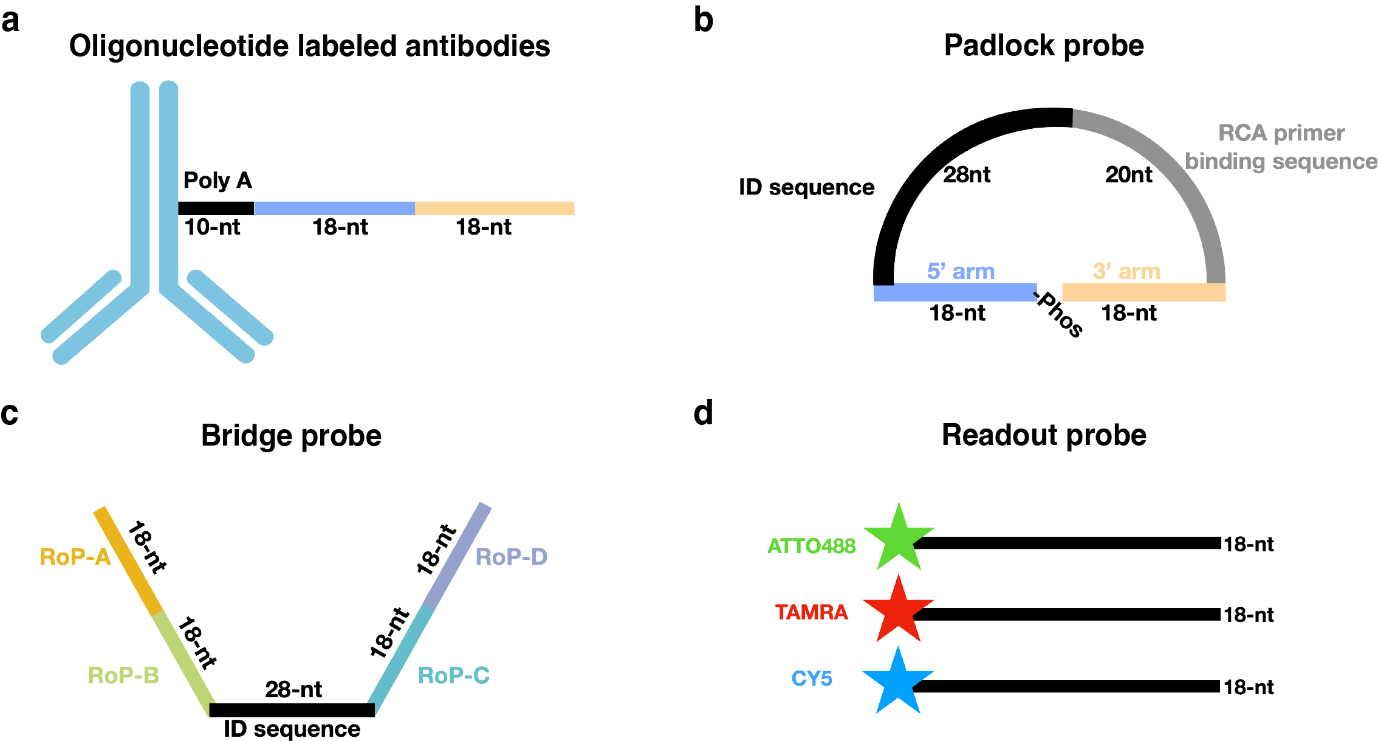

**Supplementary Fig. 1: a,** The oligonucleotide labelled to antibodies are designed as 10 nt polyadenylate and two 18 nt sequences (purple and light-yellow lines) complementary to unique padlock probes. **b,** PLPs consist of two 18 nt sequences (purple and light-yellow lines) recognizing the oligonucleotides of protein targets or cellular mRNAs, a 28 nt ID sequence (black line) for corresponding to unique bridge probes, and a 20 nt RCA primer binding sequence. **c,** BrPs are composed of a 28 nt ID sequence (black line) identical to the ID sequence of padlock probes, four 18nt sequences for different readout probes binding. **d,** RoPs are an 18 nt single strand DNA conjugated with three different fluorophores (ATTO488, TAMRA and CY5).

Supplementary Fig. 2: Generation of the ARTseq-FISH.

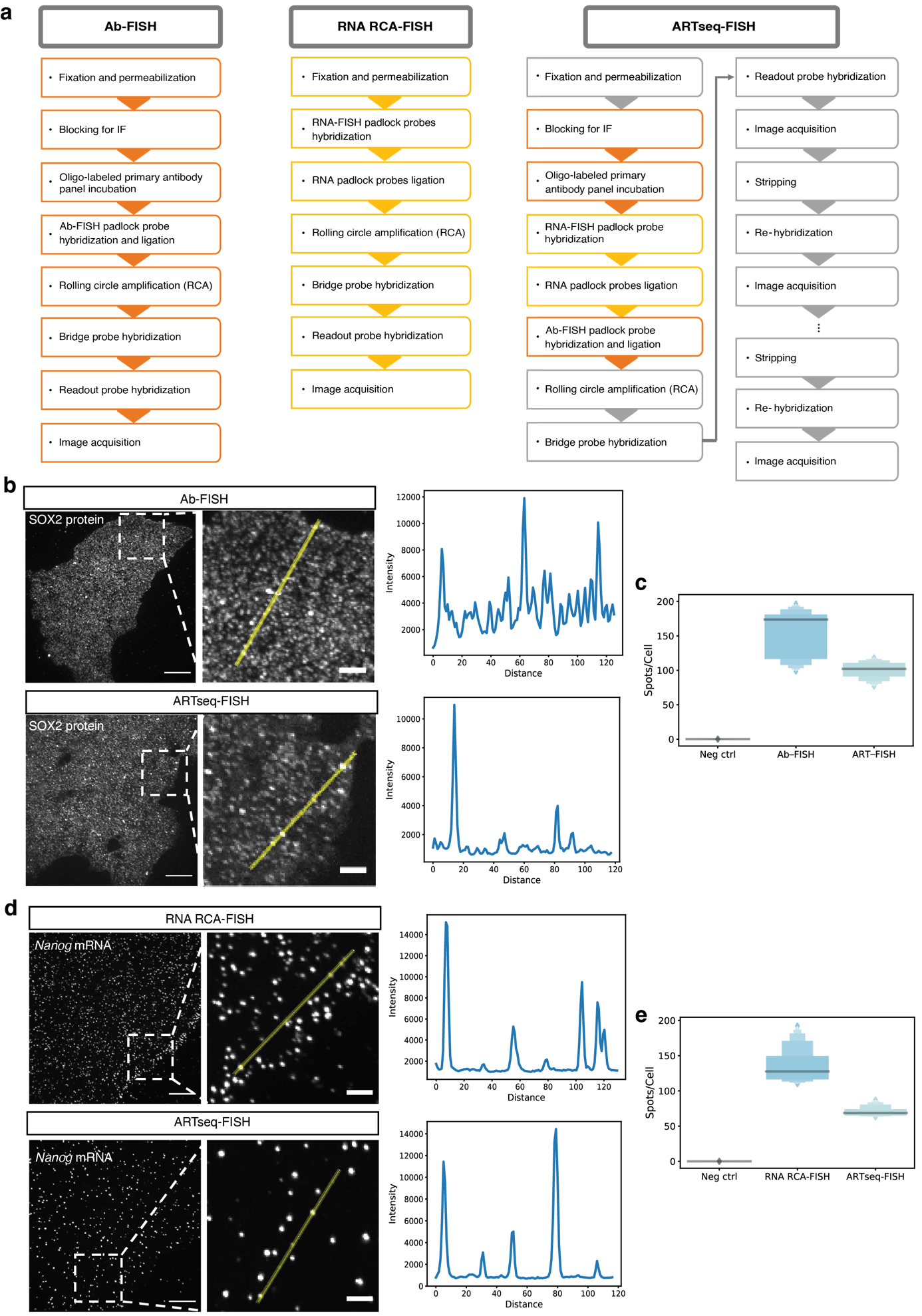

**Supplementary Fig. 2:** **a**, Flow chart of the experimental procedures showing protein and mRNA detection separately and simultaneously with ARTseq-FISH. The protocols of Ab-FISH and RNA RCA-FISH are demonstrated in orange and yellow boxes respectively. The simultaneous protein and mRNA detection protocol, i.e., ARTseq-FISH, is illustrated by orange, yellow and grey boxes, which show the Ab-FISH specific-, RNA RCA-FISH specific, and combined steps respectively. **b**, Representative max intensity and Z projection images of SOX2 protein detection by Ab-FISH and ART-FISH. Line plots showed the intensity of the selected spots in zoomed images. Scale bar, 20 µm, 5 µm. **c**, Quantification of spots per cell of SOX2 protein detected by Ab-FISH and ARTseq-FISH. Neg ctrls were performed by ARTseq-FISH protocols without PLPs added. (Neg ctrl n=362, Ab-FISH n=256, ARTseq-FISH n=291). Line in the boxenplot denoting the median. **d**, Representative max intensity Z projection images of *Nanog* mRNA detection by RNA RCA-FISH and ART-FISH. Line plots show the intensity of the selected spots in zoomed images. Scale bar, 20 µm, 5 µm. **e**, Quantification of spots per cell of *Nanog* mRNA detected by RNA-FISH and ARTseq-FISH. Neg ctrl were performed by ARTseq-FISH protocols without PLPs added. (Neg ctrl n=360, RNA RCA-FISH n=1193, ARTseq-FISH n=454). Line in the boxenplot denoting the median.

Supplementary Fig. 3: Negative controls in ARTseq-FISH.

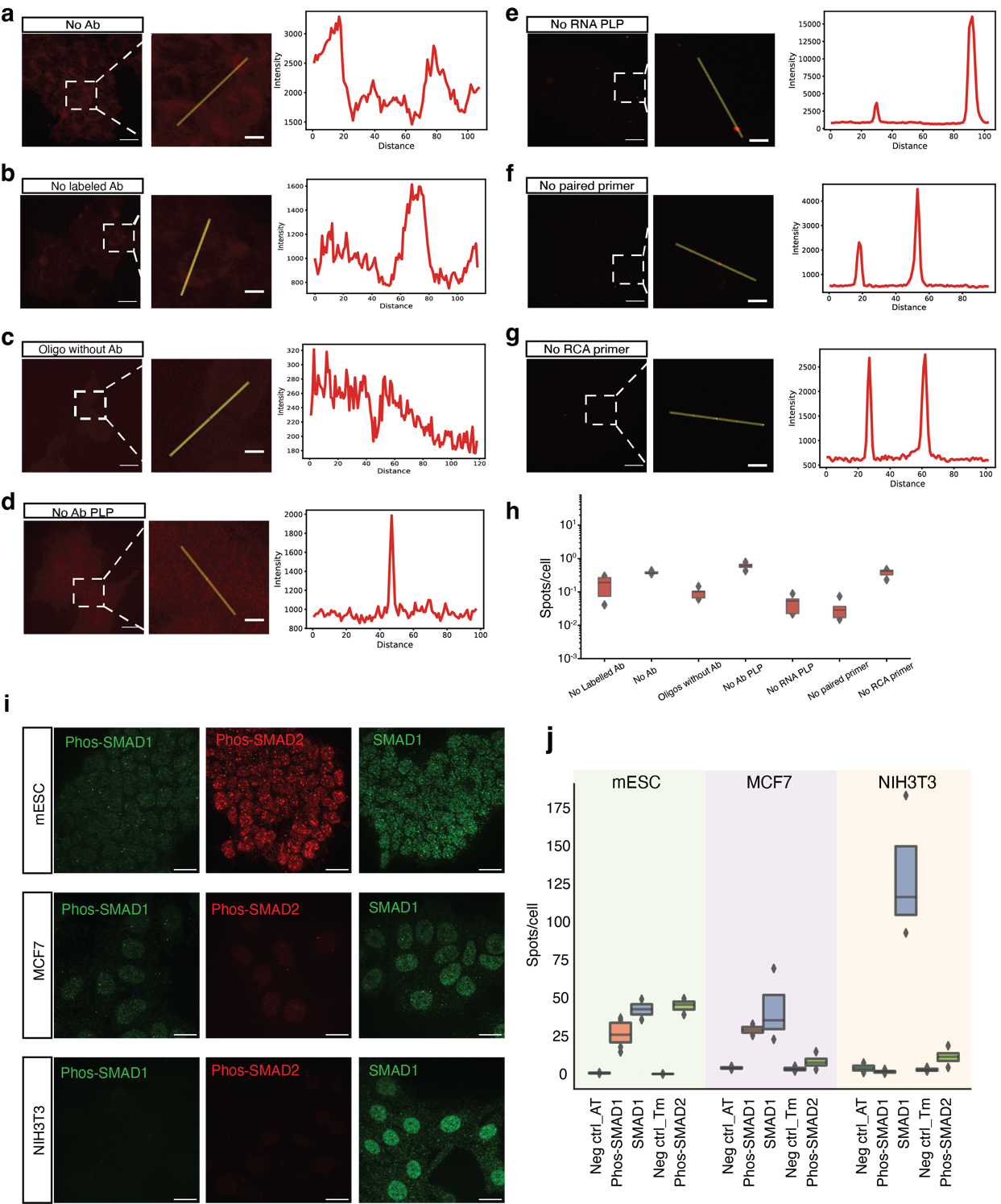

**Supplementary Fig. 3:** **a-g,** Representative max intensity and Z projection images in TAMRA channel from each negative control were shown. Neg ctrl were assigned as followed **a,** No antibodies (Abs) were incubated; **b,** Abs without oligonucleotides labelled were incubated; **c,** Only oligonucleotides were added but no Abs; **d,** No Abs PLP and **e,** no mRNA PLP were added; **f,** Scrambled (i.e., non-complementary) RCA primers were added; **g,** No RCA primers were added. Scale bar, 20 µm. **h,** Quantitative box plot of spots per cell in all the negative controls (No Ab n=195, No Labelled Ab n=198, Oligos without Ab n=310, No Ab PLP n=266, No mRNA PLP n=241, No paired primer n=324, No RCA primer n=206). Line in the boxenplot denoting the median. **i,** Example images of Phos-SMAD1, Phos-SMAD2, and SMAD1 proteins detected by ARTseq-FISH in mESC, MCF7, and NIH3T3 cells ^29, 30^. These data are showing the specificity of the ARTseq-FISH antibodies. **j,** Quantification of detected Phos-SMAD1, Phos-SMAD2, and SMAD1 proteins in mESC, MCF7, and NIH3T3 cells. Line in the boxenplot denoting the median.

Supplementary Fig. 4: Different methods of PLPs-based RNA FISH.

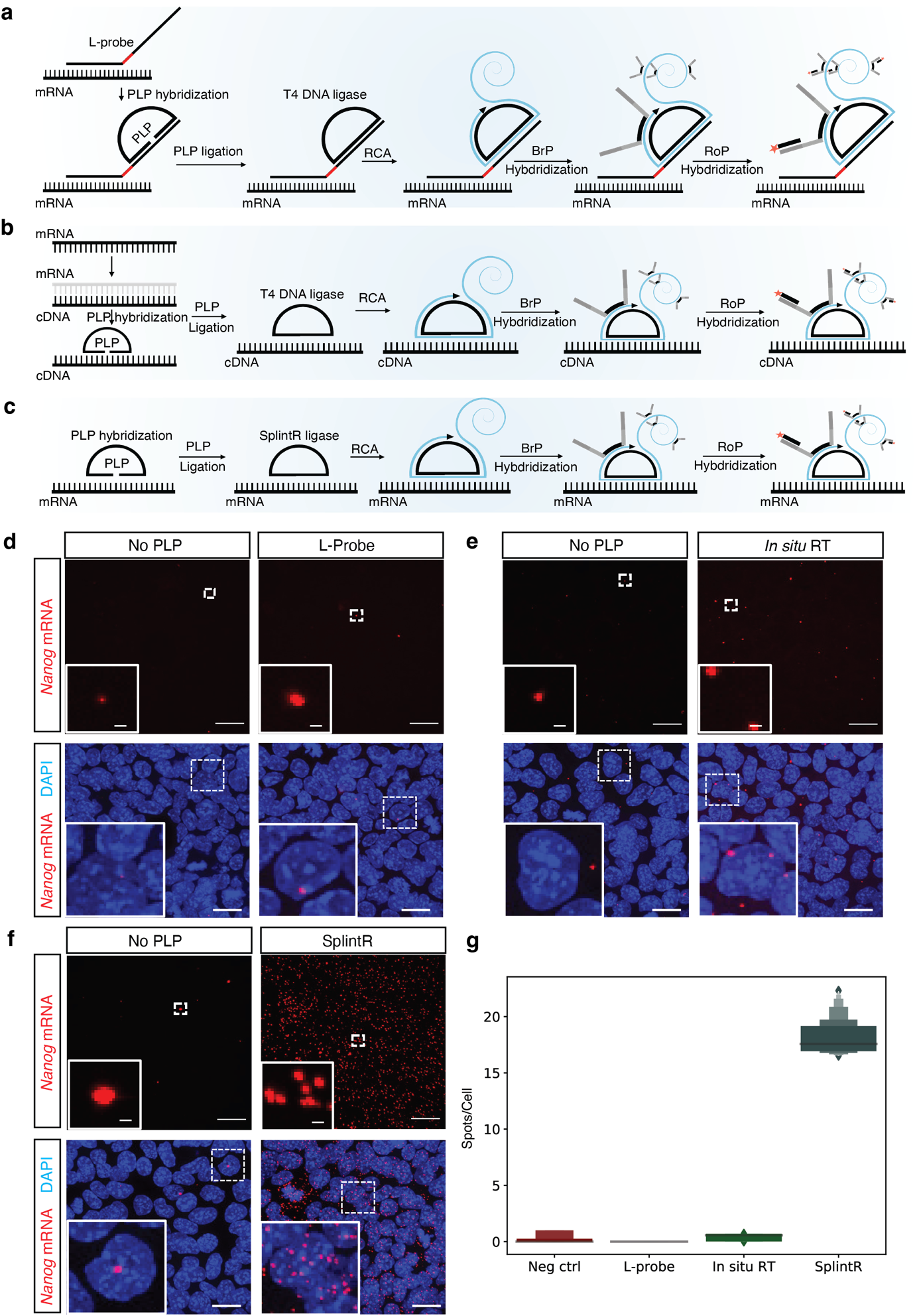

**Supplementary Fig. 4: a-c,** Schematics of different methods of PLPs-based RNA RCA-FISH. **d-f,** Representative max intensity and Z projection images of each method. Negative controls (Neg ctrl) are samples without PLPs. Zoomed images showed the representative spots. DAPI (blue) is nuclei staining. Scale bar, 20 µm and 1 µm. **g,** Quantification of spots per cell using these three different mRNA detection methods. (Neg ctrl n=656, L-probe n=109, Neg ctrl n=656, In situ RT n=376, Neg ctrl n=656, SplintR n=778). Line in the boxenplot denoting the median.

Supplementary Fig. 5: Optimizing the PLPs hybridization and ligation of RNA RCA-FISH.

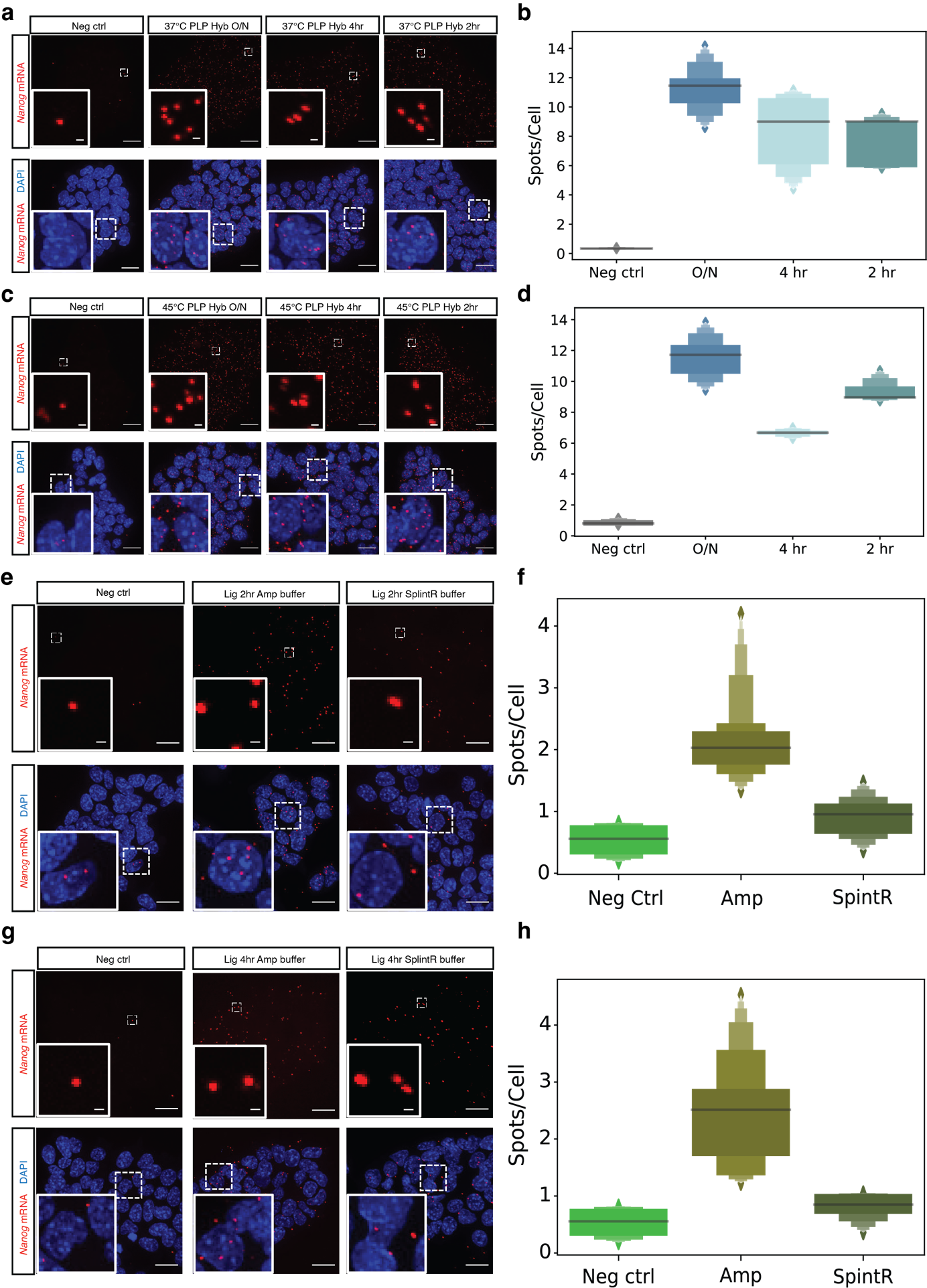

**Supplementary Fig. 5: a and c,** PLPs hybridization at 37°C or 45°C for 2 hours (hr), 4 hours (hr) and overnight. Representative max intensity and Z projection images of detected *Nanog* mRNA under different conditions. Zoomed images showed the representative spots. DAPI (blue) is nuclei staining. Scale bar, 20 µm and 1 µm. **b and d,** Quantification of *Nanog* mRNA spots per cell under different conditions of PLPs hybridization. (Neg ctrl n=536, 37°C O/N n=369, 37°C 4hr n=271, 37°C 2hr n= 380; Neg ctrl n=209, 45°C O/N n=339, 45°C 4hr n=535, 45°C 2hr n= 476). Line in the boxenplot denoting the median.**e and g,** PLPs ligation for 2 hr and 4 hr with Ampligase buffer or SplintR buffer. Representative max intensity and Z projection images of detected *Nanog* mRNA under different conditions. Zoomed images showed the representative spots. DAPI (blue) is nuclei staining. Scale bar, 20 µm and 1 µm. **f and h,** Quantification of *Nanog* mRNA spots per cell under different conditions of PLPs ligation. (Neg ctrl n=181, 2hr Ampligase buffer n=614, 2hr SpintR buffer n=,656; Neg ctrl n=181, 4hr Ampligase buffer n=398, 4hr SpintR buffer n=670). Line in the boxenplot denoting the median.

Supplementary Fig. 6: Optimizing the PLPs ligation condition of RNA RCA-FISH.

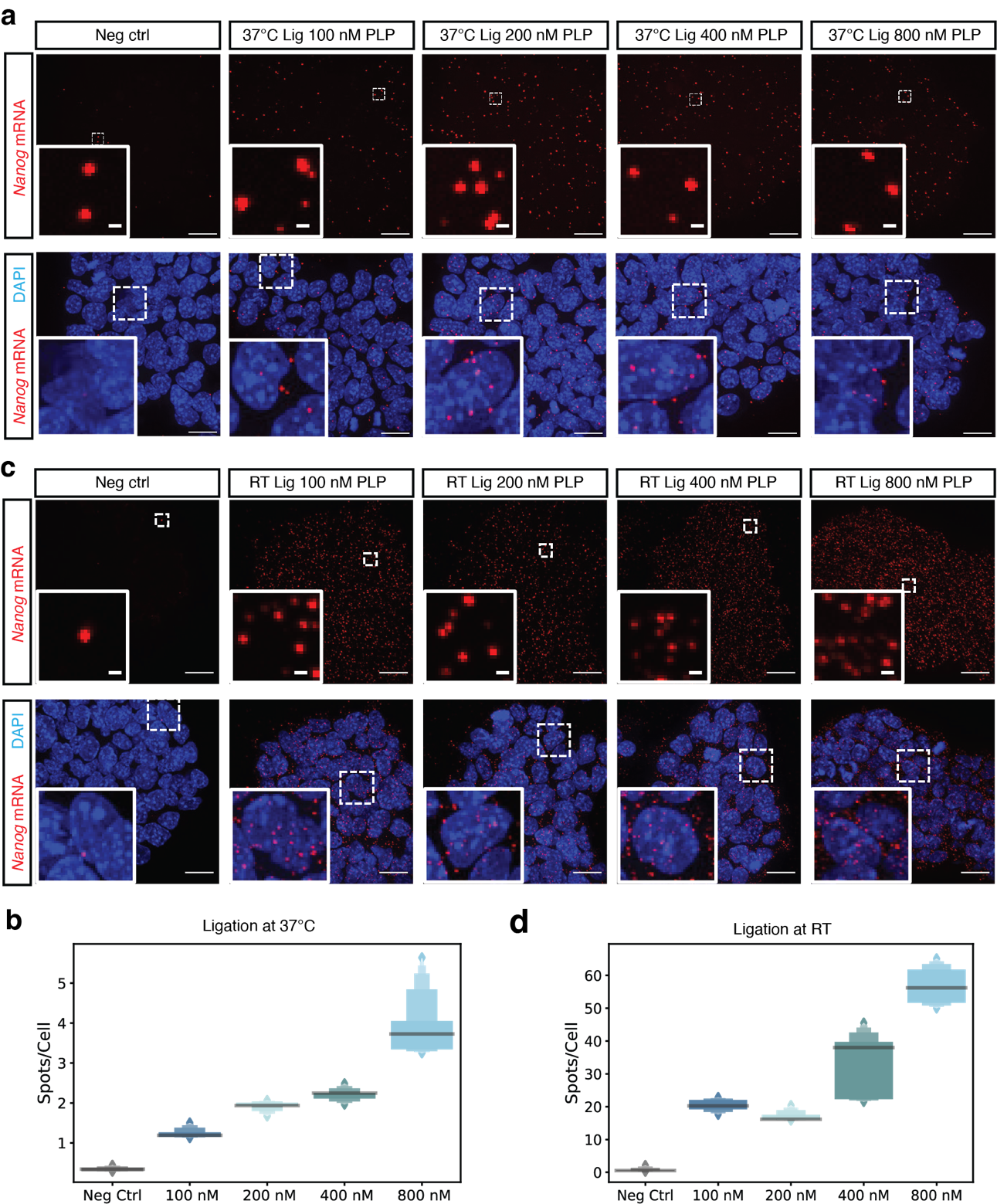

**Supplementary Fig. 6: a and c,** Representative max intensity and Z projection images of *Nanog* mRNA detection with different concentrations (100 nM, 200 nM, 400 nM and 800 nM) of PLPs ligation at 37°C or room temperature. Zoomed images showed the representative spots. DAPI (blue) is nuclei staining. Scale bar 20 µm, 1 µm. **b and d,** Quantification of spots per cell of *Nanog* mRNA detected under different PLP concentration. (Neg ctrl n=543, 37°C 100 nM n=507, 37°C 200 nM n=546, 37°C 400 nM n=539, 37°C 800 nM n=419; Neg ctrl n=433, RT 100 nM n=547, RT 200 nM n=437, RT 400 nM n=531, RT 800 nM n=499;). Line in the boxenplot denoting the median.

Supplementary Fig. 7: Optimizing the PLPs concentration of Ab-FISH.

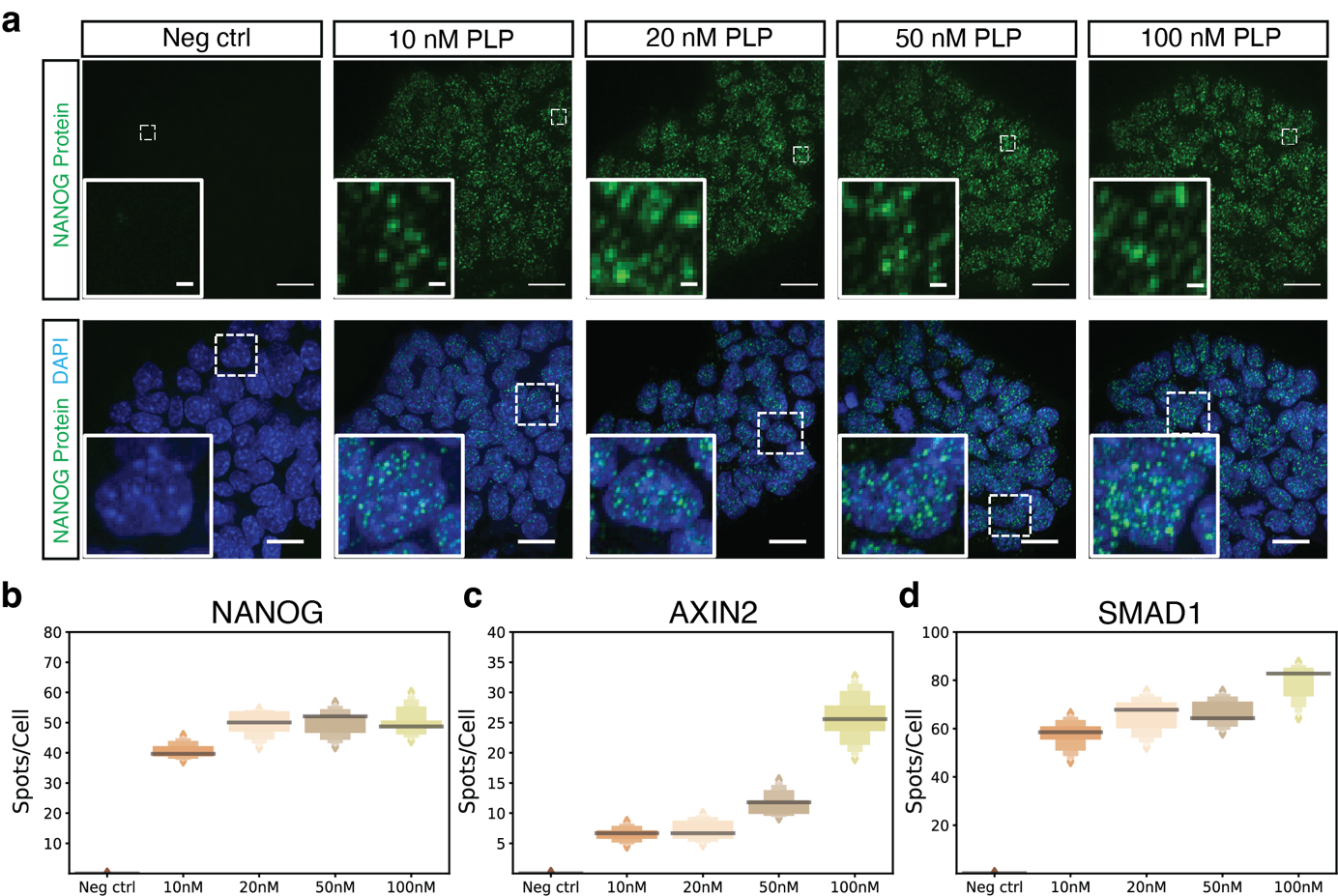

**Supplementary Fig. 7: a,** Three proteins were detected with different concentrations (10 nM, 20 nM, 50 nM and 100 nM) of PLPs simultaneously in three different channels. Representative max intensity and Z projection images of detected NANOG protein with different concentrations of PLPs. Zoomed images showed the representative spots. DAPI (blue) is nuclei staining. Scale bar 20 µm, 1 µm. **b-d,** Quantification of spots per cell of NANOG, AXIN2 and SMAD1 protein. (Neg ctrl n=424, 10 nM n=330, 20 nM n=319, 50 nM n=310, 100 nM n=308). Line in the boxenplot denoting the median.

Supplementary Fig. 8: Optimizing the RCA condition.

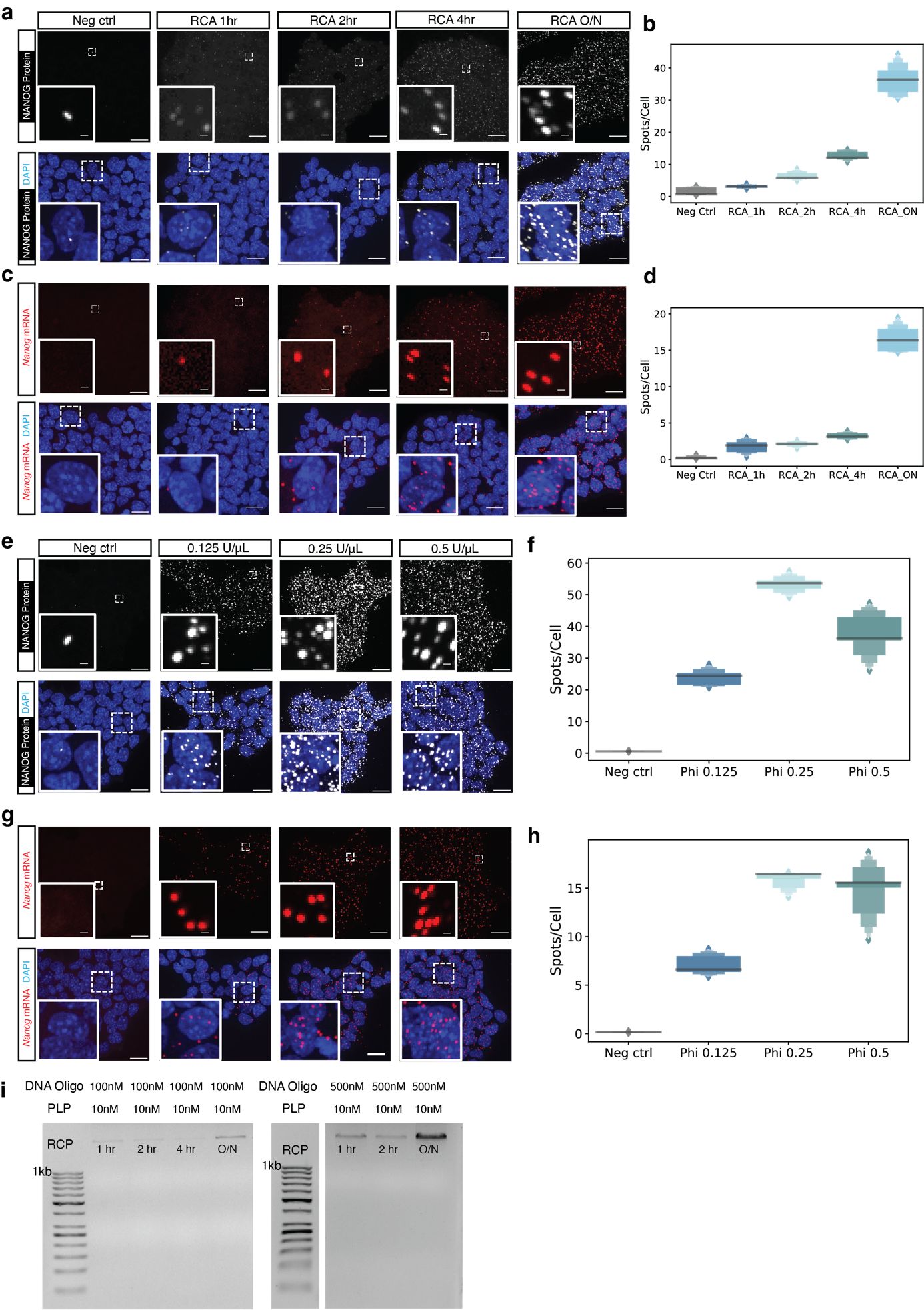

**Supplementary Fig. 8:** **a and c,** ARTseq-FISH with RCA incubation time of 1 hr, 2 hr, 4 hr and overnight. Representative max intensity and Z projection images of detected *Nanog* mRNA and NANOG protein with different RCA time. Zoomed images show the representative spots. DAPI (blue) is nuclei staining. Scale bar 20 µm, 1 µm. **b and d,** Quantification of spots per cell of *Nanog* mRNA and NANOG protein. (Neg ctrl n=424, RCA 1hr n=638, RCA 2hr n=401, RCA 4hr n=466, RCA ON n=328). Line in the boxenplot denoting the median. **e and g,** RCA with different concentration of phi29 polymerase (0.125U/μL, 0.25U/μL and 0.5U/μL). Representative max intensity and Z projection images of detected *Nanog* mRNA and NANOG protein under different phi29 concentrations. Zoomed images showed the representative spots. DAPI (blue) is nuclei staining. Scale bar, 20 µm and 1 µm. **f and h,** Quantification of *Nanog* mRNA and NANOG protein spots per cell under different conditions of phi29 concentrations. (Neg ctrl n=148, phi29 0.125U/μL n=334, phi29 0.25U/μL n=290, phi29 0.5U/μL n=314). Line in the boxenplot denoting the median. **i,** RCA with different concentration of oligonucleotides (100 nM and 500 nM). Agarose gel (0.8% agarose) analysis of the rolling circle amplified products (RCPs) under different incubation time (1 hr, 2 hr, 4 hr and overnight).

Supplementary Fig. 9: Optimizing the concentration of BrPs and RoPs.

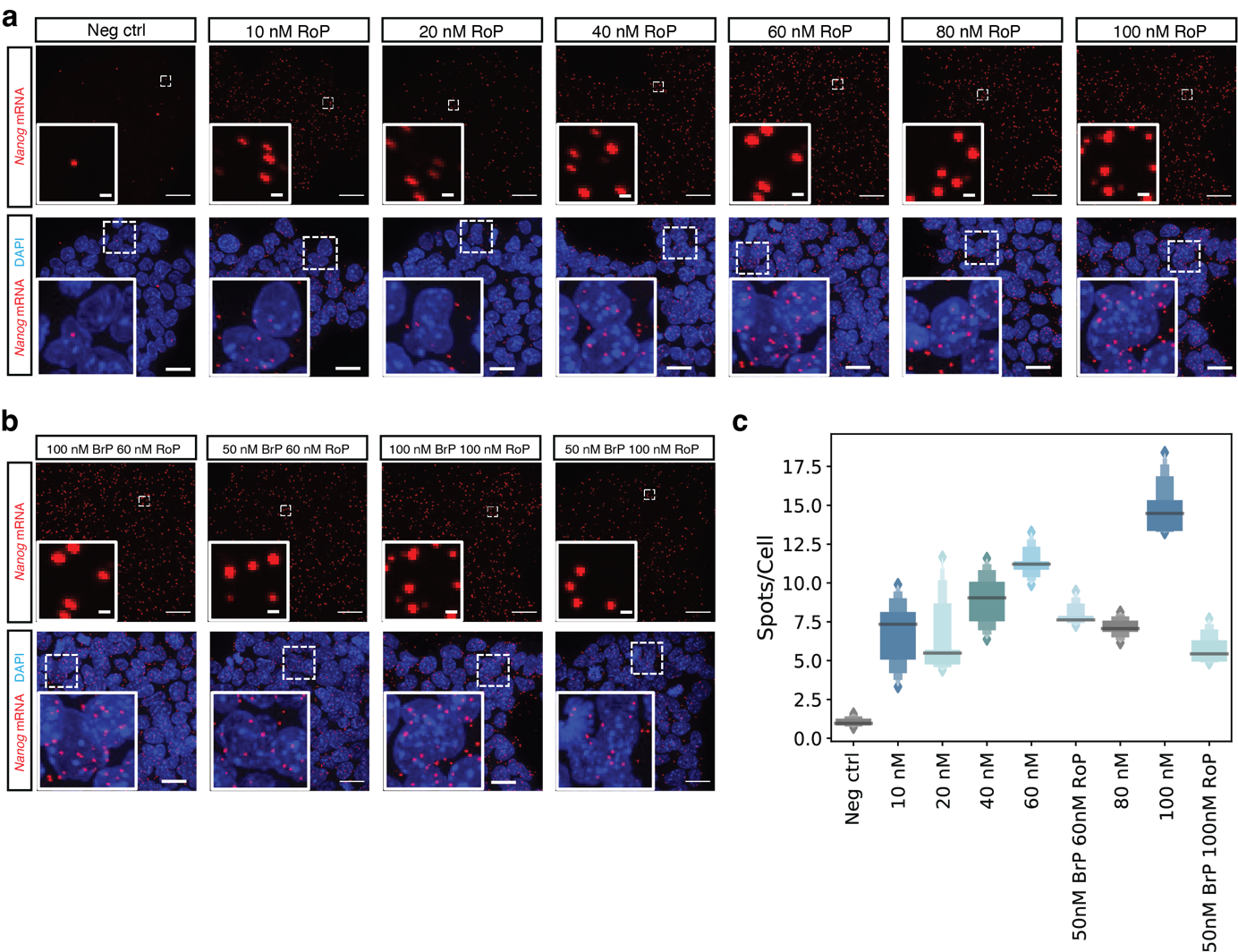

**Supplementary Fig. 9:** **a,** ARTseq-FISH detection of *Nanog* mRNA with 100 nM BrP and different concentrations (10 nM, 20 nM, 40 nM, 60 nM, 80 nM, 100 nM) of RoP. **b,** ARTseq-FISH detection of *Nanog* mRNA with 50 nM BrP and 60 nM or 100 nM of RoP. Representative max intensity and Z projection images of detected *Nanog* mRNA under different conditions. Zoomed images showed the representative spots. DAPI (blue) is nuclei staining. Scale bar 20 µm, 1 µm. **c,** Quantification of *Nanog* mRNA spots per cell under different conditions of BrP and RoP concentrations. (Neg ctrl n=182, 10 nM RoP n=391, 20 nM RoP n=341, 40 nM RoP n=357, 60 nM RoP n=442, 50 nM BrP 60 nM RoP n=435, 80 nM RoP n=291, 100 nM RoP n=308, 50 nM BrP 100 nM RoP n=329). Line in the boxenplot denoting the median.

Supplementary Fig. 10: Optimizing mRNA PLPs selection and concentration.

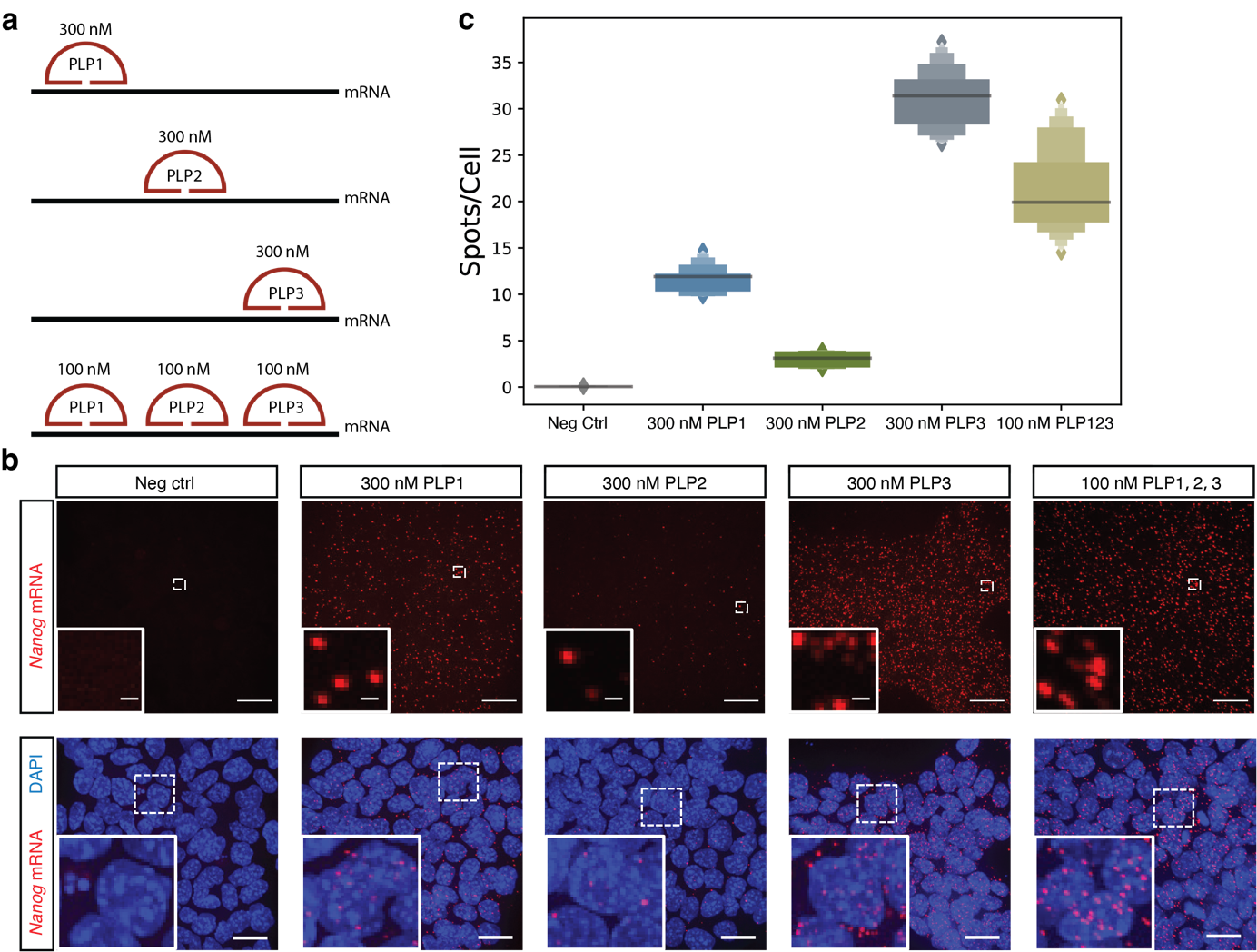

**Supplementary Fig. 10:** **a,** Schematics of targeting *Nanog* mRNA in three different selected regions of the exons. **b,** Representative max intensity and Z projection images of *Nanog* mRNA detection by different PLPs targeting different regions of exons. Zoomed images showed the representative spots. DAPI (blue) is nuclei staining. Scale bar, 20 µm and 1 µm. **c,** Quantification of *Nanog* mRNA as spots per cell under different conditions of PLPs targeting region. (Neg ctrl n=210, 300 nM PLP1 n=174, 300 nM PLP2 n=230, 300 nM PLP3 n=214, 100 nM PLP1, 2, 3 n=450). Line in the boxenplot denoting the median.

Supplementary Fig. 11: Post-fixation is necessary in sequential hybridization.

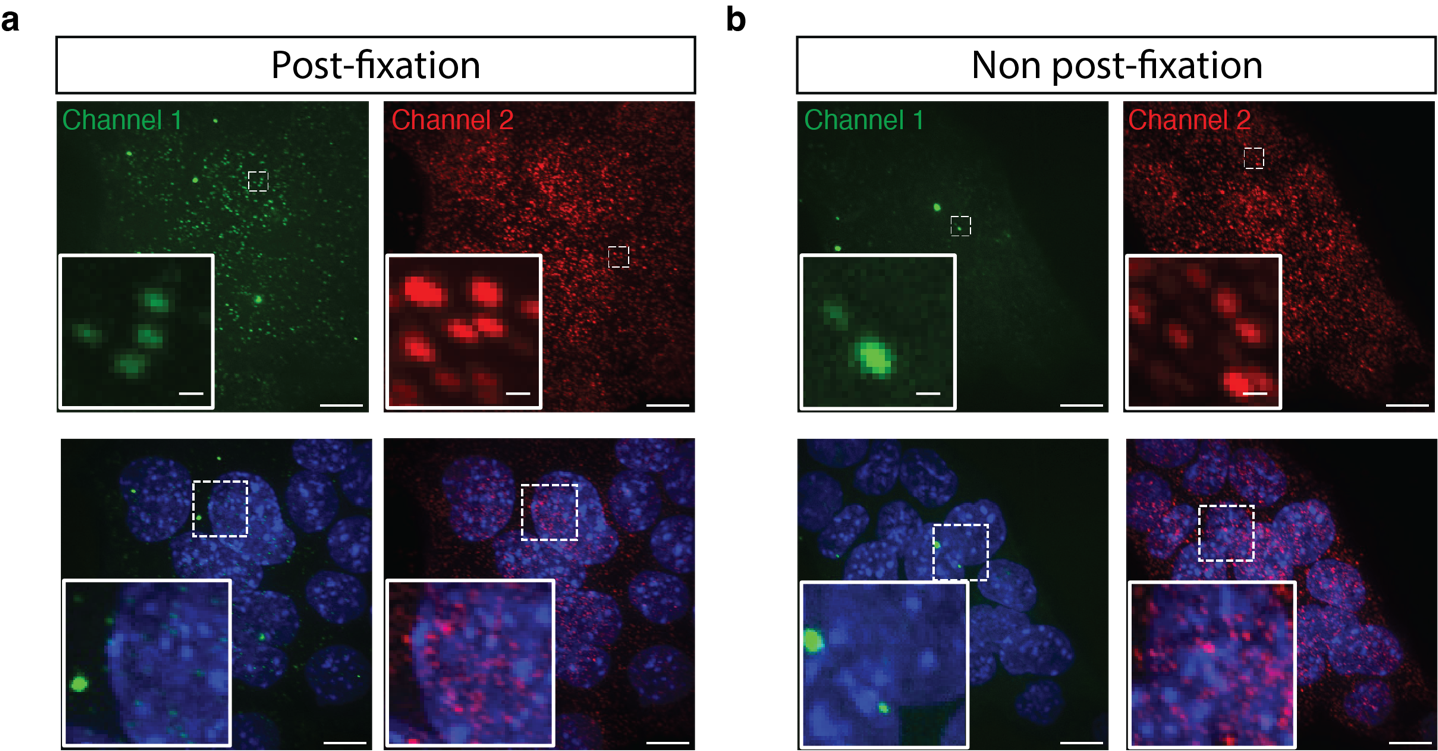

**Supplementary Fig. 11:** **a and b,** Second round hybridization results in two channels from samples with post-fixation **a,** and without post-fixation **b,** after RCA step. Representative max intensity and Z projection images of detected targets including mRNAs and proteins by ARTseq-FISH; Zoomed images show representative spots. DAPI (blue) is nuclei staining. Scale bar 20 µm, 1 µm.

Supplementary Fig. 12: Optimizing antibody dilution in ARTseq-FISH.

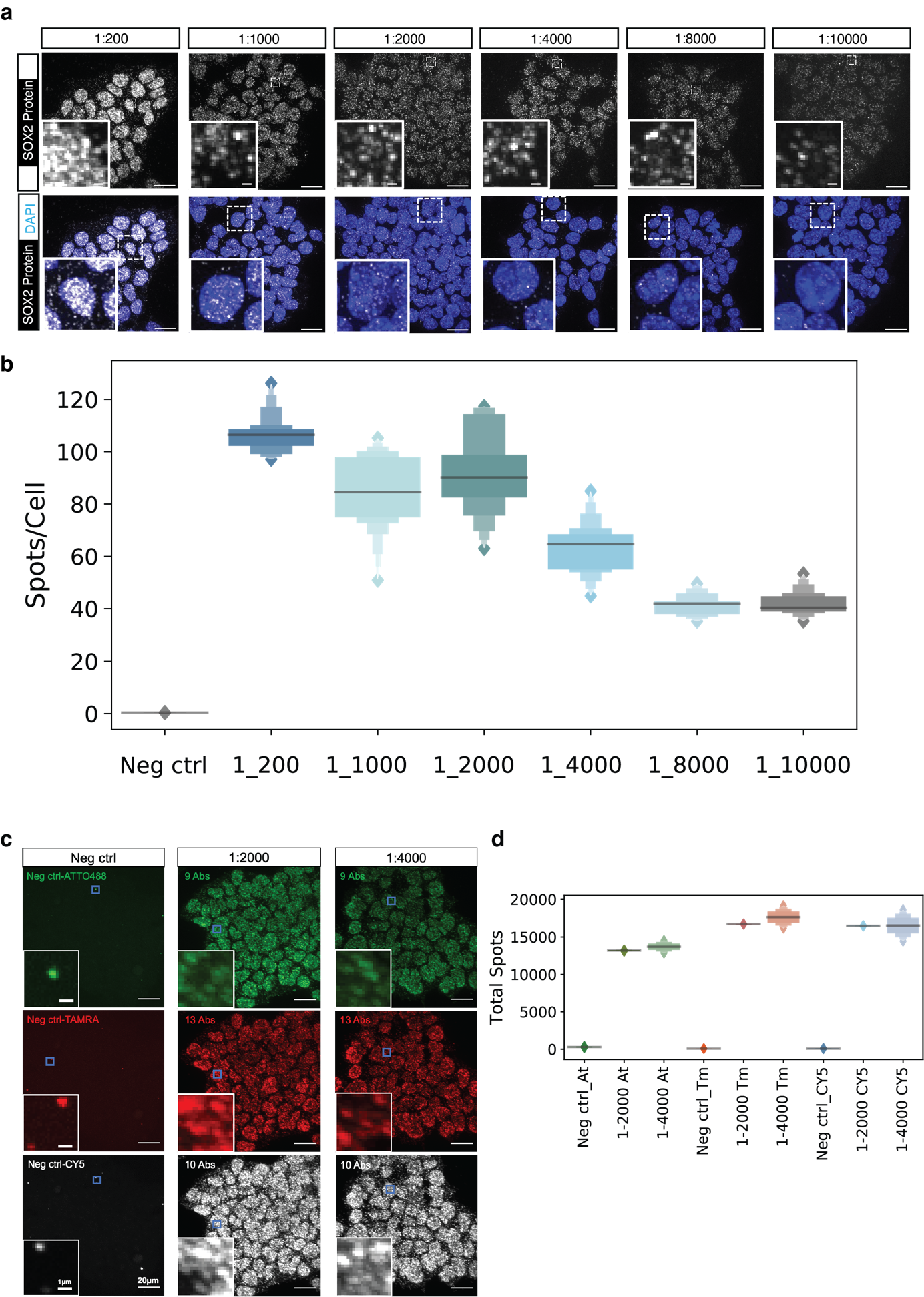

**Supplementary Fig. 12:** **a,** ARTseq-FISH detecting SOX2 protein with different antibody dilution ratios. Representative max intensity and Z projection images of SOX2 protein detected by ARTseq-FISH with different antibody dilution ratio; Zoomed images show representative spots. DAPI (blue) is nuclei staining. Scale bar 20 µm, 1 µm. **b,** Quantification of spots per cell of SOX2 protein. (Neg ctrl n=491, 1-200 n=689, 1-1000 n=1332, 1-2000 n=690, 1-4000 n=784, 1-8000 n=913, 1-10000 n=782). Line in the boxenplot denoting the median. **c,** ARTseq-FISH detecting 32 proteins with different antibody dilution ratios. Zoomed images show representative spots. Scale bar 20 µm, 1 µm. **d,** Quantification of average spots per cell of detected proteins in each channel. There are 9 proteins detected simultaneously in Channel 1 (Atto488), 13 proteins detected in Channel 2 (TAMRA), and 10 proteins detected in Channel 3 (CY5) (Neg ctrl n=132, 1-2000 n=138, 1-4000 n=103). Line in the boxenplot denoting the median.

Supplementary Fig. 13: Colocalization of SMAD1 protein by ARTseq-FISH.

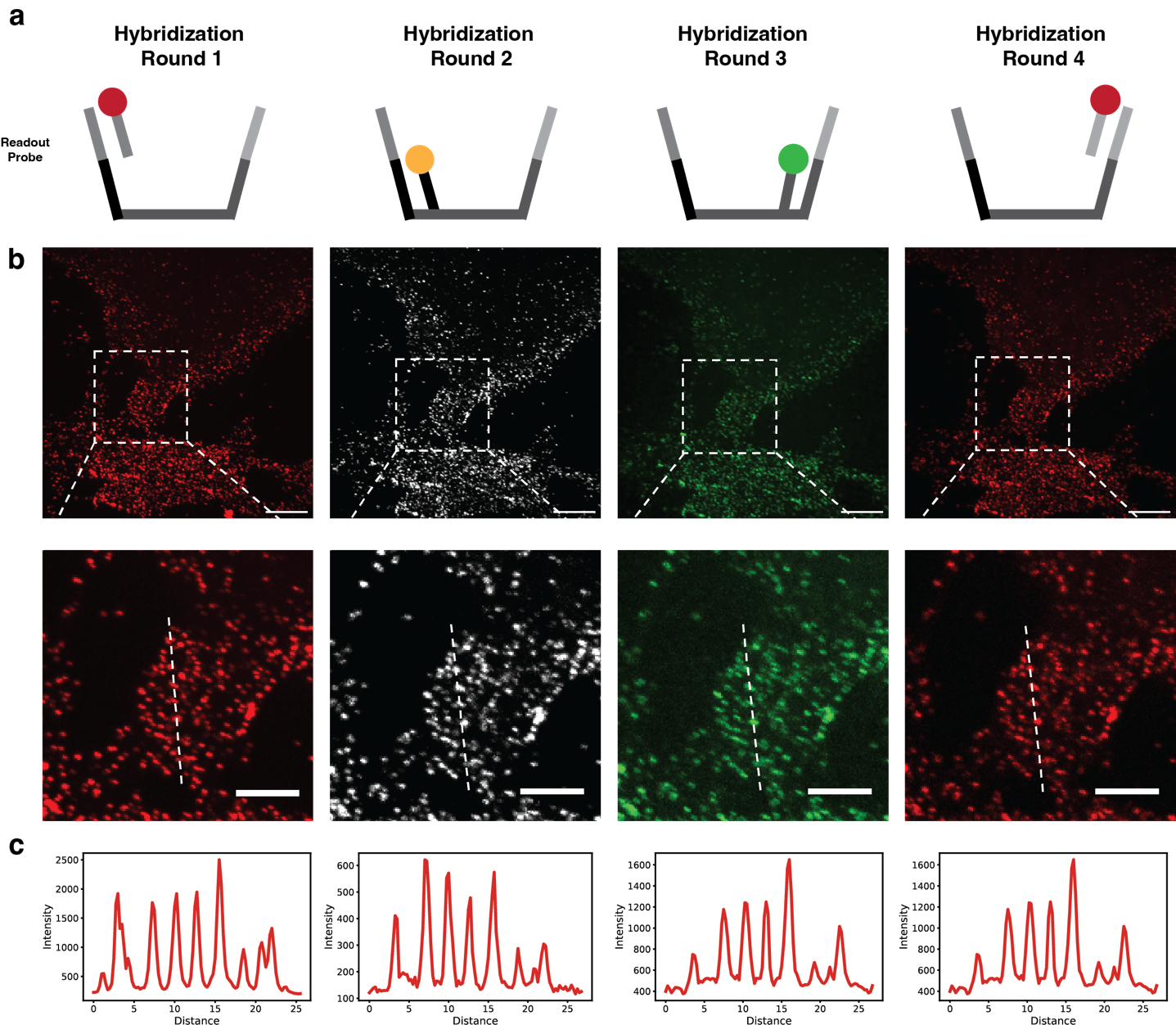

**Supplementary Fig. 13:** **a,** Schematics of sequentially detecting one target by ARTseq-FISH. **b,** Representative max intensity and Z projection images of SMAD1 protein detected by ARTseq-FISH in four hybridization rounds by four different readout probes; Zoomed image showed representative spots. Scale bar, 20 µm, 10 µm. **c,** Intensity of the spots in selected region (dashed line) from the zoomed images in **(b)**.

Supplementary Fig. 14: Sequential hybridization in ARTseq-FISH.

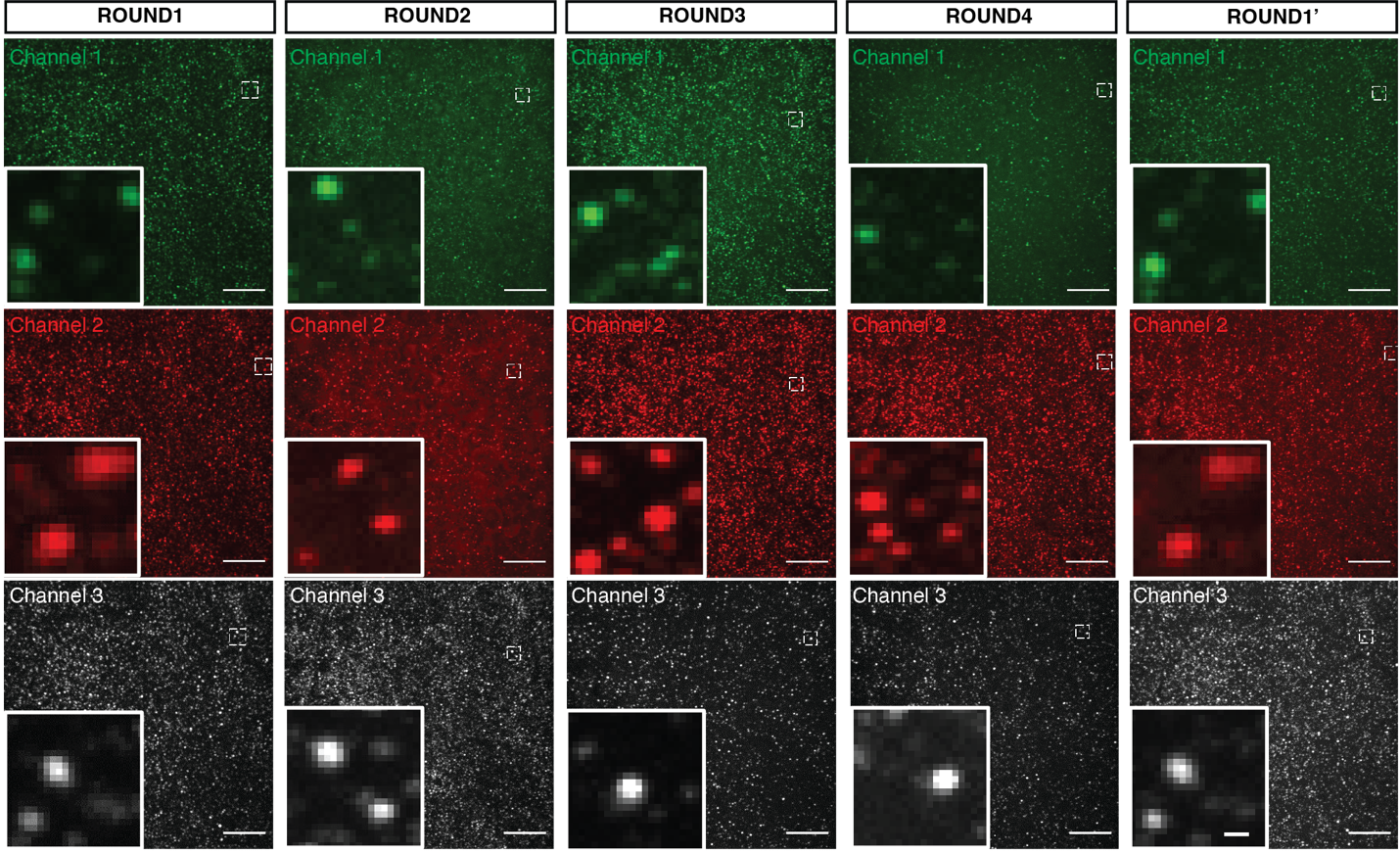

**Supplementary Fig. 14:** Representative max intensity and Z projection images of 64 targets including mRNAs and proteins detected by four hybridization rounds in three different channels. Round 1' is the fifth round with the same set of the readout probes used in round 1. Zoomed images showed the representative spots. Scale bar, 20 µm and 1 µm.

Supplementary Fig. 15: Colocalization of targets detected in round 1 and round 5.

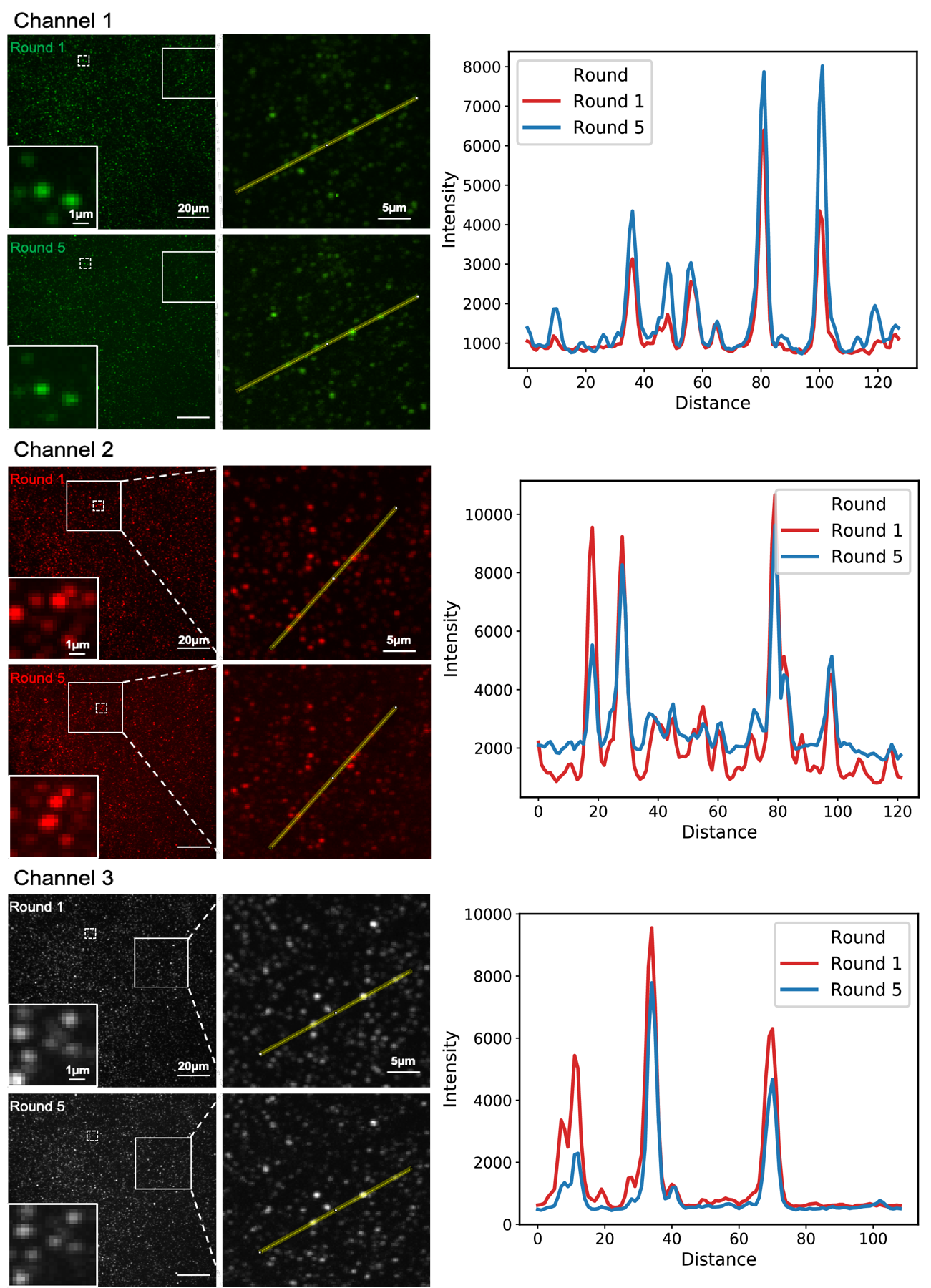

**Supplementary Fig. 15:** Same set of readout probes were used to detect the same targets in round 1 and round 5 (i.e., 1’). Representative max intensity and Z projection images of colocalized spots in two repeated hybridization rounds (round 1 and round 5). Zoomed images demonstrate representative colocalized spots. The line plot shows the intensity of selected spots demonstrating colocalization. Scale bar, 20 µm and 1 µm.

Supplementary Fig. 16: Stripping efficiency in ARTseq-FISH.

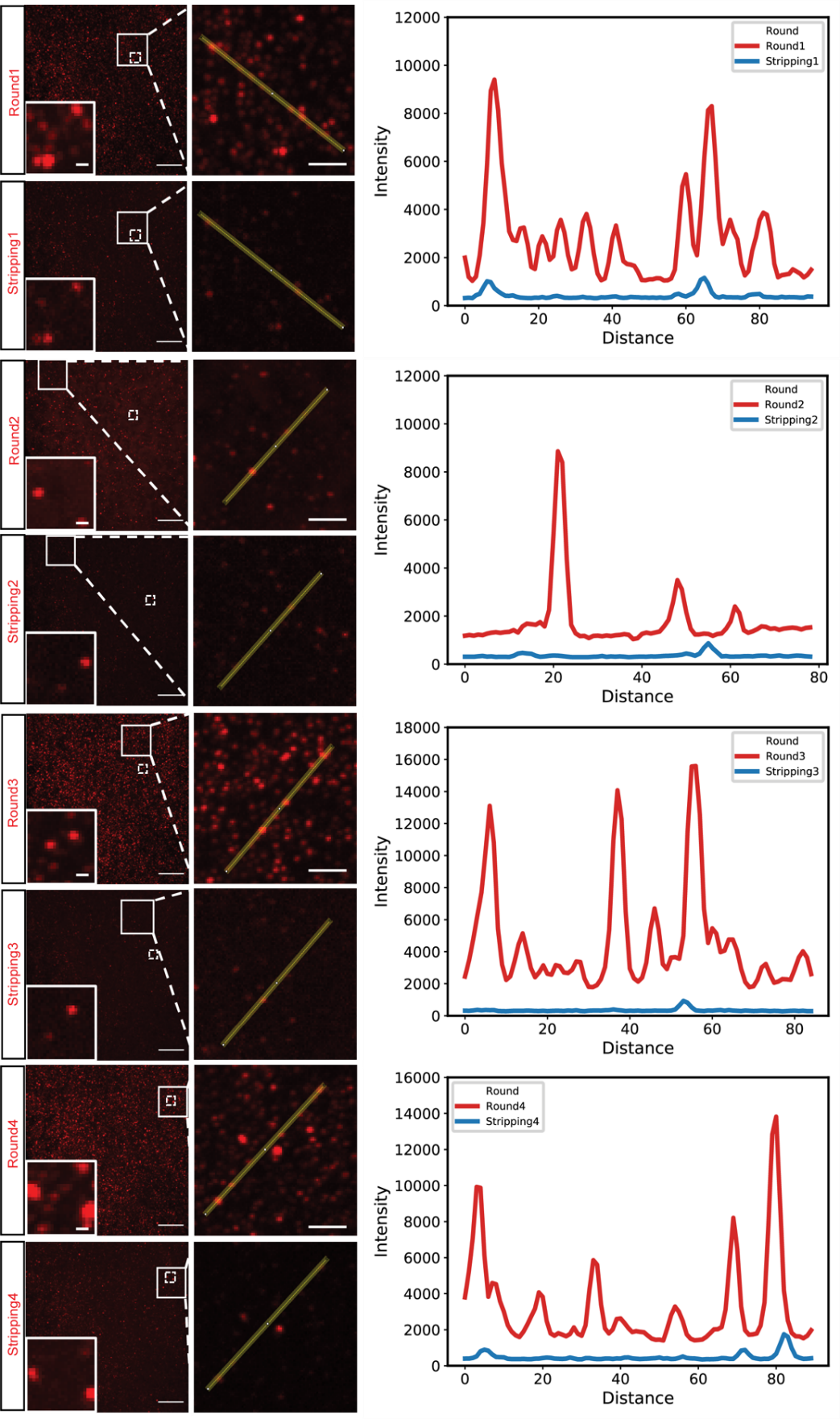

**Supplementary Fig. 16:** Representative max intensity and Z projection images of each hybridization round and after stripping off the readout probes of each round. Zoomed images demonstrate representative spots. The line plots show the intensity of the selected spots in zoomed images. The intensity of selected spots after stripping (blue line) dropped lower than the background of the selected region (red lines) in the images of hybridization. Scale bar, 20 µm, 5 µm and 1 µm.

Supplementary Fig. 17: Validation of protein targets by ARTseq-FISH.

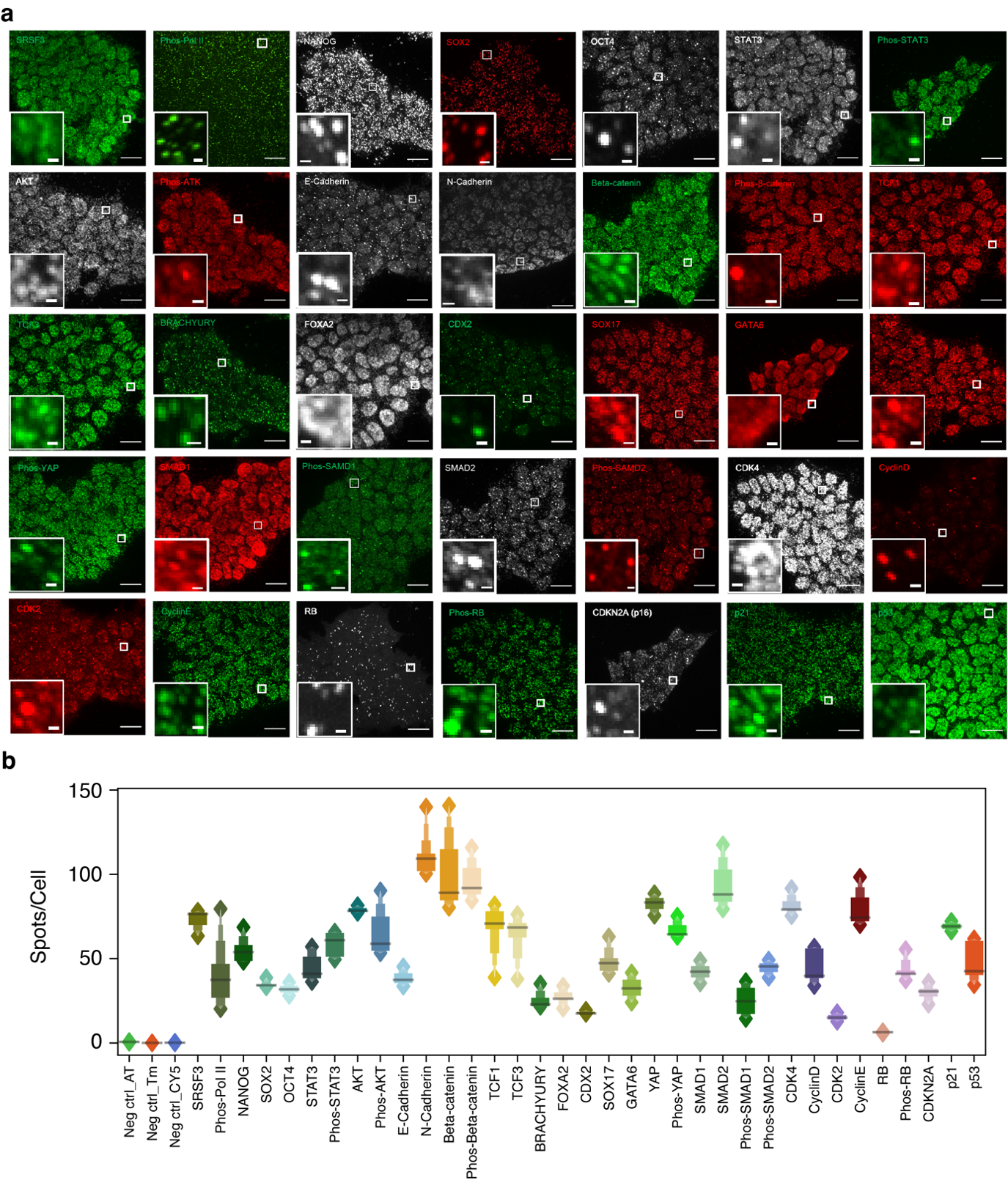

**Supplementary Fig. 17: a,** Representative max intensity and Z projection images of each protein target. Zoomed images show representative spots. Scale bar, 20 µm and 1 µm. **b,** Quantification of spots per cell for validated antibodies. Negative controls were incubated without antibodies. (Neg ctrl n=303, SOX2 and RB n=365, Phos-AKT, BRACHYURY and NANOG n=477, SRSF3, Phos-SMAD2, and STAT3 n=265, Cyclin E, Phos-Beta-catenin and OCT4 n=223, CDKN2A, GATA6 and Phos-STAT3 n=222, Beta-catenin and SMAD2 n=226, Phos-YAP and SMAD1 n=319, P21, CDK2 and E-Cadherin n=232, N-Cadherin n=1360, CDX2, YAP and AKT n=212, TCF3, TCF7 and FOXA2 n=314; Phos-RB, SOX17 and CDK4 n=366, Cyclin D and p53 n=395, phos-SMAD1 n=652, Phos-Pol II n=1000). Line in the boxenplot denoting the median.

Supplementary Fig. 18: Validation of mRNA targets by ARTseq-FISH.

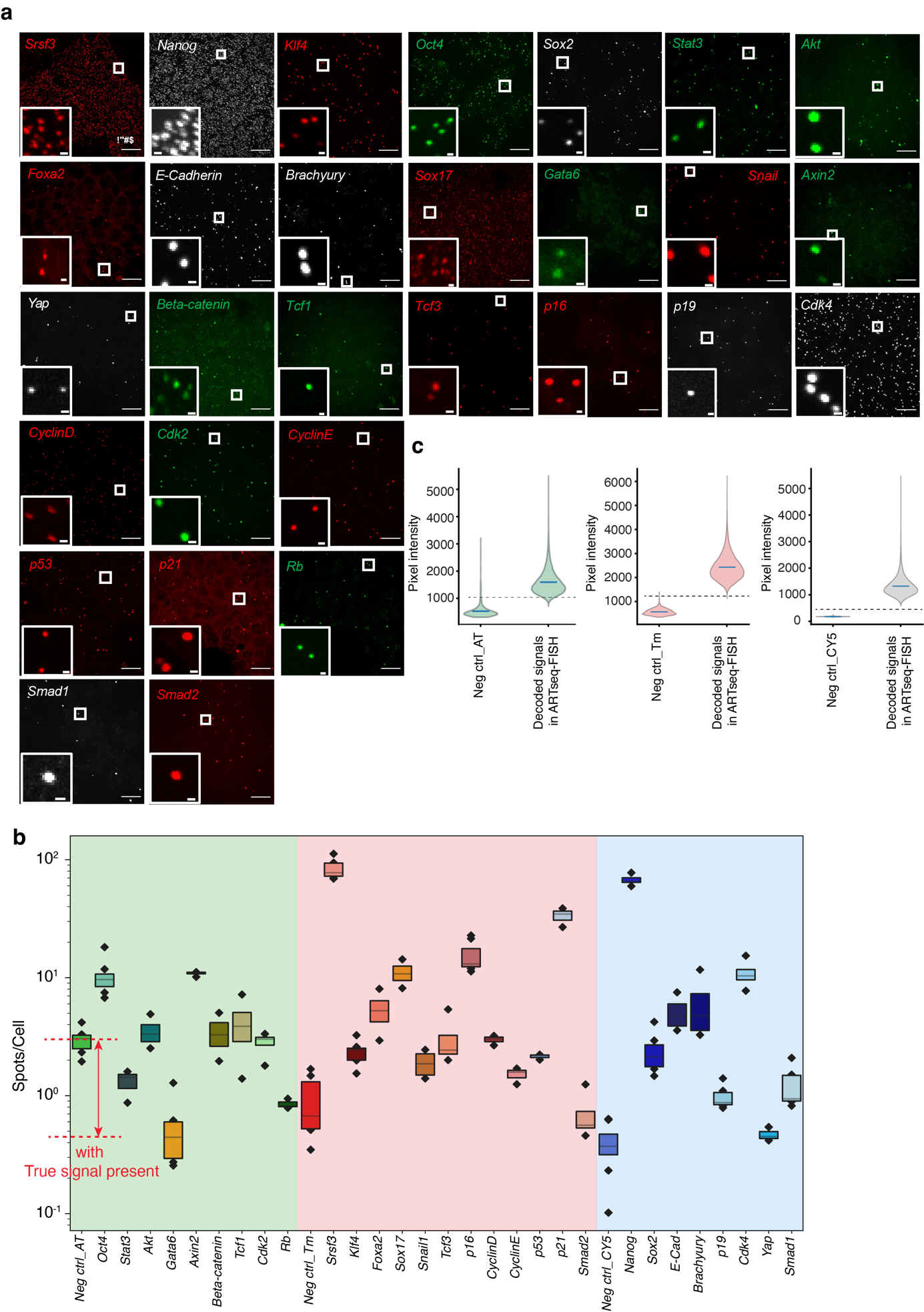

**Supplementary Fig. 18:** **a,** Representative max intensity and Z projection images of each mRNA target. Zoomed images show representative spots. Scale bar, 20 µm and 1 µm. **b,** Quantification of spots per cell for targeted mRNAs. Negative controls were performed without PLPs. (Neg ctrl n=579, *Akt* and *Smad2* n=455, *Tcf1*, *Tcf3* and *Brachyury* n=503, *Foxa2*, *E-cadherin* n=470, *Cdk2*, *Cdk*4 and *Snail* n=482, *Axin2* and *CyclinD* n=454, *Gata6*, *p16* and *Smad1* n=466, *Klf4* and *p19* n=488, *Oct4*, *Nanog* and *Sox*2 n=408, *p53* n=180, *Sox*17 and *Beta-catenin* n=241, *Stat3* and *CyclinE* n=200, *Yap*, *Rb* and *p21* n=233). Line in the boxenplot denoting the median. **c,** Pixel intensity of spots identified in the negative control compared to decoded signals in ARTseq-FISH.

Supplementary Fig. 19: Comparison of ARTseq-FISH and IF in detection of proteins.

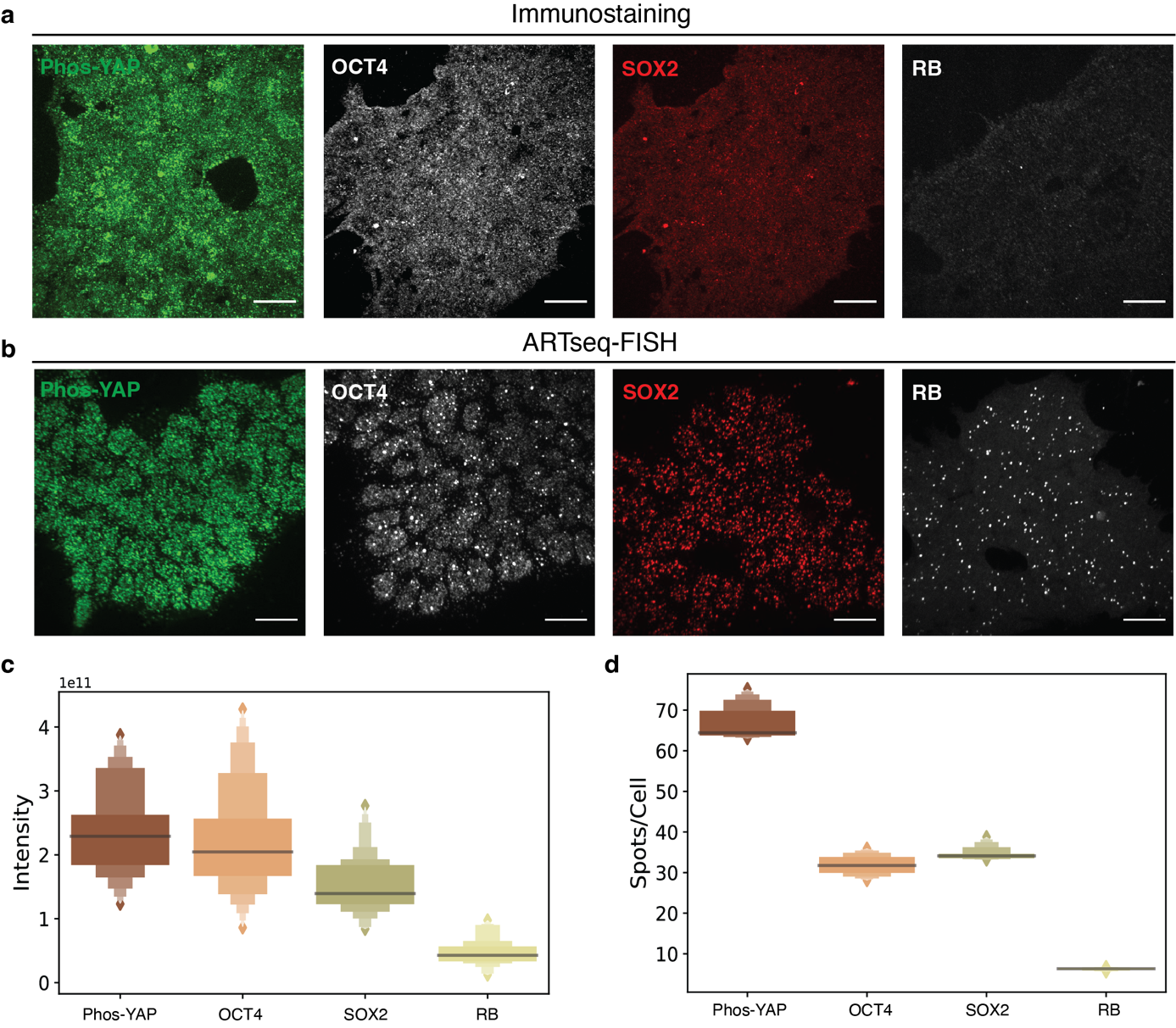

**Supplementary Fig. 19:** **a and b,** Representative max intensity and Z projection images of selected antibodies detected by immunostaining and ARTseq-FISH. Scale bar, 20 µm. **c,** Measuring cell fluorescence in immunostaining images of selected antibodies (Phos-YAP, OCT4, SOX2 and RB) with corrected total cell fluorescence (CTCF). CTCF = Total cell intensity - (area of selected cell * mean background intensity). The selected cells of each antibody were from three different images. And 10 cells are selected from each image. The mean background intensity is the average fluorescence reads of 10 areas in these three images. Line in the boxenplot denoting the median. **d,** Quantification of spots per cell of selected antibodies (Phos-YAP, OCT4, SOX2 and RB) applied in Ab-FISH. The counting results of each antibody match the trends of fluorescence intensity of each antibody shown in panel. Line in the boxenplot denoting the median.

Supplementary Fig. 20: Comparison between ARTseq-FISH with conventional smFISH.

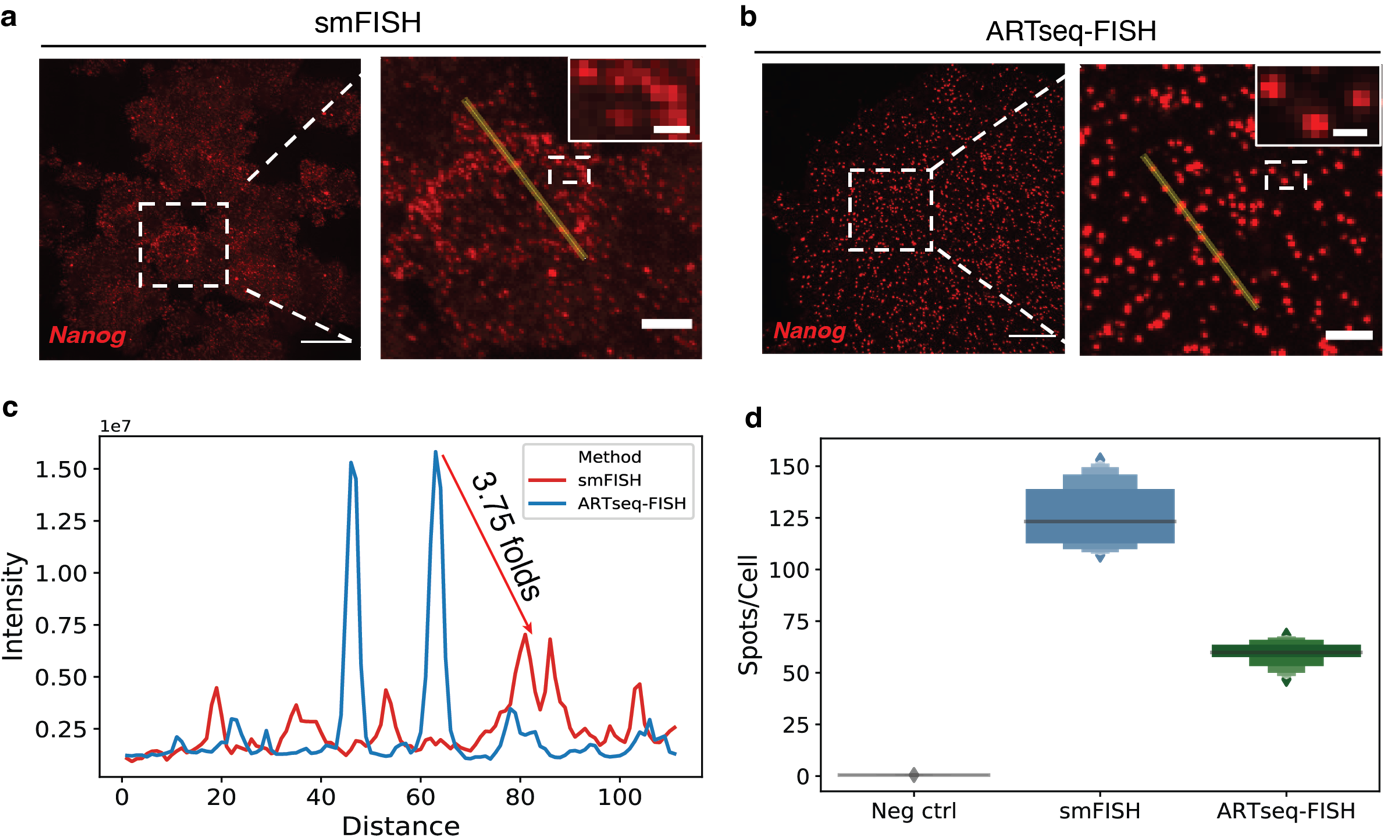

**Supplementary Fig. 20:** **a-d** *Nanog* mRNA detected by RNA RCA-FISH and conventional smFISH. **a and b** Representative max intensity and Z projection images show mRNA spots from these two methods. Zoomed images show the representative spots. Scale bar, 20 µm, 5 µm and 1 µm. **c,** Line plot illustrated the grey value of selected spots in zoomed images in panel **a and b**. The signal to noise ratio in RNA RCA-FISH is much higher than in smFISH. Arrow shows the representative differences of the signal to noise ratio in these two methods. **d,** Quantification of spots per cell of *Nanog* mRNA by smFISH and RNA RCA-FISH. (Neg ctrl n=241, smFISH n=203, RNA RCA-FISH n=332). Line in the boxenplot denoting the median.

Supplementary Fig. 21: Reproducibility of ARTseq-FISH.

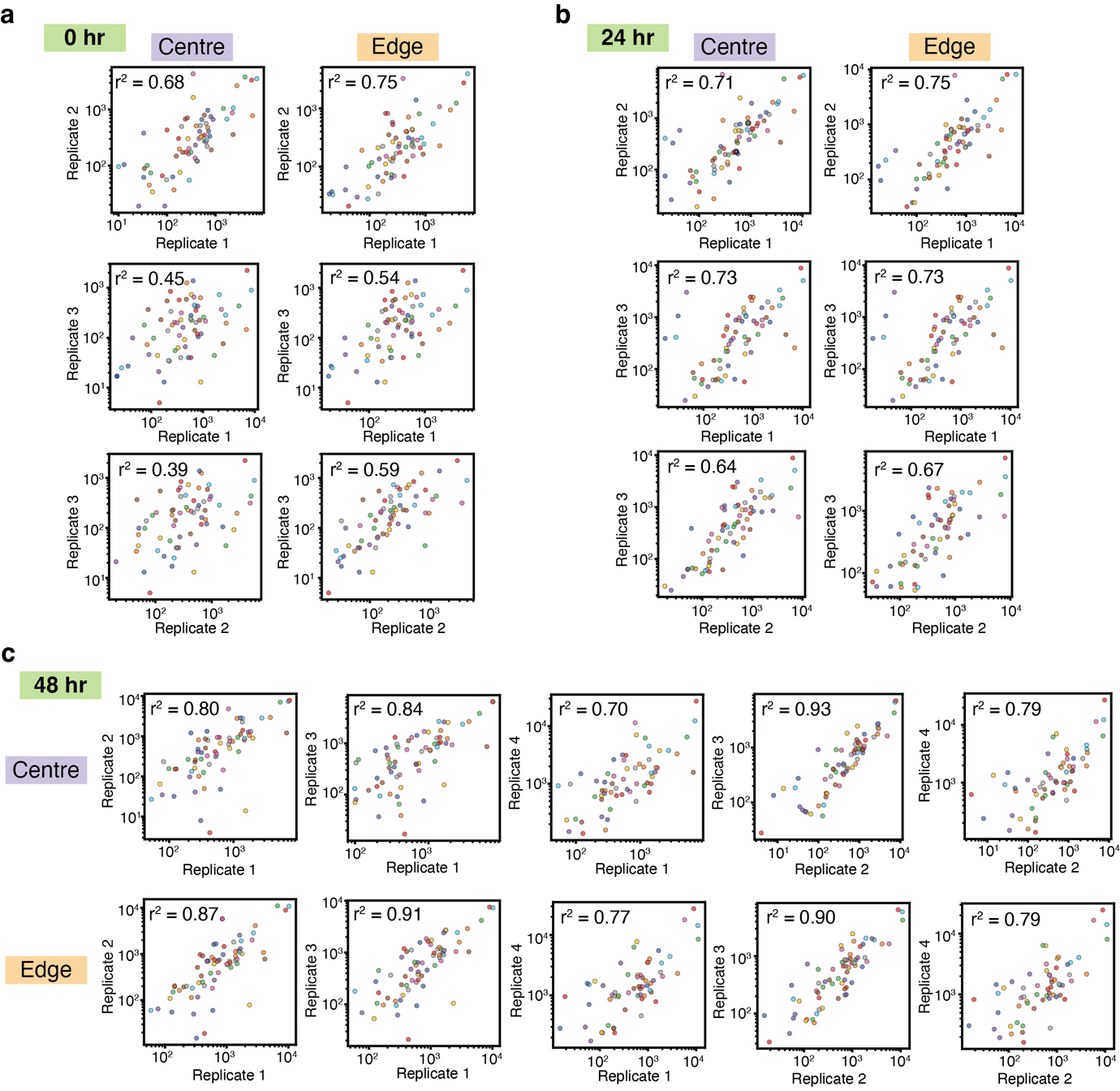

**Supplementary Fig. 21:** The correlation between different replicates of micropatterned mESCs with different positions (Centre and Edge) on the micropatterns. **a,** Three replicates of micropatterned mESCs from 0 hr after LIF withdraw. **b,** Three replicates of micropatterned mESCs from 24 hr after LIF withdraw. **c,** Four replicates of micropatterned mESCs from 48 hr after LIF withdraw.

Supplementary Fig. 22: Heterogeneity of serum cultured mESCs.

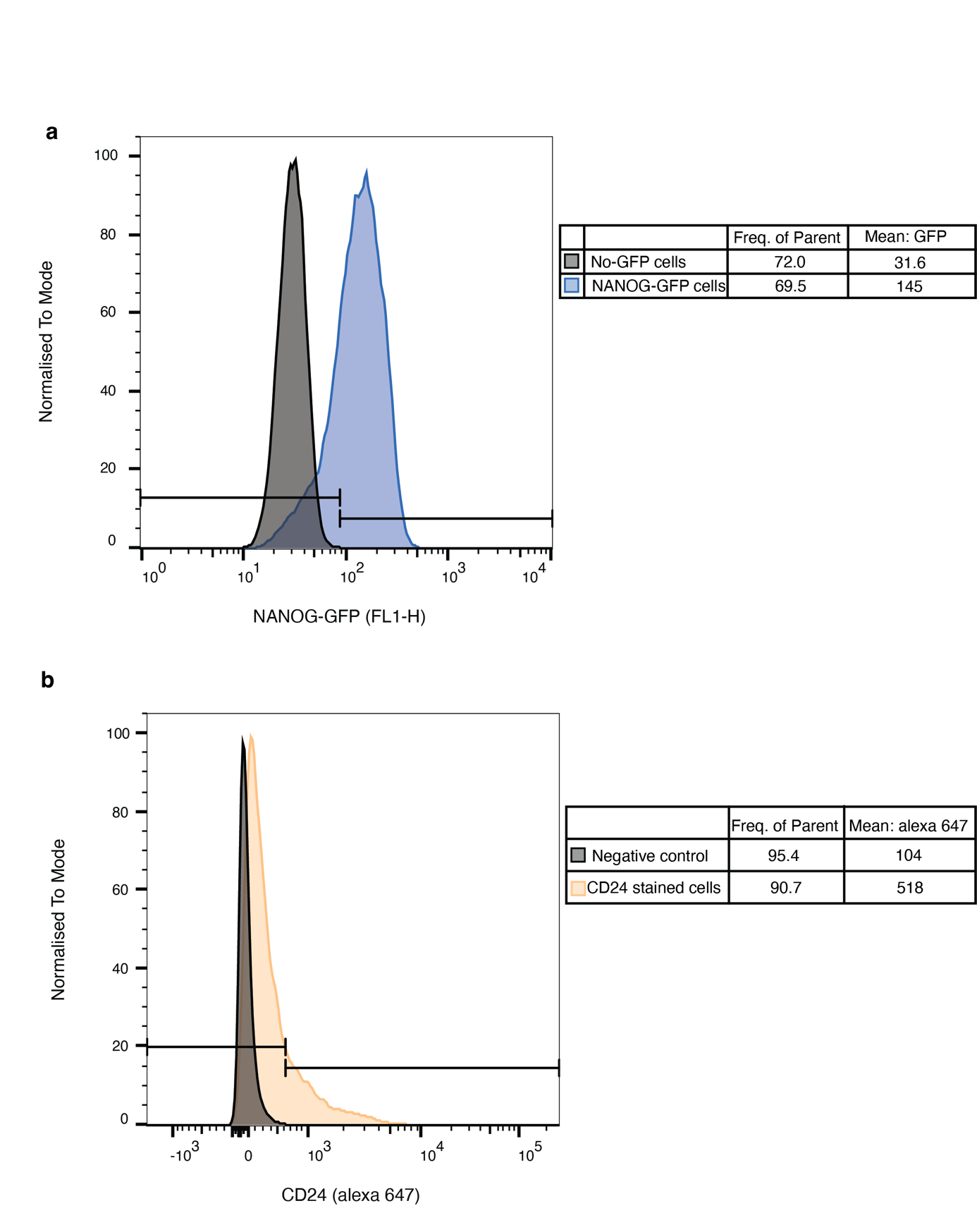

**Supplementary Fig. 22:** Serum cultured mESCs demonstrated different expression level of pluripotency marker (NANOG) and differentiation marker (CD24). **a,** Histogram of NANOG-GFP mESCs and negative control mESCs distribution in GFP channel. **b,** Histogram of CD24 stained mESCs and negative control mESCs distribution in Alexa647 channel.

Supplementary Fig. 23: Normalisation of the immunofluorescent images from the micropatterns.

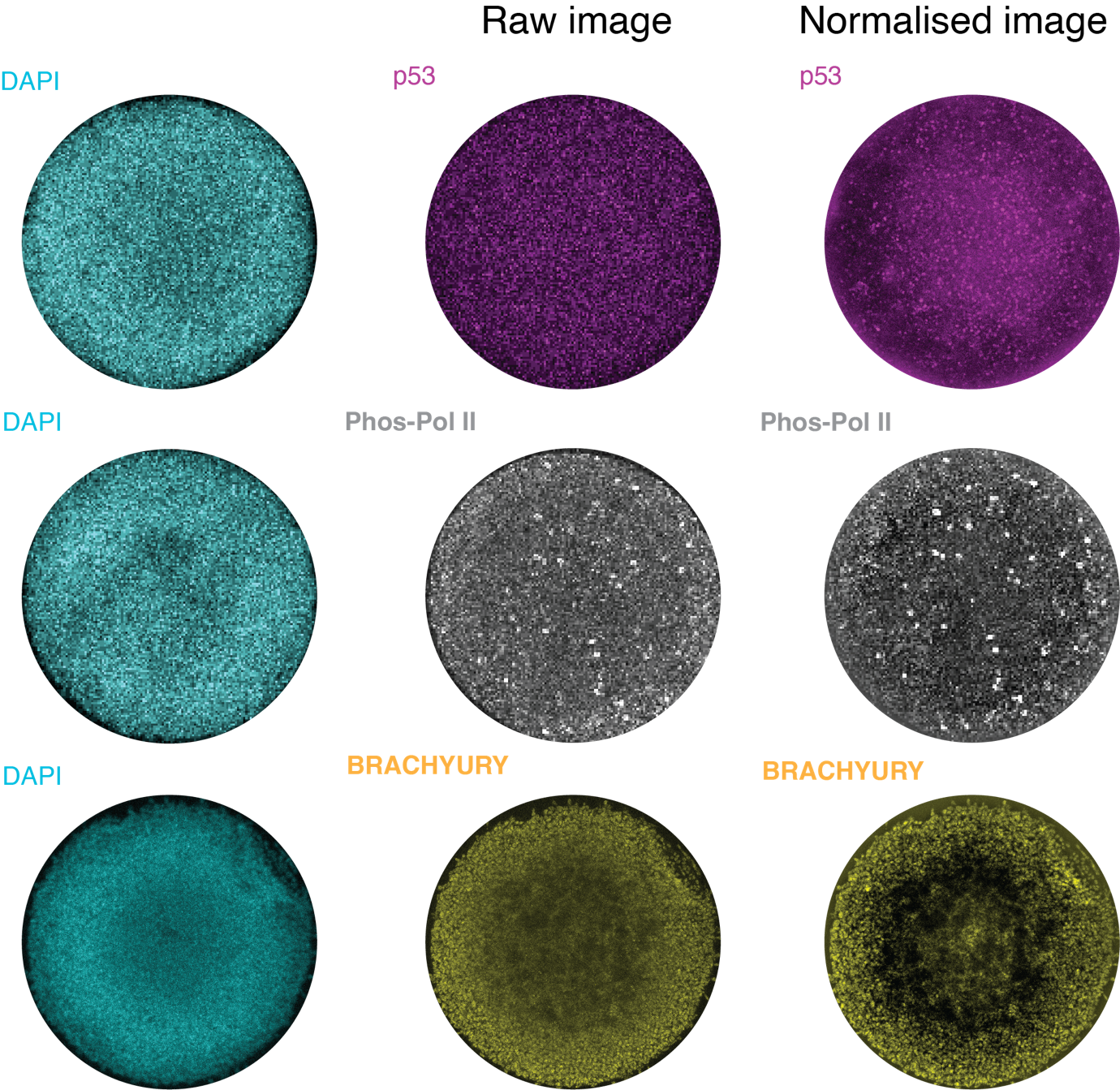

**Supplementary Fig. 23:** Images of immunostaining with p53, Phos-pol II and BRACHYURY antibodies on the micropatterned mESCs. Immuno-stained Images were normalised by the DAPI fluorescence intensity at each z-stack. Each individual image was normalised by their own mean, the DAPI image is smoothed using a gaussian mean filter (mask size = 101x101, original image size = 1024x1024). The DAPI is subsequently subtracted from the immuno-stained image. The filtered images were subsequently shown including the full z projection.

Supplementary Fig. 24: Mean fold changes of the targets between localized towards the edge and centre of the micropattern.

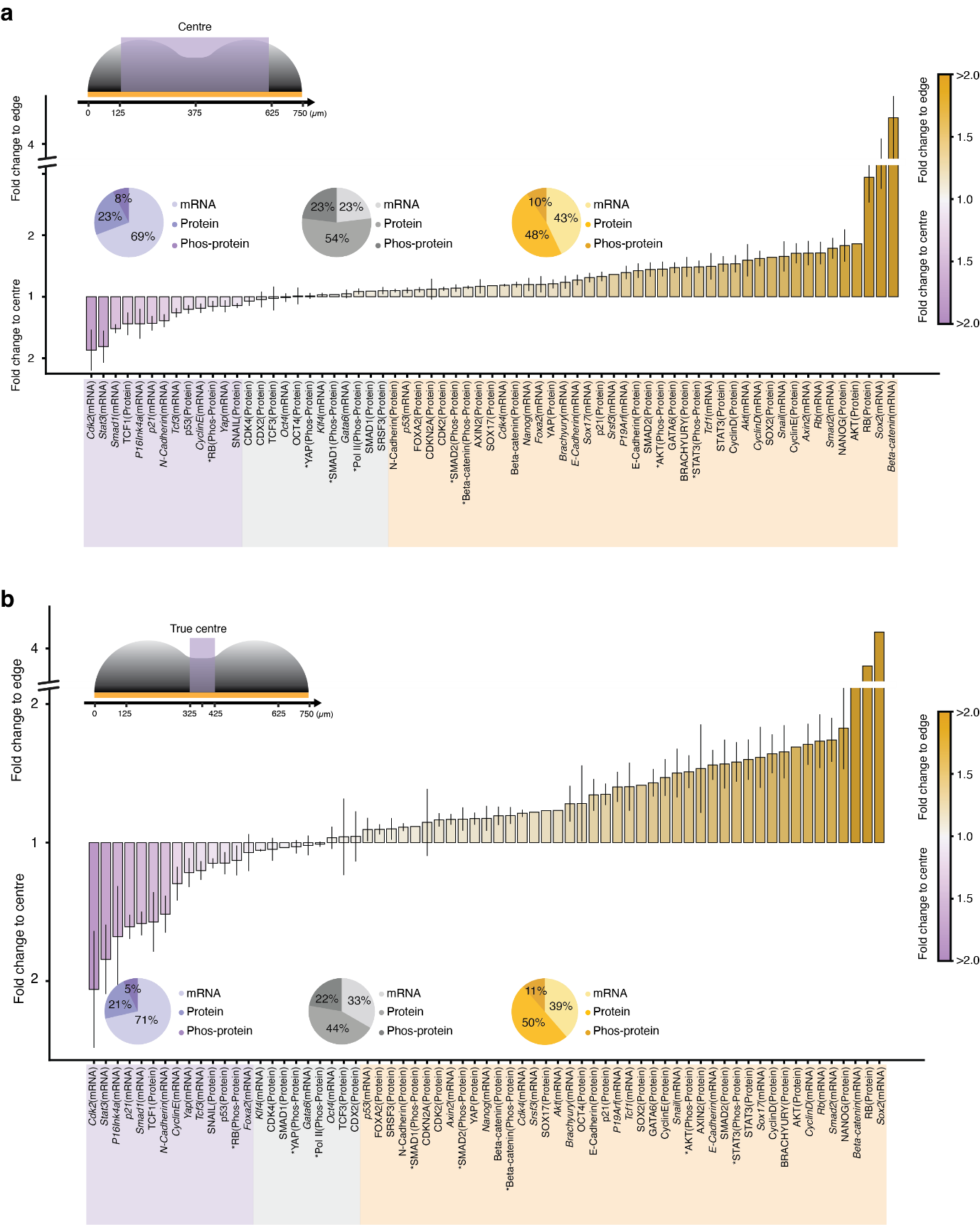

**Supplementary Fig. 24:** Mean fold change of the expression levels of the targets in cells at the edge or the centre of the micropattern at 48 hours after LIF withdrawal across the 4 biological replicates. In panel **a**, the centre represents the region exclusive of the edge. In panel **b**, the centre represents the region at the centre of the micropattern. Error bars represents SEM of 4 biological replicates.

Supplementary Fig. 25: Expression of selected targets across the micropattern and correlation.

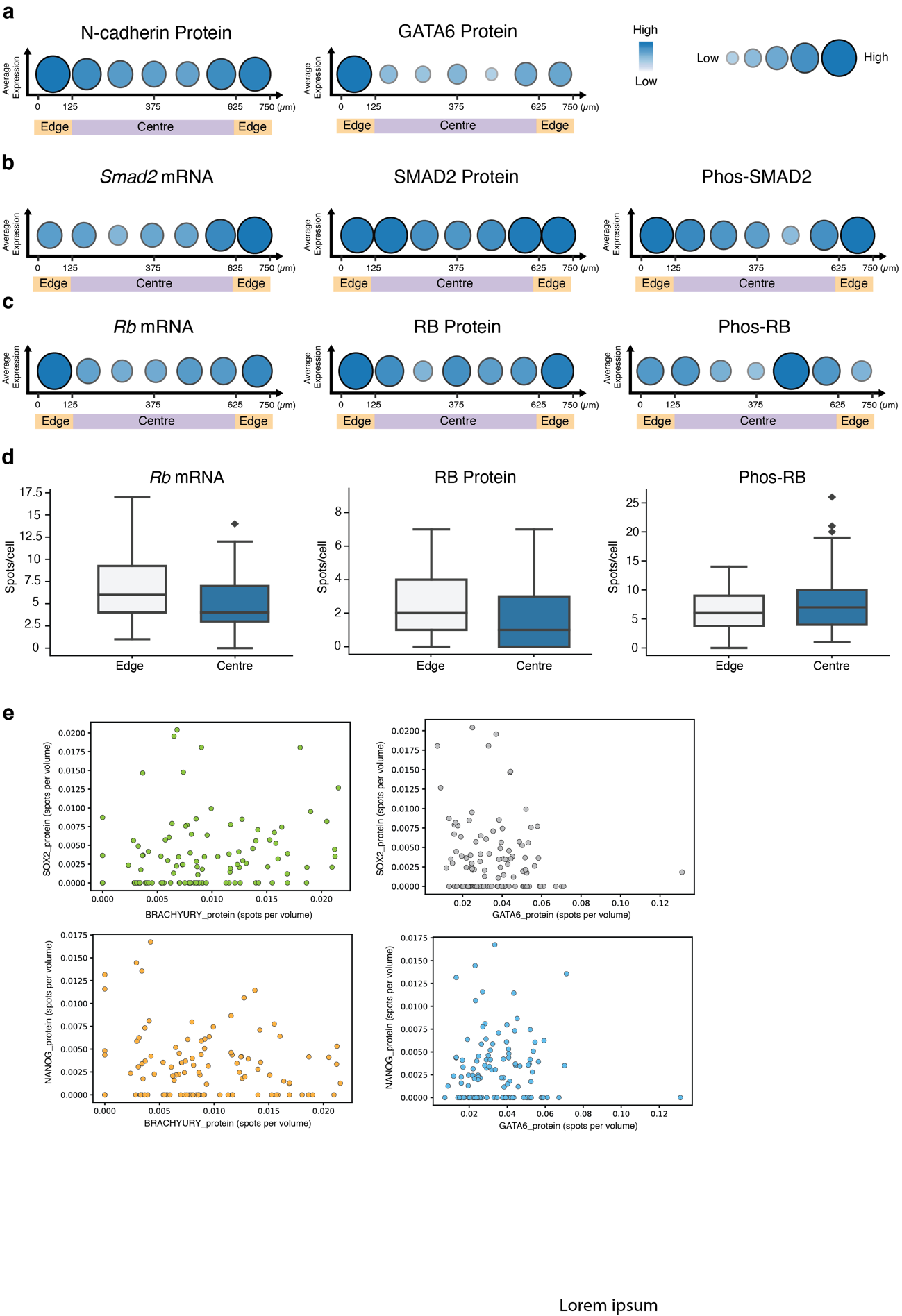

**Supplementary Fig. 25:** **a-c,** Average expression profile of spots per cell at seven positions across the micropattern for N-cadherin protein and GATA6 protein **(a)**, *Smad2* mRNA, protein and phosphoprotein **(b)**, *Rb* mRNA, protein and phosphoprotein **(c),** colour shaded area below x-axis illustrates the defined "edge" and "centre" of the micropattern cross-section. **d,** Quantification of spots per cell of *Rb* mRNA, protein and phosphorylated protein at the edge and centre of the micropattern. Line in the boxplot denoting the median. **e,** Correlation between SOX2 and BRACHYURY, SOX2 and GATA6, NANOG and BRACHYURY, NANOG and GATA6.

Supplementary Fig. 26: OCT4 protein detected by IF and ARTseq-FISH.

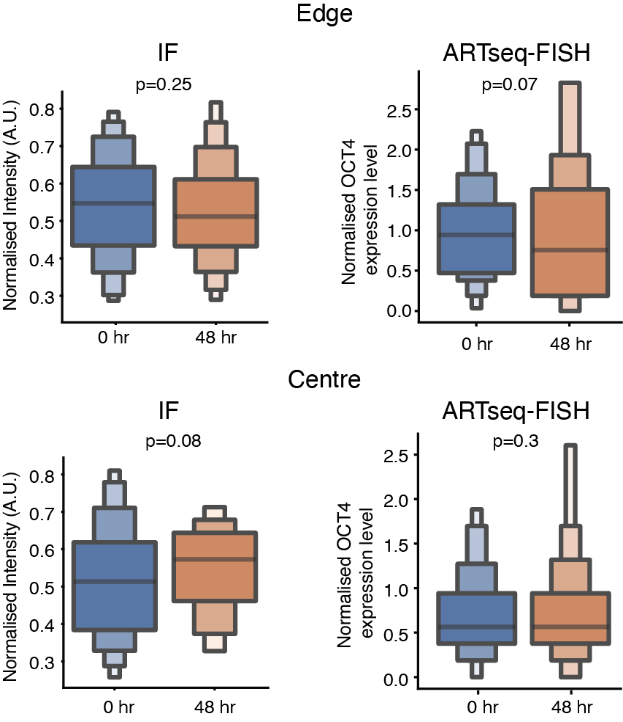

**Supplementary Fig. 26:** Boxenplots shows the normalised intensity of OCT4 detected by immunofluorescence at the edge (top left) and centre (bottom left) of the micropattern at 0 hr and 48 hours post LIF withdrawal. Boxenplots shows the normalised expression of OCT4 protein detected by ARTseqFISH at the edge (top right) and centre (bottom right) of the micropattern at 0 hours and 48 hours post LIF withdrawal. Line in the boxenplot denoting the median.

Supplementary Fig. 27: Ki-67 staining on the micropatterned mESCs overtime after LIF withdrawal.

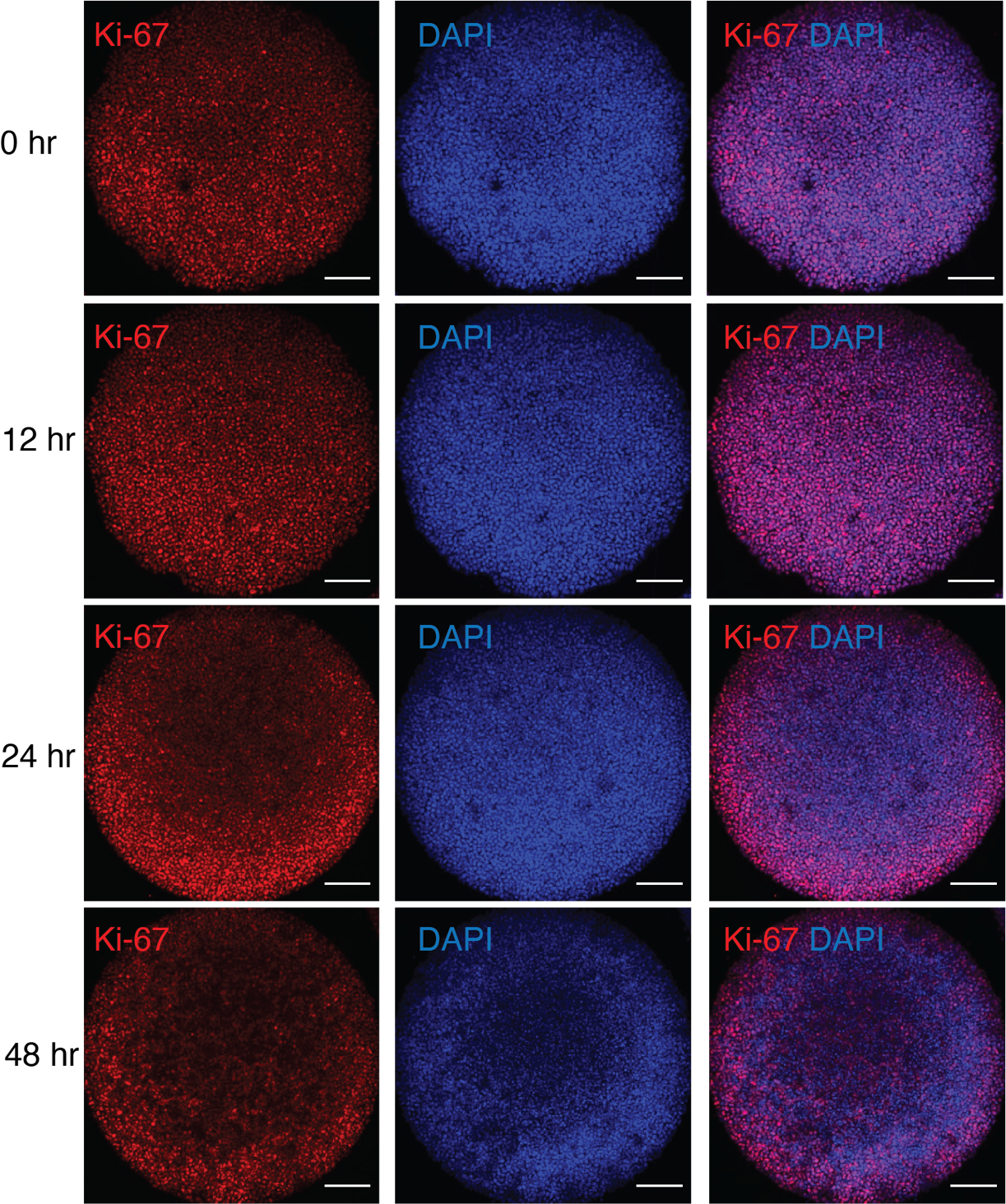

**Supplementary Fig. 27:** Representative images showing Ki-67 staining of the micropatterned mESCs at 0, 12, 24 and 48 hours after LIF withdrawal. Images from different time points are set as the same maximum and minimum grey value. Scale bar, 120 µm.

Supplementary Fig. 28: p53 expression profile and apoptosis in the micropatterned mESCs.

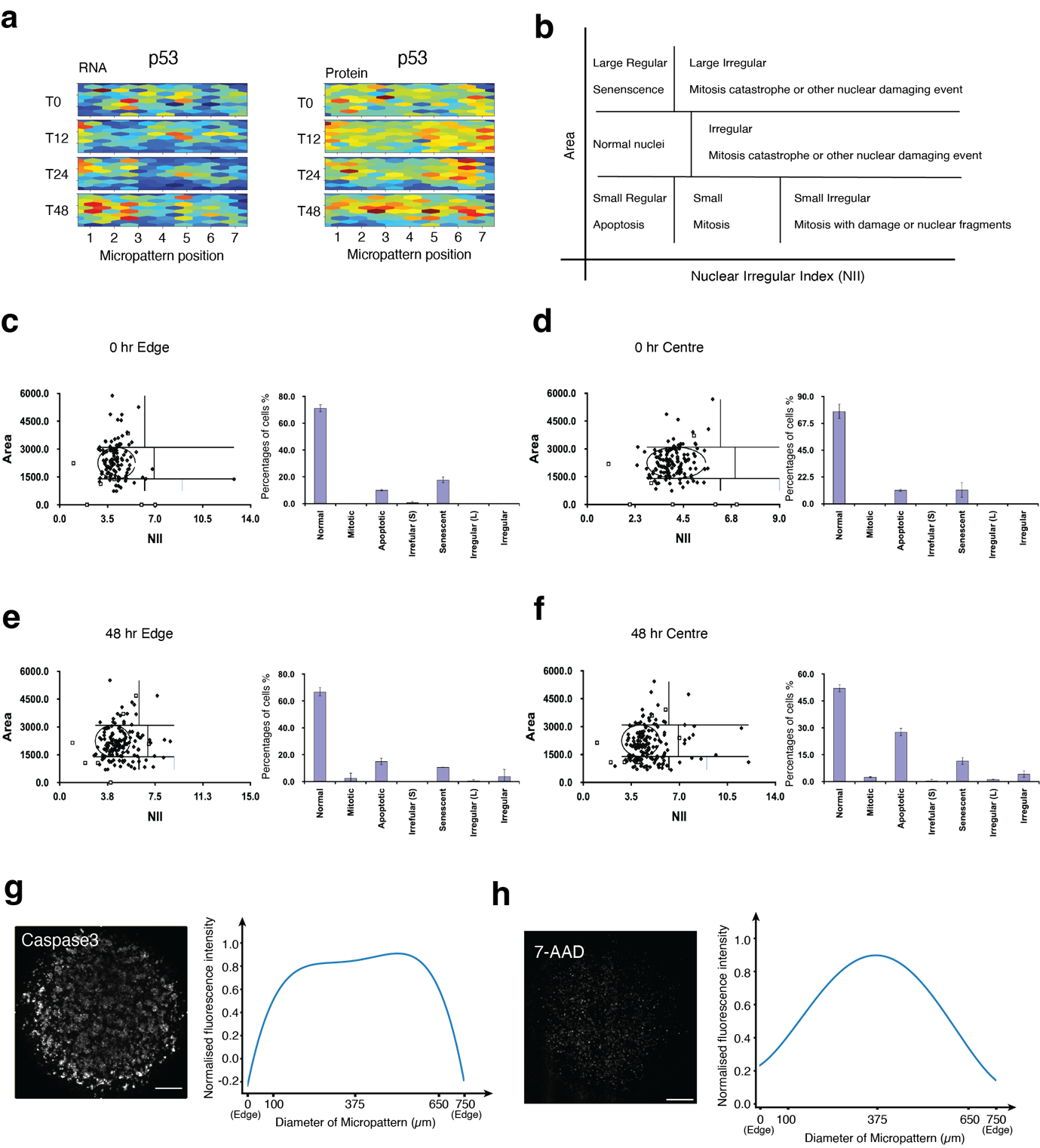

**Supplementary Fig. 28:** **a,** Heatmap of p53 mRNA and protein expression profile across the micropattern at 0, 12, 24 and 48 hours after LIF withdrawal. **b**, Schematic shows the distribution of nuclei in a plot of area versus nuclear irregularity index (NII). **c-f**, The distribution of nuclei and the percentage of cells in different cellular stages at 0 and 48 hours after LIF withdrawal at the edge and centre of the micropattern. Error bars represent the standard deviation across >100 nuclei per condition. **g-h**, Representative images of Caspase 3 and 7-AAD staining on micropatterned mESCs at 48 hours after LIF withdrawal (left). Scale bar, 120 µm. Quantification of relative fluorescent intensity of Caspase 3 and 7-AAD staining in mESCs across the micropattern at 48 hours after LIF withdrawal.

Supplementary Fig. 29: Cell cycle phase in the micropatterned mESCs over time after LIF withdrawal.

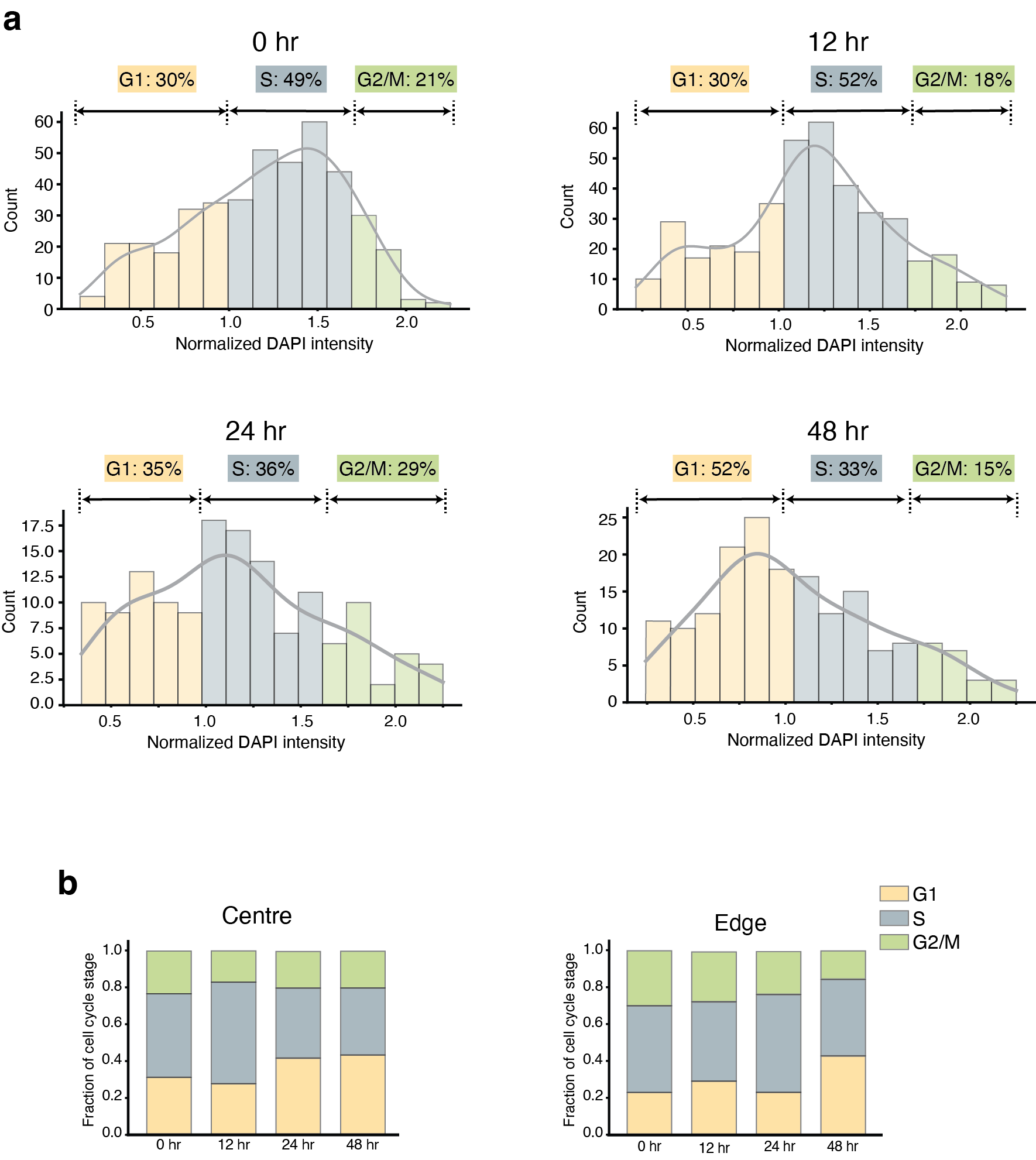

**Supplementary Fig. 29: a,** Histogram of cell cycle phases of cells at 0 hours, 12 hours, 24 hours, and 48 hours after LIF withdrawal. **b,** Fraction of cell cycle stages in cells located at the centre and the edge of the micropattern at 0 hours, 12 hours, 24 hours, and 48 hours after LIF withdrawal.

Supplementary Fig. 30: Cell cycle stages of mESCs quantified by flow cytometry.

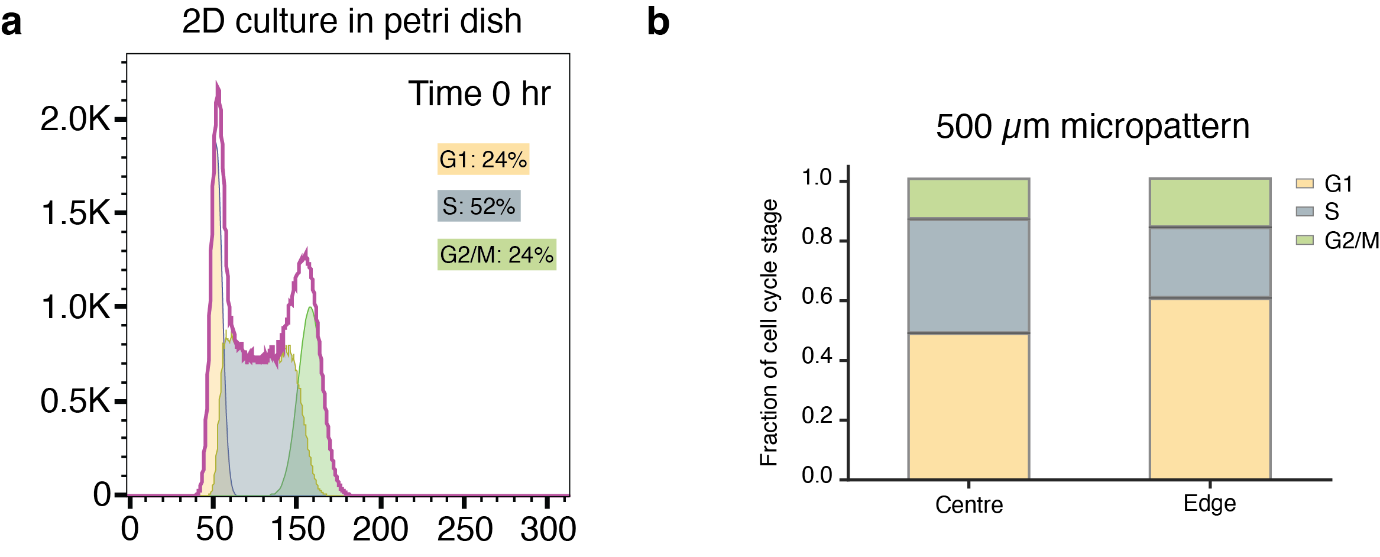

**Supplementary Fig. 30:** **a,** Flow cytometry results showing the cell cycle distribution of the mESCs cultured on 2D petri dish at 0 hours LIF withdraw.

Supplementary Fig. 31: Single cell tracking of Fucci mESCs and NANOG-GFP mESCs.

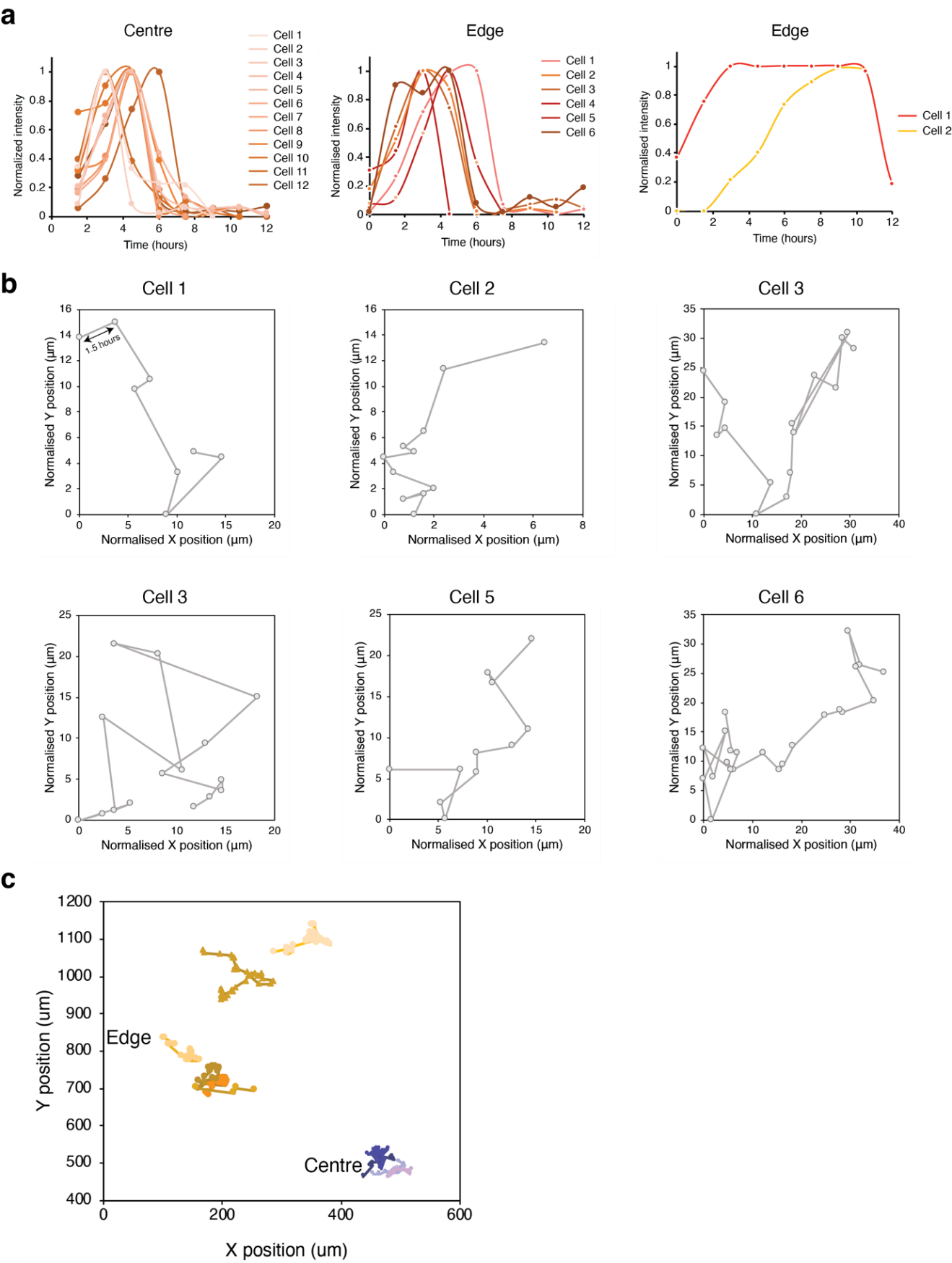

**Supplementary Fig. 31:** **a,** Single cell KO-2 (G1) traces of tracked cells that enter G1 within the first 5 hours of LIF withdrawal at the centre and edge of the micropattern. **b,** Normalised X-Y displacement of 6 individual NANOG-GFP mESCs. Each point is 1.5 hours apart. **c,** X-Y displacement of 8 individual Fucci mESCs at the centre (orange) and edge (purple) of the micropattern after LIF withdrawal. Each trajectory is one cell, and each point is 1.5 hours apart.

### Supplementary Tables Caption

Supplementary Table 1 Probe sequences in ARTseq-FISH.

Probes and oligonucleotides used for detecting mRNAs and proteins in ARTseq-FISH.

Supplementary Table 2 Antibody list.

The information of the antibodies used in this study, including names, cat numbers, suppliers, detecting total protein or non-phosphorylated protein.

Supplementary Table 3 *Nanog* smFISH probes.

Probe sequences of the *Nanog* mRNA detection by smFISH.

Supplementary Table 4 Barcode.

The barcode used to decode the targets after sequential hybridization. Each target is decoded by the readout colour in particular hybridization round.

Supplementary Table 5 Categories of targets.

The categories of all the targets, and the targets used in the heatmaps in Figure 5.
